# A common garden of *Halichondria* sponges: taxonomic revision of Northeast Pacific Halichondriidae reveals many cryptic introduced species

**DOI:** 10.1101/2024.11.04.621922

**Authors:** Thomas L. Turner, Christine Morrow, Bernard Picton, Claire Goodwin, Robert W. Thacker

## Abstract

Sponges (phylum Porifera) possess biochemical, cellular, and physiological traits with valuable biotechnical applications. However, our ability to harness these natural innovations is limited by a classification system that does not fully reflect their evolutionary history. In this study, we uncover numerous cryptic species within the genus *Halichondria* that are morphologically indistinguishable from the well-known *Ha. panicea*. Many of these species have habitat preferences and geographic distributions that strongly suggest they have been dispersed by human activity. Most of these species are broadly sympatric with their closest relatives, and these overlapping distributions allow us to use patterns of DNA variation to infer reproductive isolation between clades in nature. With reproductively isolated species thus delineated, we can use DNA states as taxonomic characters to formally describe them. Though much remains to be learned about these newly discovered species, the natural "common gardens" of these sponges in California, New York, and other locations provide opportunities to test hypotheses about their diversification in future work.

## Introduction

The common ancestor of extant sponges is estimated to have lived over 700 million years ago (Dohrmann and Wörheide, 2017; Pankey et al., 2022; Rossi et al., 2024). This ancient origin suggests that some sponge lineages have been independently evolving for longer than the most distantly related animals within Bilateria. Sponges have undoubtedly diversified in biochemical, cellular and physiological dimensions during this prolonged time, developing numerous innovations with potential applications (Dunn et al., 2015; Mehbub et al., 2024). These innovations remain largely undiscovered, and it seems likely that a major impediment to advancing sponge biology is the lack of a stable classification system that reflects the evolutionary history of the phylum. The difficulty in assigning sponge specimens to species and determining where those species fall in the tree of life creates uncertainty that deters investigation across multiple disciplines.

Creating an accurate taxonomy of sponges based on morphological characters has proven challenging due to a paucity of informative character states. Sponge taxonomy is based mainly on skeletal characters: spicule morphology and the arrangement of spicules and/or spongin fibers into a three-dimensional skeleton (Hooper and Van Soest, 2002). While these features have been useful for diagnosing many species and higher taxonomic groups (Hooper and Van Soest, 2002; Morrow and Cárdenas, 2015), molecular phylogenies have revealed that skeletal traits alone are inadequate due to both insufficient differences between closely related species and convergently evolving traits in distantly related species (de Paula et al., 2012; Erwin and Thacker, 2007; Gazave et al., 2010; Morrow et al., 2013; Turner and Pankey, 2023; Xavier et al., 2010). These limitations are frequently encountered in sponge families with simple spicules and a loosely organized skeleton, such as the Halichondriidae. This family is of significant interest due to its pharmaceutical potential (Hirata and Uemura, 1986; Rodriguez Jimenez et al., 2021; Zhong et al., 2024) and as a model system, as this family includes one of the most studied sponges in the world, *Halichondria panicea* (Pallas, 1766). This species is found near many marine labs, is amenable to laboratory culture, and as a result, has been used to ask questions about sponge physiology, symbiosis, ecology, immunology, and more (Barthel, 1986; Barthel and Detmer, 1990; Barthel and Wolfrath, 1989; Carrier et al., 2023; Forester, 1979; Knobloch et al., 2019; Schmittmann et al., 2021; Thomassen and Riisgård, 1995; Witte et al., 1994). However, this research is built atop a shaky foundation, because the taxonomy and systematics of *Ha. panicea* and its relatives is very poorly delineated. The World Register of Marine Species lists over 70 previously named species that are now considered synonyms of *Ha. panicea* (de Voogd, et al., 2024). Other species that remain putatively valid, like *Ha. bowerbanki* (Burton, 1930), are often difficult to differentiate from *Ha. panicea* and their validity remains in dispute. Cryptic species within populations ascribed to *Ha. panicea* have remained a possibility, as this species has supposedly been found in most of the world’s oceans and in a wide variety of climates. Under what can be called a "cryptic endemic species in different regions" hypothesis, different regions of the world could contain morphologically indistinguishable species that are endemic to each region (Erpenbeck, 2004). An alternative hypothesis, which can be called the "single globally-distributed invasive species" hypothesis, is that *Ha. panicea* has been unintentionally introduced by human activity to many regions of the world, as was recently shown for the co-familial species *Hymeniacidon perlevis* (Samaai et al., 2022; Turner, 2020).

Below, we test these hypotheses by integrating extensive field collections, genomic analyses, and traditional morphological characters. For the genomic analyses, we use a combination of Illumina and Sanger sequencing to assemble mitochondrial genomes and nuclear ribosomal loci. Because the mitochondrial and nuclear genomes are unlinked, reciprocal monophyly across these compartments is strong evidence of reproductive isolation when species are sympatric (Jennings, 1917; Rannala and Yang, 2020). In cases where we find strong evidence of reproductive isolation, we formally describe these new species. Formally describing newly discovered species is important because unique identifiers (such as Linnean names) are crucial for making species visible to the research community and linking data across databases like GenBank, the World Register of Marine Species, the Global Biodiversity Information Facility, and museum inventories (Delić et al., 2017). Formally describing new species requires finding unique characters that differentiate them from previously described species. When morphological characters cannot be found, DNA states can provide characters to define and diagnose species (Delić et al., 2017; Eitel et al., 2018; Jörger and Schrödl, 2013; Lawley et al., 2021; Tessler et al., 2022). The use of DNA sequences to formally describe new species has a complicated history, with some recent works claiming that the publication of a consensus DNA "barcode" (a few hundred basepairs of the *cox1* locus) is sufficient to describe a new species by itself (Sharkey et al., 2021). This latter approach has been heavily criticized, but these criticisms often explicitly note that the use of DNA characters to describe morphologically cryptic species is valid when done properly (Ahrens et al., 2021; Meier et al., 2022; Zamani et al., 2022). For the species described herein, we were therefore careful to avoid the pitfalls of these earlier works, and instead follow the example of authors (e.g., Eitel et al., 2018; Lawley et al., 2021) who used DNA-based descriptions that were consistent with the International Code of Zoological Nomenclature (ICZN, 1999). It remains true that using DNA characters as taxonomic states has drawbacks: for example, the identification of unknown samples will be limited by preservation methods, sample age, and the expertise of investigators. However, these problems would likely be even greater for alternative phenotypic traits such as proteomics or histology, which would require less ubiquitous expertise and less common preservation methods. We hope that our approach will lay the groundwork for future research into the phenotypic differences among these newly described species.

## Methods

### Collections

Sponges were collected from the Atlantic Coast of Canada (13 samples, Claire Goodwin), Northern Ireland (10 samples, Christine Morrow and Bernard Picton), Sweden (9 samples, Raquel Pereira), Portugal (2 samples, Raquel Pereira), Panama (6 samples, Robert Thacker) and three locations in the USA: New York (14 samples, Robert Thacker), Washington (19 samples, Thomas Turner, Robert Thacker, and Brooke Weigel), and California (165 samples, Thomas Turner). Collections were most extensive in California, where 95 subtidal sites, 14 intertidal sites, and 23 marinas were surveyed. Subtidal sites were surveyed by divers, with no new collections from deep water (maximum depth 32 m). Previously vouchered material was acquired on loan from the Royal British Columbia Museum (voucher numbers with RBC), the Florida Museum (voucher numbers with UF or BULA), the Smithsonian Institution’s National Museum of Natural History (voucher numbers with USNM), the Naturalis Biodiversity Center (voucher numbers with ZMA), and the California Academy of Sciences (voucher numbers with CASIZ). Freshly collected material was deposited in these collections and also the Santa Barbara Museum of Natural history (voucher numbers with SBMNH), the Cheadle Center at the University of California, Santa Barbara (voucher numbers with UCSB), the Atlantic Reference Centre Museum (voucher numbers with ARC), the Ulster Museum (voucher numbers with BELEM), the Yale Peabody Museum (voucher numbers with YPM), and the National History Museum of Los Angeles (voucher numbers with NHMLA). A table listing all examined samples with voucher numbers, GenBank accessions, collection dates and locations, and other metadata is available as supplementary data via Data Dryad (https://doi.org/10.5061/dryad.bvq83bkk6).

### Morphology

Spicules were examined after digesting sponge subsamples in bleach. For some samples, a single piece of sponge, including both the ectosome and choanosome, was used for digestion. When sizes of spicules were compared between ectosome and choanosome, these domains were carefully separated with a razor and digested individually. Spicule measurements were taken from images using ImageJ (Schneider et al., 2012), with pixel-to-mm conversion determined using a calibration slide. The length of each spicule was measured as the longest straight line from tip to tip, even if the spicules were curved or bent. Width was measured at the widest point, excluding any adornments like swollen tyles. All spicule measurements are available as supplementary data via Data Dryad.

To analyze the choanosomal skeletons, sponge fragments were first frozen into ice cubes and then hand-cut into tissue sections. These sections were cleared using one of two methods. A few samples were transferred from a 95% ethanol bath to a 100% ethanol bath, and then cleared with histoclear (National Diagnostics). Most samples were digested in a mixture of 97% Nuclei Lysis Solution (Promega; from the Wizard Genomic DNA Purification Kit) and 3% 20 mg/ml Proteinase K (Promega). This method removed cellular material while preserving spicules and the spongin network (if present). For the ectosomal skeletons, the sponge surface was removed with a razor. In many cases, these skeletons could be imaged directly without further processing; if opaque, they were processed similarly to choanosomal sections.

To measure ectosomal tract thickness, we averaged the measurements of the three thickest visible tracts, excluding large matted areas. This process was difficult to standardize and the measurements are likely less reproducible than spicule size measurements. We also categorized ectosomal skeletons based on their morphology, noting that there were potentially distinct types. The five categories used were: 1. Multilayered and gothic-crypt-like; 2. Sieve-like (a dense mat with approximately round open spaces); 3. Lace-like (a fairly isotropic mesh); 4.

Intermediate forms between categories 3 and 5; 5. A network of spicule tracts (resembling either straight lines or meandering lines, similar to dry or wet spaghetti). In some cases, ectosomal skeletons were found as dense mats of spicules without open spaces, but in these cases, other areas of the same sponge exhibited one of the above types.

### Illumina sequencing

DNA was extracted with several different kits including the Wizard Genomic DNA Purification kit (Promega) and the Qiagen Blood & Tissue kit. Prior to Illumina sequencing, samples were treated with RNase A and repurified with the DNA Clean and Concentrator (Zymo). Illumina sequencing and library preparation were performed at the DNA Technologies Core Facility at the University of California, Davis. Libraries were made with the Super-High-Throughput (SHT) technique, which involved dual indexing each sample and pooling 96 samples before sequencing. Paired-end 150 bp reads were generated using the Illumina NovaSeq instrument, resulting in a median of 2.1 million reads per sample. The number of reads varied significantly across samples, ranging from a maximum of 17 million reads to a minimum of 208 reads. The reasons for this variability were not explored in detail, but DNA extractions appeared to be of variable quality even before library construction and pooling, perhaps because of sample preservation differences or the secondary chemistry of samples.

Reads were trimmed with fastP (Chen et al., 2018) and then de novo assembled into contigs with Megahit (Li et al., 2015). The number of contigs per sample ranged from 0 to 1,080; the median number of contigs for samples with at least one contig was 22. Blast searches revealed that many contigs were microbial (or occasionally from other taxa), but the sponge’s nuclear ribosomal locus was usually the highest coverage contig. Indeed, in samples where only one or two contigs assembled, the ribosomal locus was usually one of them. In samples where the full ribosomal locus assembled de novo, this contig was then used as a reference sequence, and reads from that sample were aligned to it with bwa-mem (Li and Durbin, 2009). SAMtools (Danecek et al., 2021) was then used to convert SAM files from this alignment into a consensus sequence, which served as the final sequence for that sample. For samples where no ribosomal contig assembled at the megahit step, the most closely related sequence available (either from another sample in our analysis or from GenBank) was used as reference sequence, and bwa-mem was used to align that sample’s reads to this reference. In some cases, ribosomal contigs assembled with megahit were incomplete or fragmented, so these partial contigs were spliced into the nearest available reference sequence to create a hybrid reference for bwa-mem alignment. This approach successfully yielded nearly complete sequences of the 18S, 28S, and intergenic transcribed spacer regions for most samples, though some had a significant proportion of uncalled bases ("N"). Many of these samples had been previously sequenced at the 28S locus with Sanger sequencing, and these sequences were combined with Illumina sequencing when available. Combining Sanger and Illumina data often resulted in resolving some uncalled bases but in no cases were Illumina consensus sequences found to have called bases that conflicted with Sanger data, suggesting error rates of called bases are low. Ribosomal sequences were assembled for 110 samples, with a median number of called bases (excluding Ns) of 6,429 bp (ranging from 2,227 to 6,589 bp). The number of called bases for each sample is listed in the supplementary table available from Data Dryad.

Mitochondrial contigs were present in some of the Megahit assemblies but were generally less complete than ribosomal loci. To generate more complete mitochondrial genomes, we aligned reads to existing reference sequences using bwa-mem. The references used included *Halichondria okadai* (MG267395), *Halichondria galea* sp. nov. (KX244759), *Halichondria dokdoensis* (MK105765), *Halichondria akesa* sp. nov. (MH756604), *Halichondria panicea* (MH756603), and *Hymeniacidon perlevis* (KF192342).

For species lacking closely related reference sequences (e.g., the Bowerbanki species group, as defined in figure 1, and *Hymeniacidon pierrei* sp. nov.), we created hybrid reference sequences. This procedure involved replacing parts of the closest available reference genome with de novo contigs from the megahit assembly. Although this method allowed us to assemble substantial portions of the mitochondrial genome for samples without good reference sequences, these assemblies were less complete, often containing more uncalled bases ("Ns") due to regions being too divergent for accurate short-read alignment. For instance, the average number of called bases in *Halichondria panicea* mitogenomes was 19,228, while for *Halichondria bowerbanki*, it was 16,100. In total, we assembled mitochondrial genomes for 75 samples, with a median of 16,888 called bases. As seen with ribosomal loci, combining these sequences with Sanger data at cox1 did not find errors in called bases. The number of called bases for each sample is listed in the supplementary table available from Data Dryad.

**Figure 1.**
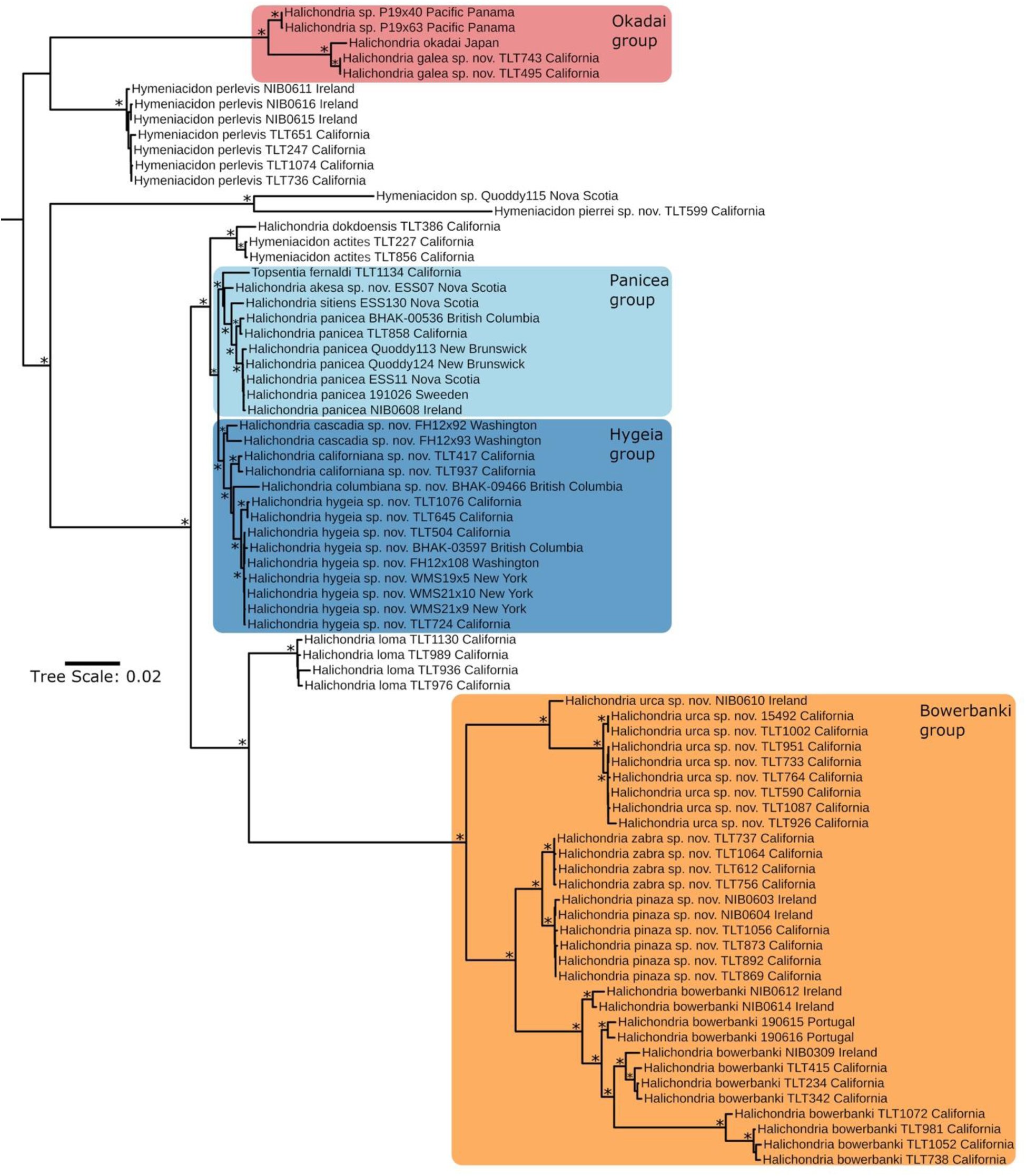
Maximum likelihood phylogeny for samples with assembled mitogenomes and nearly complete ribosomal loci. Asterisks at nodes indicate a UFbootstrap ≥ 99%. Scale bar indicates substitutions per site.

A sequence of the cox1 locus from the *Ha. panicea* neotype was generated using sequence capture techniques (Agne et al., 2022) and was generously shared with us by Dirk Erpenbeck.

### Sanger sequencing

The cox1 region commonly sequenced in sponges is the Folmer region, a 709 bp segment amplified using the primers LCO1490 and HCO2198 (Folmer et al., 1994). We used these primers for a few samples when other primers failed to amplify, though they are less preferred due to their shorter length and their tendency to co-amplify microbial sequences. We also attempted to amplify DNA using the primers LCO1490 and cox1-R1, and were successful with some samples (Rot et al., 2006). These primers target a larger 1264 bp region that includes both the Folmer region and the "co1-ext" region, which has been useful for distinguishing sponge species (Erpenbeck et al., 2006). However, we found that *Halichondria loma* and the Bowerbanki species group (Figure 1) had mismatches with the LCO1490 primer, and amplifications usually failed with both primer sets. To improve amplification success, we designed new primers, halcox1-L and halcox1-R, which were effective for all tested *Halichondria* species. These primers amplify an 886 bp region that encompasses most of the Folmer region and part of the co1-ext region. We used these new primers for all *Halichondria* samples not already processed with Illumina or other primer sets. Primer sequences are provided in Table S2.

We sequenced two spans of the 28S nuclear locus. The D1-D2 region, which spans 940 bp and is highly variable, was amplified in all samples using primers 15F and 878R (Morrow et al., 2012). We also amplified the adjacent D3-D5 region in some samples using primers 830F and 1520R. This primer pair amplifies a 710 bp fragment that overlaps the D1-D2 fragment by 20 bp (Morrow et al., 2012).

The ND1 locus was amplified in selected taxa to verify that fixed differences detected in Illumina sequencing were fixed in a larger sample. Primers were newly designed, but universal primers for all target species were not found. For the *Ha. dokdoensis*, *Ha. actities*, the Panicea species group (Figure 1), and Hygeia species group except for *Ha. columbiana* sp. nov., we used primers nd1-pan-R and nd1-pan-L. For *Ha. columbiana* sp. nov., nd1-pan-R was used with nd1- hygc-L. For the Bowerbanki species group, nd1-bow-R and nd1-bow-L were used. For *Ha. loma*, nd1-loma-R was used with nd1-bow-L.

PCR was performed using a Biorad thermocycler (T100) using the following conditions: 95°C for 3 min, followed by 35 cycles of 94°C for 30 sec, [annealing temperature] for 30 sec, 72°C for [extension time], followed by 72°C for 5 minutes. The annealing temperatures and extension times for each primer pair were as follows: halcox1 (54°C annealing, 60 second extension), LCO & R1 primers (50°C annealing, 75 second extension), LCO & HCO primers (52°C annealing, 60 second extension), 28S region (53°C annealing, 60 second extension for both primer sets), ND1 locus (54°C annealing, 60 second extension, all primer sets). PCR was performed in 50 μl reactions using the following recipe: 24 μl nuclease-free water, 10 μl 5x PCR buffer (Gotaq flexi, Promega), 8 μl MgCl, 1 μl 10 mM dNTPs (Promega), 2.5 μl of each primer at 10 μM, 0.75 bovine serum albumin (10 mg/ml, final concentration 0.15 mg/ml), 0.25 μl Taq (Gotaq flexi, Promega), 1 μl template. Post-PCR, samples were cleaned using ExoSAP-IT (Applied Biosystems) and then sequenced by Functional Biosciences with Big Dye v3.1 on ABI 3730xl instruments. We verified that the resulting sequences were of sponge origin using BLAST. All sequences have been deposited in GenBank under the accession numbers listed in the supplementary table available from Data Dryad.

### Phylogenies

Sequences were aligned using MAFFT v.7 (Katoh et al., 2017), and the alignment files are available as supplementary data. Phylogenies were estimated with maximum likelihood in IQ-Tree (Nguyen et al., 2015; Trifinopoulos et al., 2016). The ultrafast bootstrap and Shimodaira-Hasegawa approximate likelihood ratio test (Sh-aLrT) were used to measure node confidence (Hoang et al., 2018). The optimal model for each tree was selected using ModelFinder (Kalyaanamoorthy et al., 2017) based on the Bayesian Information Criterion. Mitochondrial phylogenies were estimated from the concatenation of all known mitochondrial genes; when they were combined with ribosomal loci (Figure 1), they were analyzed as a separate partition. The GTR+F+I+G4 model was selected for all ribosomal trees/partitions, while mitochondrial models differed: GTR+F+I+G4 for all concatenated mitochondrial genes, TN+F+I+G4 for cox1 only, and TIM2+F+G4 for nd1 only. Figures were produced by exporting IQ-Tree files to the Interactive Tree of Life webserver (Letunic and Bork, 2019).

Single-locus phylogenies of cox1, nd1, and 28S were made with a combination of Sanger sequenced loci and Illumina sequences that were trimmed to a single locus. We used the NCBI taxonomy browser and blast to assemble all available sequences related to the Halichondriidae, along with appropriate outgroups. For cox1, sequences were excluded if they did not include most of the Folmer barcoding region. For 28S, sequences in the D1 phylogeny were excluded if they did not include the highly variable C2-D2 barcoding region. We named four clades of the phylogeny, to facilitate discussions of component taxa, as shown on figures 1 and 3. Alignment files are available as supplementary files in Data Dryad.

## Results and Discussion

### Molecular phylogenies

Our phylogeny for the 75 samples with the most comprehensive data includes both the nuclear ribosomal locus (averaging 6,445 bp) and the coding regions of the mitochondrial genome (averaging 17,153 bp; Figure 1). Previously, comparable data was available for only one species, *Ha. okadai*, which is included in the analysis. We successfully assembled nearly complete nuclear ribosomal loci for an additional 28 samples, including several additional species.

However, we did not have sufficient reads to assemble the mitochondrial genome for these samples. Mitochondrial and ribosomal phylogenies are shown separately for the Bowerbanki group (Figure 2). For the remaining specimens, we constructed a ribosomal-only phylogeny (Figure 3).

**Figure 2.**
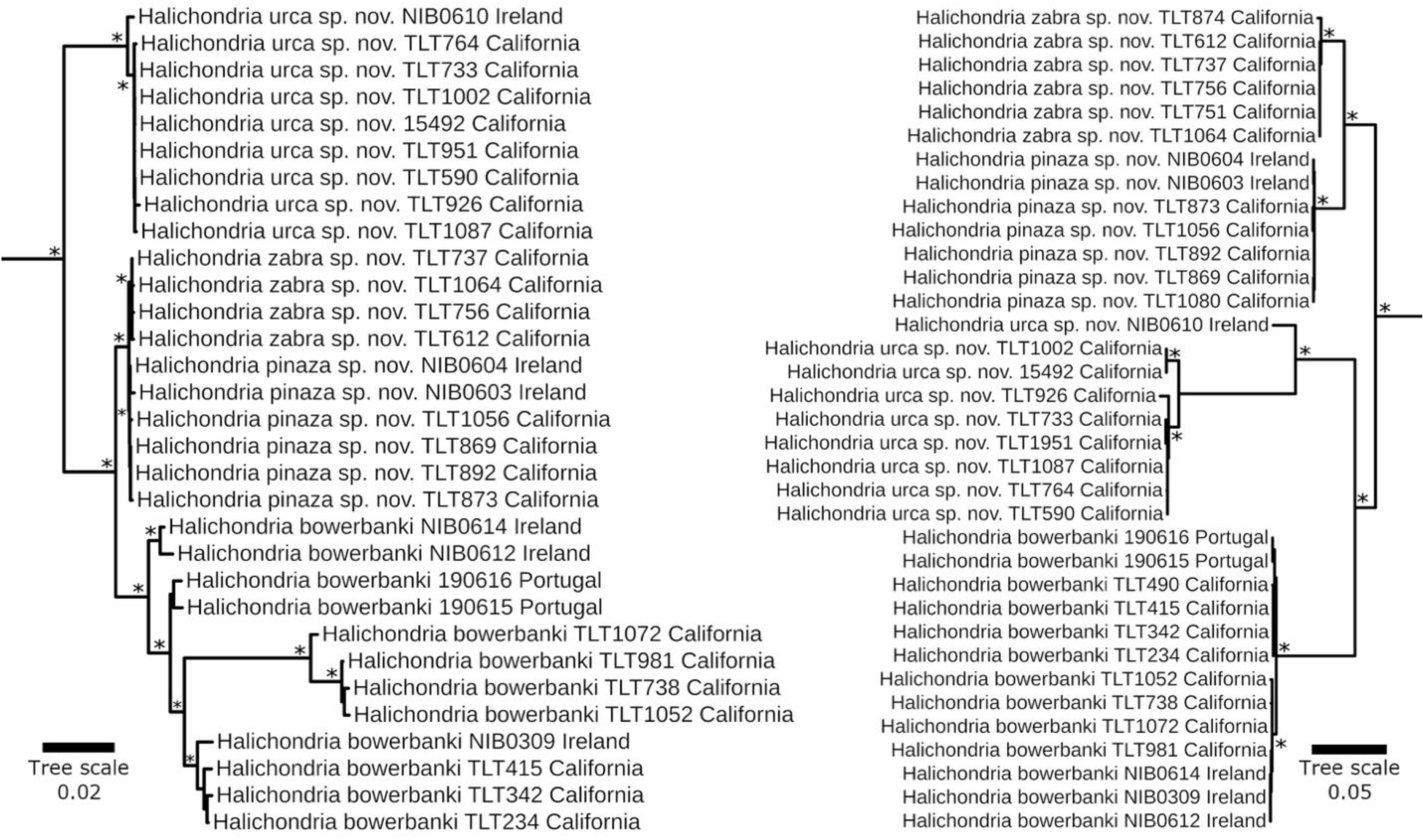
Maximum likelihood phylogenies for the mitogenome (left) and ribosomal locus (right) for the Bowerbanki group. Asterisks at nodes indicate a UFbootstrap ≥ 99%. Scale bar indicates substitutions per site.

**Figure 3.**
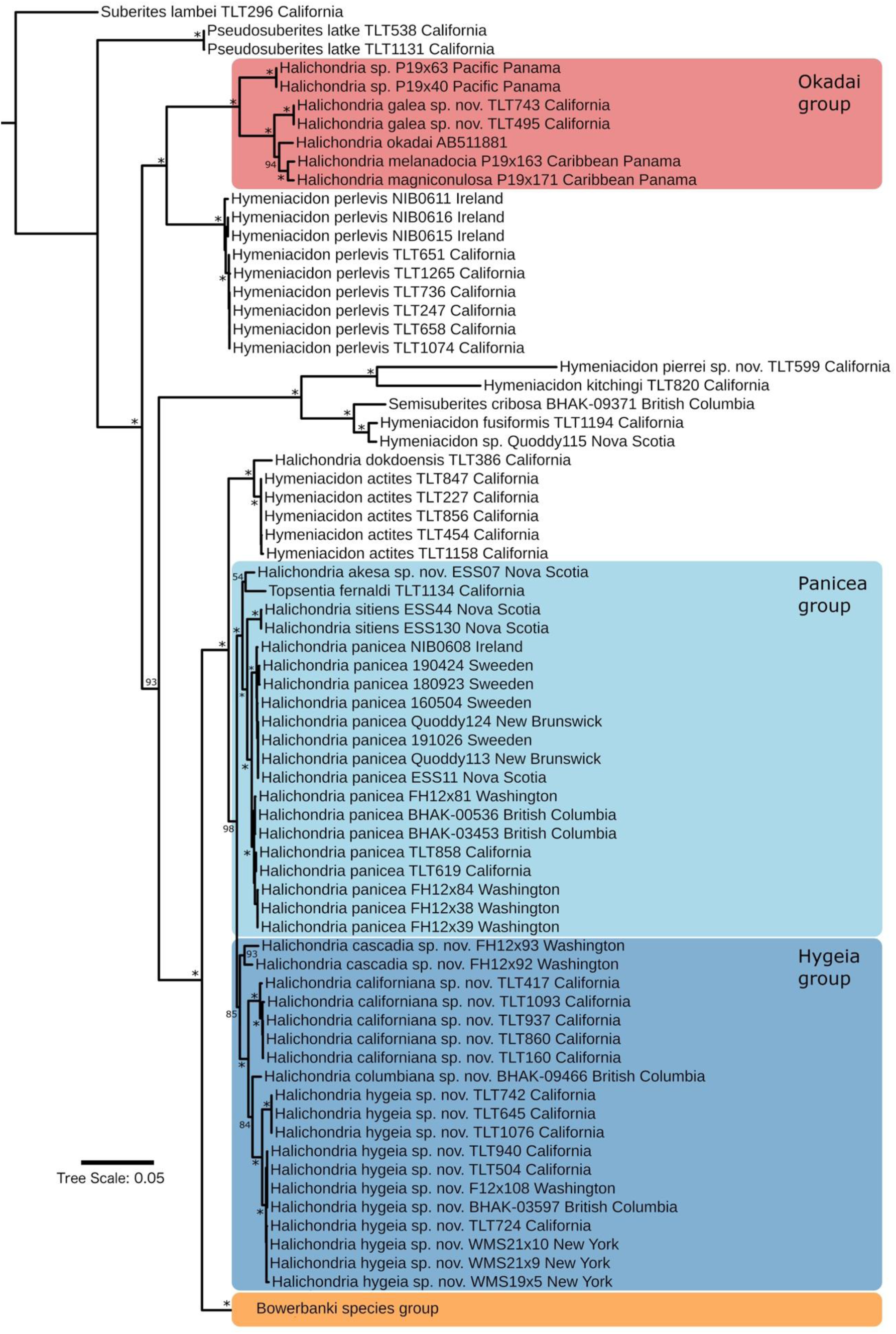
(above). Maximum likelihood phylogeny for samples with nearly complete ribosomal loci. Asterisks at nodes indicate a UFbootstrap ≥ 99%. Scale bar indicates substitutions per site.

Additionally, we constructed phylogenies using Sanger sequencing data from two mitochondrial loci (cox1 and nd1) and one nuclear locus (28S). These phylogenies were instrumental in validating genetic differences identified in the Illumina data and allowed us to include previously sequenced vouchers. In total, we sequenced at least one locus from 232 samples, and these phylogenies are included in the supplementary materials as (Figures S1-S3).

We consistently found highly differentiated clades across all phylogenies. Our first task was to determine which clade corresponded to the type species *Ha. panicea.* Due to the ambiguity of the original description, a neotype for the species was previously erected (Erpenbeck and van Soest, 2002). The neotype was collected from Burnham-on-Crouch, Essex, England, near the mouth of the Thames River, which is postulated to be near the original type location. Using sequence capture, Erpenbeck and colleagues generated 207 bp of sequence data from the hypervariable C2 region of the 28S barcoding locus from the neotype, and generously shared this previously unpublished sequence with us. Our 28S phylogeny confidently places this sequence within the clade designated *Ha. panicea* (Figure 1). It is an exact match to all three of the other samples from the United Kingdom, as well as samples from both the Atlantic and Pacific coasts of North America. These perfectly-matched samples were all previously identified as *Ha. panicea* based on their morphology, range, and habitat, and all fall into the *Ha. panicea* clade (Figure 1), supporting the genetic determination that this clade corresponds to the name *Ha. panicea*.

Similarly, we used a previously sequenced sample (Ulster Museum voucher BELUM:Mc4003; GenBank vouchers HQ379241, HQ379316, HQ379382, and KC902247) to determine which clade corresponds to the name *Ha. bowerbanki*. This name replaces *Ha. coalita* of Bowerbank, which was preoccupied by *Ha. coalita* (Müller, 1776). The type material consists only of microscope slides of isolated spicules. Like *Ha. panicea*, the type location is near the mouth of the Thames. However, two of the slides Bowerbank originally examined were from sponges collected at Strangford Lough, Northern Ireland. This location matches that of the Ulster Museum sample, which also aligns with the growth form described by Johnson (1842) and Bowerbank (1866). We are therefore confident that this clade corresponds to the name *Ha. bowerbanki*.

### Inference of reproductive isolation: Bowerbanki group

Our next task was to determine which clades in our phylogeny correspond to species other than *Ha. panicea* and *Ha. bowerbanki*. To infer reproductive isolation among species with overlapping geographic ranges, we compared the phylogenies of multiple loci that assort independently during meiosis. For example, the mitochondrial and nuclear ribosomal phylogenies for the clade we named the Bowerbanki group display reciprocal monophyly (Figure 2). Reciprocal monophyly in both genomic compartments allows us to infer a minimum of four species in this group: *Ha. bowerbanki* and three new species — *Ha. urca* sp. nov., *Ha. zabra* sp. nov., and *Ha. pinaza* sp. nov. (new species descriptions are provided in the systematics section below). Among these species, *Ha. zabra* sp. nov. and *Ha. pinaza* sp. nov. are the closest relatives. They are well-separated at the ribosomal locus, but Sanger sequencing of the *cox1* mitochondrial locus did not reveal fixed genetic differences between them (Figure S1). However, in the concatenated alignment of all mitochondrial coding regions, we identified 24 fixed differences between these clades, including one fixed difference in the cox1 gene outside the region bounded by our Sanger sequencing primers. With genetic divergence (Dxy) of only 0.25% across the mitochondrial genes, the branch separating these clades is short and difficult to visualize (Figure 2), but monophyly of each clade is well supported (99% and 100% bootstrap support for *Ha. pinaza* sp. nov. and *Ha. zabra* sp. nov., respectively).

To confirm that these genetic differences are consistent in a larger sample, we sequenced a region of the nd1 mitochondrial gene containing three apparent fixed differences. In a larger sample of 16 *Ha. zabra* sp. nov. and 7 *Ha. pinaza* sp. nov. samples, these differences remained fixed. Similarly, we sequenced the most variable region of the 28S nuclear ribosomal locus in 23 *Ha. zabra* sp. nov. and 7 *Ha. pinaza* sp. nov. samples, confirming numerous fixed differences (divergence = 1.32%). This co-assortment of genetic variation in the mitochondrial and nuclear genomes is strong evidence of reproductive isolation due to the co-occurrence of these species in California. They are sympatric even at a micro-geographic scale, with samples collected side-by-side on the same floating dock in Tomales Bay in Northern California, and on floating docks in Santa Barbara Harbor, over 500 km to the south. Likewise, *Ha. urca* sp. nov. and *Ha. bowerbanki* were sympatric at a micro-geographic scale. All four species in this species group were collected simultaneously from floating docks in Santa Barbara Harbor, with varying degrees of sympatry across other harbors and marinas throughout the state. There were numerous alleles fixed within the *Ha. urca* sp. nov. and *Ha. bowerbanki* clades in both genetic compartments, with monophyly highly supported in both cases (Figure 2).

Our inference of four species within the Bowerbanki group is likely conservative, as phylogenetic analyses suggest at least one additional species. Within the clade we designate as *Ha. urca* sp. nov., one sample from Ireland is genetically distinct. Unlike the *Ha. pinaza* sp. nov. samples, where Irish and California specimens are nearly identical at the 28S locus (divergence = 0.2%), the Irish *Ha. urca* sp. nov. shows a 4.6% sequence divergence at this locus. A sample of *Ha. urca* sp. nov. from New York is more similar to the California samples than to the Irish one, indicating that there might be a single population spanning both ocean basins, with the Irish sample potentially representing a different species or a highly divergent population. Additional *Ha. urca* sp. nov. data from the Atlantic includes three sponges from Virginia (SERC/MarineGeo 2018 Lower Chesapeake BioBlitz, available in GenBank) and one from New York (Pankey et al., 2022). These samples, previously identified as *Ha. bowerbanki*, are recognized as *H. urca* sp. nov. in our cox1 phylogeny (Figure S1). Combining these specimens with our samples results in a total of 5 Atlantic and 16 Pacific *Ha. urca* sp. nov. at this locus, with a genetic divergence of 0.38% between them. Although this divergence is slight, it is eight times greater than the divergence observed between *Ha. zabra* sp. nov. and *Ha. pinaza* sp. nov., which we can confidently say are reproductively isolated in sympatry. The differentiation seen in cox1 for Atlantic versus Pacific *Ha. urca* sp. nov. is higher than for any other species found in both basins (Fst = 0.64 for *Ha. urca* sp. nov., compared to 0.00–0.58 for the six other species with distributions across both oceans). Some or all Atlantic populations of *Ha. urca* sp. nov. may therefore be from an additional species, but we conservatively assign all Atlantic samples to *Ha. urca* sp. nov. until additional data are collected.

We also observed considerable variation in the mitochondrial genome within the clade we assign to *Ha. bowerbanki* —far more variation than seen within any of the other species in this study (Figure 2). In contrast to the clades we designate as species, however, the mitochondrial clades within *Ha. bowerbanki* were not mirrored by divergence in the nuclear locus. This lack of divergence could occur because there are multiple species within this clade, but they are too recently diverged to have accumulated fixed differences in the ribosome (Rittmeyer and Austin, 2012). Alternatively, this species may have a larger effective population size, higher mitochondrial mutation rate, or balancing selection that has maintained mitochondrial variation within a single species (Ballard and Whitlock, 2004; Galtier et al., 2009). We therefore assign all members of this clade to *Ha. bowerbanki*.

Finally, we note that there is disagreement between the mitochondrial and nuclear compartments regarding the branching order of the species in this group. Mitochondrial genes place *Ha. bowerbanki* as sister to the clade containing *Ha. zabra* sp. nov. and *Ha. pinaza* sp. nov., while the nuclear data position *Ha. bowerbanki* as sister to *Ha. urca* sp. nov.. This discrepancy could result from mitochondrial introgression, incomplete lineage sorting, or homoplasy. Although this discordance affects the inference of historical relationships, it does not impact conclusions about current reproductive isolation and is the only instance of differing branching orders between mitochondrial and nuclear genes in our Illumina-scale dataset.

### Inference of reproductive isolation: Panicea group

*Halichondria panicea* was described from the North Atlantic, and there has been debate about whether North Pacific populations belong to the same species (Erpenbeck, 2004; Erpenbeck and van Soest, 2002). Our analysis reveals that Pacific and Atlantic populations of *Ha. panicea* are sister clades distinguishable at the DNA level (Figures 1 & 3). Fixed genetic differences between these ocean basins are evident in both nuclear and mitochondrial genomes, including two fixed differences in each of the three Sanger-sequenced loci (cox1, nd1, and 28S). The genetic divergence between Atlantic and Pacific populations of *Ha. panicea* at cox1 is similar to that observed between Atlantic and Pacific populations of *Ha. urca* sp. nov. (0.40% divergence for both; Fst = 0.58 for *Ha. panicea*, Fst = 0.64 for *Ha. urca* sp. nov.). Unlike cases where populations are sympatric, this divergence does not confirm that these populations are reproductively isolated (Rannala and Yang, 2020). Indeed, *Ha. panicea* have been reported from the Arctic. No genetic data are available to confirm the identity of Arctic *Halichondria*, but if these populations are indeed *Ha. panicea*, it is possible that populations from the Atlantic and Pacific meet and mix in these Northern waters. We therefore leave the North Pacific population of *Ha. panicea* within this species, pending further data. We note that a previous paper that attempted to compare *Ha. panicea* in the Atlantic and Pacific only used two samples from the Atlantic, and alignment of those samples to our data reveal that the Atlantic samples were, in fact, *Ha. bowerbanki* (Erpenbeck, 2004).

As noted in the morphological section, there are multiple cryptic species that are morphologically indistinguishable from *Ha. panicea*. It is therefore surprising that, of the three species most closely related to *Ha. panicea*, two are morphologically distinct. The most closely related species is *Ha. sitiens* (Schmidt, 1870), which, like *Ha. panicea*, is found in both the North Pacific and North Atlantic. *Ha. sitiens* is clearly differentiated from *Ha. panicea* by spicule size and the presence of surface papillae. These papillae have historically been used to define subgenera within *Halichondria*, with *Ha. sitiens* as the type species of subgenus *Eumastia* and *Ha. panicea* as the type species of subgenus *Halichondria* (Erpenbeck and van Soest, 2002). The close relationship between these two species clearly invalidates these subgenera. Although only one *Ha. sitiens* was sequenced using Illumina technology, two Pacific and four Atlantic samples were sequenced at each of the three Sanger loci. In contrast to *Ha. panicea*, all *Ha. sitiens* samples are nearly identical, showing minimal genetic diversity both within and between populations (divergence between ocean basins <0.1% at each locus, with no fixed differences).

Outside of the *Ha. panicea* + *Ha. sitiens* clade were two other species; short branches did not allow us to resolve their branching order. One of these is represented by a single sample that is morphologically indistinguishable from *Ha. panicea.* This sample (ESS07) was collected at the same time and place as three other samples that were found to be *Ha. panicea* (Brig’s Shoal, in the Eastern Shore Islands of Nova Scotia). Though this clade is represented by only a single sample in our collections, the mitochondrial genes are nearly identical (99.9%) to a previously published mitochondrial genome from Iceland (GenBank accession MH756604).

Previous work ascribed this genotype to *Ha. panicea* (Knobloch et al., 2019), but the closer relationship of *Ha. sitiens* to true *Ha. panicea* clearly indicates that a new name for this taxon is needed. We describe it here as *Ha. akesa* sp. nov. This species differs from the *Ha. panicea* neotype DNA fragment at 3 sites (of 207); divergence vs. all *Ha. panicea samples* across the Sanger regions of cox1 and 28S is 0.50% and 0.80%, respectively.

The final species in the Panicea group serves to emphasize the inadequacy of spicule- based taxonomy in this family. *Topsentia fernaldi* (Sim and Bakus, 1986) has a very dense skeleton of long oxeas that make it quite distinct from *Ha. panicea*. The genus *Topsentia* is among the most polyphyletic of sponge genera, with previous molecular phylogenies placing some *Topsentia* species very far from other members of the Halichondriidae, in a sister clade to the order Axinellida (Pankey et al., 2022; Redmond et al., 2013). Reconciling the taxonomy and phylogeny of this genus (and the other polyphyletic genera included here) will require data from additional species, especially type species, and is beyond the scope of this paper.

### Inference of reproductive isolation: Hygeia group

The most widely distributed species in the clade we dub the Hygeia group is the newly described *Ha. hygeia* sp. nov., which was found throughout the Northeast Pacific and in the Northwest Atlantic. In the Pacific, its range overlaps broadly with another new species, *Ha. californiana* sp. nov. These two species are distinctly separated at the ribosomal locus, showing 14 fixed differences (1.5% divergence) in the Sanger-sequenced region of 28S (Figure S2).

Although they could not be differentiated in the Sanger-sequenced region of the cox1 mitochondrial locus (Figure S1), Illumina sequencing revealed numerous other fixed differences in the mitochondrial genome. Sanger sequencing of the nd1 mitochondrial gene (Figure S3) confirmed 4 fixed differences in a larger sample (0.8% divergence, Fst = 0.73, N = 12 *Ha. californiana* sp. nov. vs. 17 *Ha. hygeia* sp. nov.). The fixed differences observed in both genomic partitions, coupled with their broad sympatry, strongly supports reproductive isolation in nature.

Two additional species are found in this group, both restricted to the Pacific Northwest of North America. Despite the limited number of samples (N=2 for each species), both are well differentiated from other species in the group based on mitochondrial and nuclear loci. Two *Ha*.

*cascadia* sp. nov. samples were sequenced using Illumina technology, and these samples are strongly supported as a sister clade to all other species in the group (Figure 1). Given that *Ha. hygeia* sp. nov. and *Ha. californiana* sp. nov. are well-supported as isolated species, and *Ha. cascadia* sp. nov. is sympatric with *Ha. hygeia* sp. nov., it is likely that *Ha. cascadia* sp. nov. is also an isolated lineage deserving of species status. We also have two samples of *Ha. columbiana* sp. nov., both from British Columbia, though only one was successfully sequenced with Illumina technology. This sample was quite unique, with many alleles differentiating it from the sympatric species *Ha. hygeia* sp. nov.. This sample formed a clade with the second sample across all three Sanger-sequenced loci, providing further evidence that this distinct clade is likely a separate species. Our inference of four species within the Hygeia species group is likely conservative, as phylogenetic analyses suggest at least one additional species. Three samples (TLT1076, TLT742, TLT645) formed a reciprocally monophyletic clade at the nuclear ribosomal locus and the mitochondrial nd1 locus (Figures S2 and S3). Given the limited sample sizes and the close relationship of these samples with other *Ha. hygeia* sp. nov., we conservatively include these samples within the species concept for *Ha. hygeia* sp. nov., pending further data.

### Additional Halichondria species found to span the temperate North Pacific

The first mitochondrial genome published for a *Halichondria* species was for a "*Halichondria sp.*" sponge sampled from Fujian Province, China (Wang et al., 2016). Sequencing of mitochondrial genes later documented this same genotype from the intertidal zone of mainland South Korea and nearby islands, and from Mission Bay, San Diego, California (Park et al., 2007). This sponge has remained undescribed, likely due to it being morphologically indistinguishable from *Ha. panicea*. Our Southern California samples have mitochondrial genomes that are nearly identical with the published sequence from China (99.95% identical), confirming that we have recollected this same species from California. Consistent with previously published data, it is only distantly related to *Ha. panicea*, despite their morphological similarity. The clade we designate here as the Okadai species group contains *Ha. okadai* and *Ha. galea* sp. nov., which are sympatric in the Korean Strait; the Caribbean species *Ha. magniconulosa* and *Ha. melanodocia*; and samples we collected from Pacific Panama that are likely from an undescribed species (Figure 3).

Remarkably, an additional species of *Halichondria* known from Asia was also discovered in California for the first time: *Ha. dokdoensis* Kang, Kim & Kim, 2022. A mitochondrial genome sequence was previously published for this species (Kim et al., 2019), and this sequence is nearly identical to the mitochondrial genome of our California sample: they differ by only 12 of 13,033 aligned base pairs (<0.1%). These samples are more closely related to *Hymeniacidon actites* than to any other sequenced *Halichondria*, and together these two species form a clade that is the sister to the combined Panicea and Hygeia species groups. We discovered a single sample of this species in California, on an oil extraction platform off the coast of Santa Barbara. As discussed in the systematics section, a publicly available sequence of the cox1 locus indicates that the species is also likely found in the Netherlands. These discoveries serve to highlight that there are likely additional species of widespread, introduced *Halichondria* that will be discovered by additional genotyping.

### Taxa are not diagnosable with traditional morphological characters

Our multi-locus molecular data, combined with broad sympatry, allows us to form strong inferences regarding which clades are species. We can use these well-delineated species to determine if known morphological characters are sufficient for their diagnosis. We first considered only the previously named species *Ha. panicea* and *Ha. bowerbanki*. Previous attempts to separate these species cite several differences including 1) longer spicules in *Ha. bowerbanki*, 2) a difference in spicule length in the choanosome vs. the ectosome in *Ha. panicea* but not *Ha. bowerbanki*, and 3) differences in the arrangement of spicules in the ectosome between species (Ackers et al., 2007; Hartman, 1958). These previous works did not use DNA to diagnose samples, so it is likely that samples from the previously unknown species discovered and described here were included in past work. Nonetheless, as detailed further below, we find that these traits do differ, on average, between these species. However, we also find variability within species that prevents them from being used to reliably diagnose all samples.

Spicules in *Ha. bowerbanki* were, on average, nearly 40% longer than those in *Ha. panicea* (Wilcoxon p<0.0001, Figure 4). However, substantial overlap of these distributions prevents this character from being used as a species diagnostic (Figure 4). Mean spicule length ranged from 202 to 357 μm for *Ha. panicea* and from 307 to 423 μm for *Ha. bowerbanki*. Both species had high variability in mean spicule length within the Atlantic and Pacific basins. We also find that spicules are, on average, 15% shorter in the ectosome than the choanosome for *Ha. panicea* but not for *Ha. bowerbanki* (Figure S4). Again, this trait proved insufficient for diagnosing all samples due to intraspecific variability. Some *Ha. panicea* specimens had spicules 30% longer in the choanosome, and two samples (one from Nova Scotia and one from Sweden) had spicules of equal length in both compartments. To verify that this variability within species was not entirely due to technical variability and sampling error, we reisolated spicules from resampled tissues. One *Ha. panicea* sample with no initial difference in spicule lengths by compartment and another with a 32% difference were re-examined. Replicate data confirmed these results: the sponge with no initial difference had only a 1% difference in the replicate (non-significant), while the sponge with a 32% difference initially showed a 26% difference in the replicate (p<0.001). In contrast, *Ha. bowerbanki* samples exhibited less variability, with ectosomal spicules varying from about 10% longer to about 10% shorter than choanosomal spicules. Attempts to replicate samples of *Ha. bowerbanki* with the greatest observed differences (11% longer choanosomal spicules vs. 9% longer ectosomal spicules) showed both samples regressing towards a mean difference of zero, consistent with the possibility that variation in *Ha. bowerbanki* is largely due to tissue sampling variation and statistical noise.

**Figure 4.**
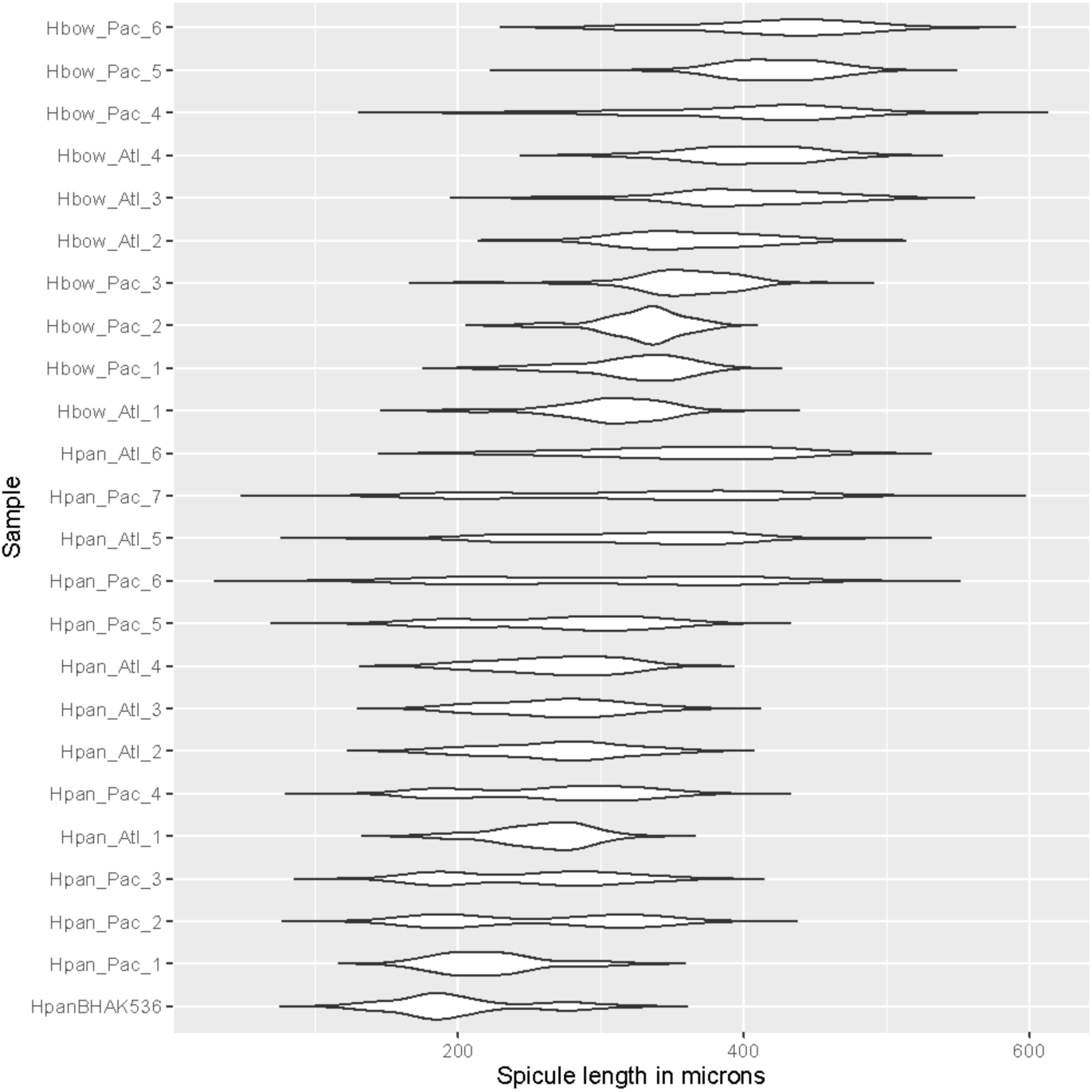
Spicule length distributions for samples of Halichondia panicea (Hpan) and Ha. bowerbanki (Hbow). Sample are sorted by mean within species, and names indicate if the sample was from the Atlantic (Atl) or Pacific (Pac). Hpan samples generally have shorter average lengths and bimodal distributions, due to length differences between the ectosome and choanosome, but some Hpan samples are longer than the shortest Hbow samples, and bimodality is not always present.

When average spicule length and length differential between compartments are plotted together, the average differences between *Ha. panicea* and *Ha. bowerbanki* are clear, and it appears likely that a combination of these two traits would be sufficient to differentiate most, but probably not all, samples (Figure 5). However, when we added the other species described here to this plot, we observed substantial overlap among them.

**Figure 5.**
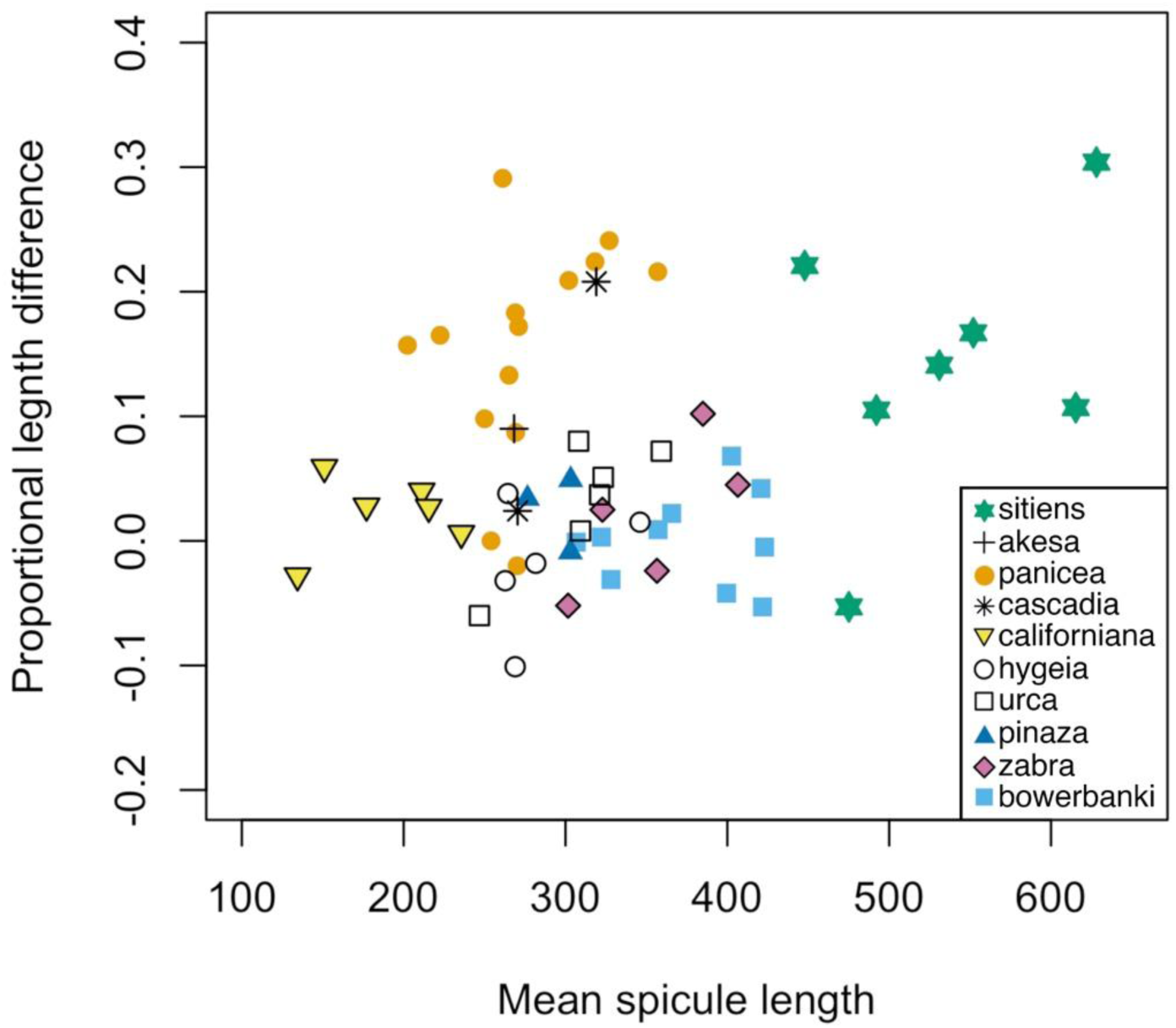
Spicule measurements across Halichondria species. Only samples where spicules were measured separately for the ectosome and the choanosome are shown. Mean spicule length is the mean for both domains combined, and proportional length difference is the choanosomal mean minus the ectosomal mean over the sample mean.

The most comprehensive publication on the morphological distinction between *Ha. panicea* and *Ha. bowerbanki* states that the arrangement of spicules in the ectosome is the most reliable diagnostic character (Hartman, 1958). Typically, *Ha. bowerbanki* exhibits a network of long, thin spicule tracts that delineate larger open spaces, while *Ha. panicea* has thicker spicule tracts forming a dense mesh surrounding smaller openings. *Ha. panicea* sometimes also features "gothic crypt-like supports" connecting the ectosomal and choanosomal skeletons (Hartman, 1958). Although these differences were observed in our samples, they proved insufficient for diagnosing all samples reliably. This trait is highly variable within a single sponge and among samples within species. While mean tract thickness differed significantly between species (69 μm for *Ha. panicea* vs. 38 μm for *Ha. bowerbanki*, Wilcoxon p<0.0001; Figure S5), there is considerable overlap in the distributions (e.g., *Ha. panicea* 39–97 μm, *Ha. bowerbanki* 29–56 μm; figure S5). We also attempted to categorize ectosomal skeletons into one of five structural types (Figure S6); this categorization again highlighted average differences but also considerable overlap. Principal components analysis combining ectosomal tract thickness, average spicule length, and length differential between compartments did not enhance the ability of these traits to diagnose species (Figure S7). Even when considering a small number of samples of only *Ha. panicea* and *Ha. bowerbanki*, separation was incomplete.

Other morphological differences between (apparent) *Ha. panicea* and *Ha. bowerbanki* have been proposed, such as smell (subjective and only available in fresh material), rigidity of branches (branches often not present), color, and general form (Ackers et al., 2007; Hartman, 1958). It is likely that these traits and others will be partially informative, but our efforts to differentiate samples based on these traits has proved inconsistent and frustrating. Moreover, many morphological traits likely respond plastically to environmental conditions such as light intensity, wave energy, and current strength (Palumbi, 1986). Some species are found primarily in sheltered harbors and marinas, while others are found in wave-swept intertidal habitats, so average morphological differences between them may be due more to habitat than genotype. For instance, a sample of *Ha. urca* sp. nov. from an intertidal zone exhibited an ectosomal skeleton typical of *Ha. panicea*, which is common in high-energy environments, whereas most *Ha. urca* sp. nov. specimens from sheltered marinas displayed a sparsely spiculated ectosome (see *Ha. urca* sp. nov. description below). Therefore, we conclude that the previously named species *Ha. panicea* and *Ha. bowerbanki* cannot be reliably distinguished based on known morphological traits alone, and the newly described *Halichondria* species should be considered cryptic species without clear morphological differences.

## Systematics

The aims of this section are two-fold. First, we formally describe the *Halichondria* species that were found to be morphologically indistinguishable from *Ha. panicea*. Second, we aim to comprehensively revise the taxonomy of the family Halichondriidae for the Northeast Pacific, from British Columbia to California. This revision encompasses all newly discovered and previously described species within this region. The only species included here that has not been found in this area is *Ha. akesa* sp. nov., which is known solely from the North Atlantic.

Diagnoses include DNA character states with the format "number:letter", where the number indicates a position in one of the DNA alignments provided (where the first base is number 1), and the letter denotes the base state at that position, which is unique compared to all other species in the genus for which sequence data are available. The notation "number- number" indicates a sequence of DNA starting at the first number and proceeding to the last number. The sample sizes (N) for each character are indicated; these can be variable as some samples were sequenced at more loci than others, or because of the presence of missing data in the sequences. Although many species of *Halichondria* lack DNA data, these previously described species were all morphologically distinguished from *Ha. panicea* and are therefore morphologically distinct from the newly described species as well.

In practice, informal diagnosis of all species is feasible by sequencing the D1-D3 region of the 28S locus and constructing a phylogeny with the provided sequences. However, clade membership alone is not sufficient for formal description under the Code of Zoological Nomenclature, and not all species exhibit unique derived states at this locus. Consequently, formal diagnosis requires sequencing different loci for some species. Morphological traits should be employed to diagnose species when comparing them to previously described *Halichondria* that have not been sequenced at these loci.

## List of species discussed in systematics section Order Suberitida, Family Halichondriidae

Genus Halichondria

*Halichondria panicea* (Pallas, 1766)

*Halichondria akesa* sp. nov. *Halichondria sitiens* (Schmidt, 1870) *Halichondria hygeia* sp. nov.

*Halichondria californiana* sp. nov.

*Halichondria cascadia* sp. nov.

*Halichondria columbiana* sp. nov.

*Halichondria dokdoensis* Kang, Kim & Kim 2022

*Halichondria galea* sp. nov.

*Halichondria bowerbanki* Burton, 1930

*Halichondria urca* sp. nov. *Halichondria zabra* sp. nov. *Halichondria pinaza* sp. nov.

*Halichondria loma* Turner & Lonhart 2023

Genus Hymeniacidon

*Hymeniacidon perlevis* (Montagu, 1814) *Hymeniacidon actites* (Ristau, 1978) *Hymeniacidon ungodon* de Laubenfels 1932 *Hymeniacidon kitchingi* (Burton 1935) *Hymeniacidon pierrei* sp. nov.

*Hymeniacidon fusiformis* Turner & Lonhart 2023

*Hymeniacidon globularis* Ott, McDaniel & Humphrey 2024

Genus Semisuberities

*Semisuberites cribrosa* (Miklucho-Maclay, 1870)

Genus Topsentia

*Topsentia fernaldi* (Sim & Bakus, 1986)

*Topsentia disparilis* (Lambe 1894)

Genus Axinyssa

*Axinyssa piloerecta* sp. nov.

*Axinyssa tuscara* (Ristau 1978)

## Family Halichondriidae

The Systema Porifera defines the Halichondriidae as members of order “Halichondrida with a confused arrangement of smooth oxeas and/or styles in the choanosome and usually an organized special ectosomal skeleton consisting of tangentially arranged or densely confusedly arranged crust of oxeas and/or styles of sizes similar to or smaller than those of the choanosome” (Erpenbeck and Van Soest, 2002). Order Halichondrida was abandoned based on a lack of synapomorphic characters and its families were distributed into several orders. Family Halichondriidae joined families Suberitidae and Stylocordylidae in order Suberitida (Morrow and Cárdenas, 2015).

### Genus Halichondria

Halichondriidae with a tangential ectosomal skeleton carried by subectosomal spicule tracts or brushes separated by subdermal spaces. Megascleres exclusively oxeas or derivates in a wide size range (paraphrased from Erpenback and Van Soest, 2002). This broadly defined genus includes over 95 species (de Voogd et al., 2024).

### Halichondria panicea (Pallas, 1766)

Figures 6 & 7

## Synonyms

*Spongia panicea* Pallas, 1766

The World Porifera Database lists over 70 additional synonyms (de Voogd, et al., 2024)

## Material examined

160504, Spettan, Gullmarsfjorden, Sweden, (58.25770, 58.25770), 18 m, 2015-05-04; 160505, Spettan, Gullmarsfjorden, Sweden, (58.25770, 58.25770), 18 m, 2015-05-04; 180911, Svartejan, Idefjord, Sweden, (59.11020, 11.32190), 30 m, 2018-09-11; 180923, Lunneviken, Idefjord, Sweden, (59.05460, 11.16900), 27 m, 2018-09-23; 181204, Svartejan, Idefjord, Sweden, (59.11020, 11.32190), depth not recorded, 2018-12-04; 190424, Flatholmen, Gullmarsfjorden, Sweden, (58.26080, 11.39820), 14 m, 2019-04-24; 190624, Trindeknubben, Idefjord, Sweden, (58.78290, 10.99620), 12 m, 2019-06-24; 191026, Gamla Svinesundsbron, Idefjord, Sweden, (59.09770, 11.26860), 19 m, 2019-10-26; 191116, Yttre Vattenholmen, Idefjord, Sweden, (58.87540, 11.10560), 27 m, 2019-11-16; BHAK-00536, Fifth Beach, Shatner Point, Calvert Island, British Columbia, (51.63870, -128.15700), intertidal, 2017-07-25; BHAK-03453, Crazy Town, North Beach, Calvert Island, British Columbia, (51.66680, -128.13400), intertidal, 2017- 08-08; ESS11, Brig’s Shoal, Eastern Shore Islands, Nova Scotia, (44.62781, -62.93603), 25 m, 2021-07-19; ESS17, Brig’s Shoal, Eastern Shore Islands, Nova Scotia, (44.62781, -62.93603), 25 m, 2021-07-19; ESS18, Brig’s Shoal, Eastern Shore Islands, Nova Scotia, (44.62781, -62.93603), 25 m, 2021-07-19; ESS74, South of Brig’s Shoal, Nova Scotia, (44.60336, -62.92724), 29 m, 2021-07-23; ESS80, South of Brig’s Shoal, Nova Scotia, (44.60336, -62.92724), 29 m, 2021-07-23; FH12x36 (YPM 111889), Cattle Point, San Juan Island, Washington, (48.45030, -122.96237), intertidal (rocks), 2012-05-19; FH12x38 (YPM 111890), Cattle Point, San Juan Island, Washington, (48.45030, -122.96237), intertidal (rocks), 2012-05-19; FH12x39 (YPM 111891), Cattle Point, San Juan Island, Washington, (48.45030, -122.96237), intertidal (rocks), 2012-05- 19; FH12x81 (YPM 111892), Eagle Cove, San Juan Island, Washington, (48.46128, -123.03155), intertidal (rocks), 2012-05-22; FH12x84 (YPM 111893), Eagle Cove, San Juan Island, Washington, (48.46128, -123.03155), intertidal (rocks), 2012-05-22; FH12x85 (YPM 111894), Eagle Cove, San Juan Island, Washington, (48.46128, -123.03155), intertidal (rocks), 2012-05-22; FH12x86 (YPM 111895), Eagle Cove, San Juan Island, Washington, (48.46128, -123.03155), intertidal (rocks), 2012-05-22; NIB0608, Carrick-a-Rede, Northern Ireland, (55.24238, -6.33396), 10m (growing on kelp stipe), 2021-10-17; Quoddy113, Bald Head, Campobello, New Brunswick, (44.91220, - 66.95215), 11.6 m, 2022-10-31; Quoddy124, Marble Island, Off Indian Island, Western Isles, New Brunswick, (44.91921, -66.97054), 10.6 m, 2022-11-01; TAT026, Strawberry Draw Overhang, Tatoosh Island, Washington, (48.39190, -124.73800), intertidal, 2018-06-16; TAT031, Toad Point, Tatoosh Island, Washington, (48.38970, -124.73500), intertidal, 2018-06-17; TAT032, Toad Point, Tatoosh Island, Washington, (48.38970, -124.73500), intertidal, 2018-06- 17; TLT1293, Cape Alava, Washington, (48.17134, -124.75227), 4-12 m, 2023-07-29; TLT1370, Cape Johnson, Washington, (47.97327, -124.68220), 4-7 m, 2023-07-27; TLT1371, Cape Johnson, Washington, (47.97327, -124.68220), 4-7 m, 2023-07-27; TLT1430, Neah Bay, Washington, (48.38676, -124.63223), 6-14 m, 2023-08-01; TLT619, Scott’s Creek Beach, California, (37.04213, -122.23366), beach wrack, 2020-11-14; TLT858, Cave Landings, California, (35.17535, -120.72240), intertidal, 2021-02-06; ZMA6155, Jennny’s Cove, Lundy Island, England, 24 m, 1985-01-08.

## Diagnosis

Yellow or green *Halichondria* with a tangential ectosomal skeleton developed as a dense sieve-like mat of spicules or a lace-like network of spicule tracts. Choanosomal skeleton is confused with vague spicule tracts, sometimes accompanied by well-developed subectosomal spicule tracts descending from the ectosomal skeleton. Spicules oxeas, ranging from 100–500 μm in length, with averages per sample of 200–400 μm in length and 5–10 μm in width; usually slightly shorter in the ectosome than the choanosome. Diagnosis relative to similar species requires DNA characters, as follows: COX1. 754:A, 952:A. N=55–69 depending on position. These characters differentiate *Ha. panicea* from all other sequenced *Halichondria* except *Ha. sitiens,* which is diagnosed relative to *Ha. panicea* based on morphological characters such as spicule length and surface papillae.

28S. No unique derived states shared by all *Ha. panicea*, but the Atlantic Ocean population has two unique states, 605:T and 1549:C. N=13–21 depending on position ND1. No unique derived states shared by all *Ha. panicea*, but the Pacific Ocean population has a unique state, 451:T. N=10).

## Morphology

Thinly or thickly encrusting, sometimes with digitate tendrils (Figure 6). Oscula usually prominent, but vary from raised rims nearly flush with surface to tall oscular chimneys. Color alive varies from pale translucent yellow to golden, often variegated with green, or with the exterior entirely green or gray-green and the interior yellow. White or beige in ethanol.

**Figure 6.**
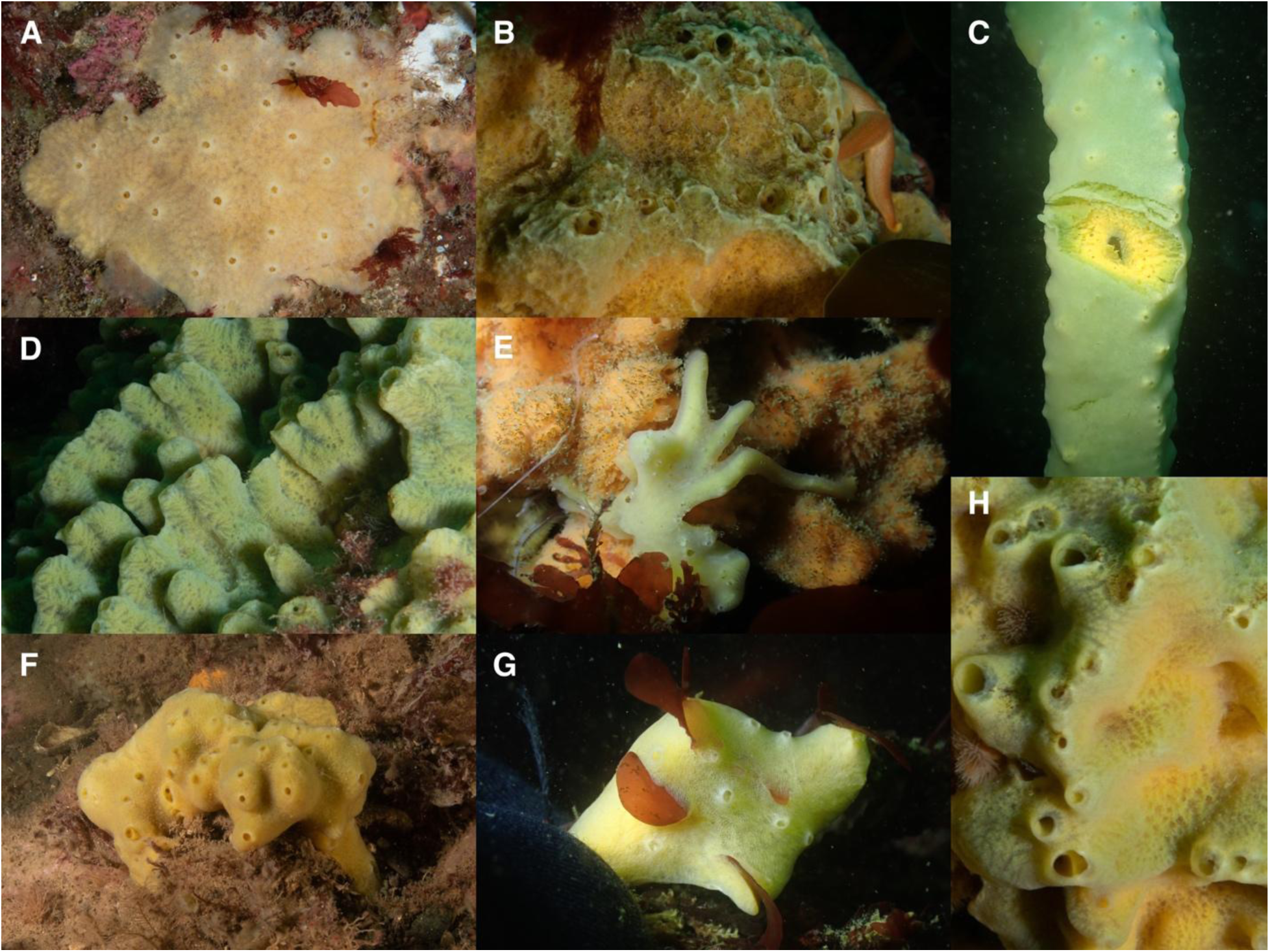
Halichondria panicea. Northwest Atlantic: A (ESS80), B (ESS17), D (ESS18), F (Quoddy124), H (ESS11). Northeast Pacific: C (TLT1430), E (TLT1293), G (TLT1371). All are photographed in situ; C shows wound where sample was taken to display internal color.

## Skeleton

Tangential ectosomal skeleton can be sieve-like, with a dense mat of spicules surrounding open areas that are mostly circular or oval shaped (Figure 7A, 7F), or more lace-like, with a web of spicule tracts surrounding open areas that tend towards triangular (Figure 7C, 7D). In some samples, this tangential ectosomal skeleton is supported by prominent spicule tracts that rise from the choanosome to fan out tangentially in the ectosome (Figure 7B, 7G). The choanosomal skeleton in these samples becomes more confused towards the interior (Figure 7H). Other samples have a confused jumble of multispicular tracts, spicule bundles, and individual spicules both near the ectosome and interior (Figure 7I). No apparent spongin.

**Figure 7.**
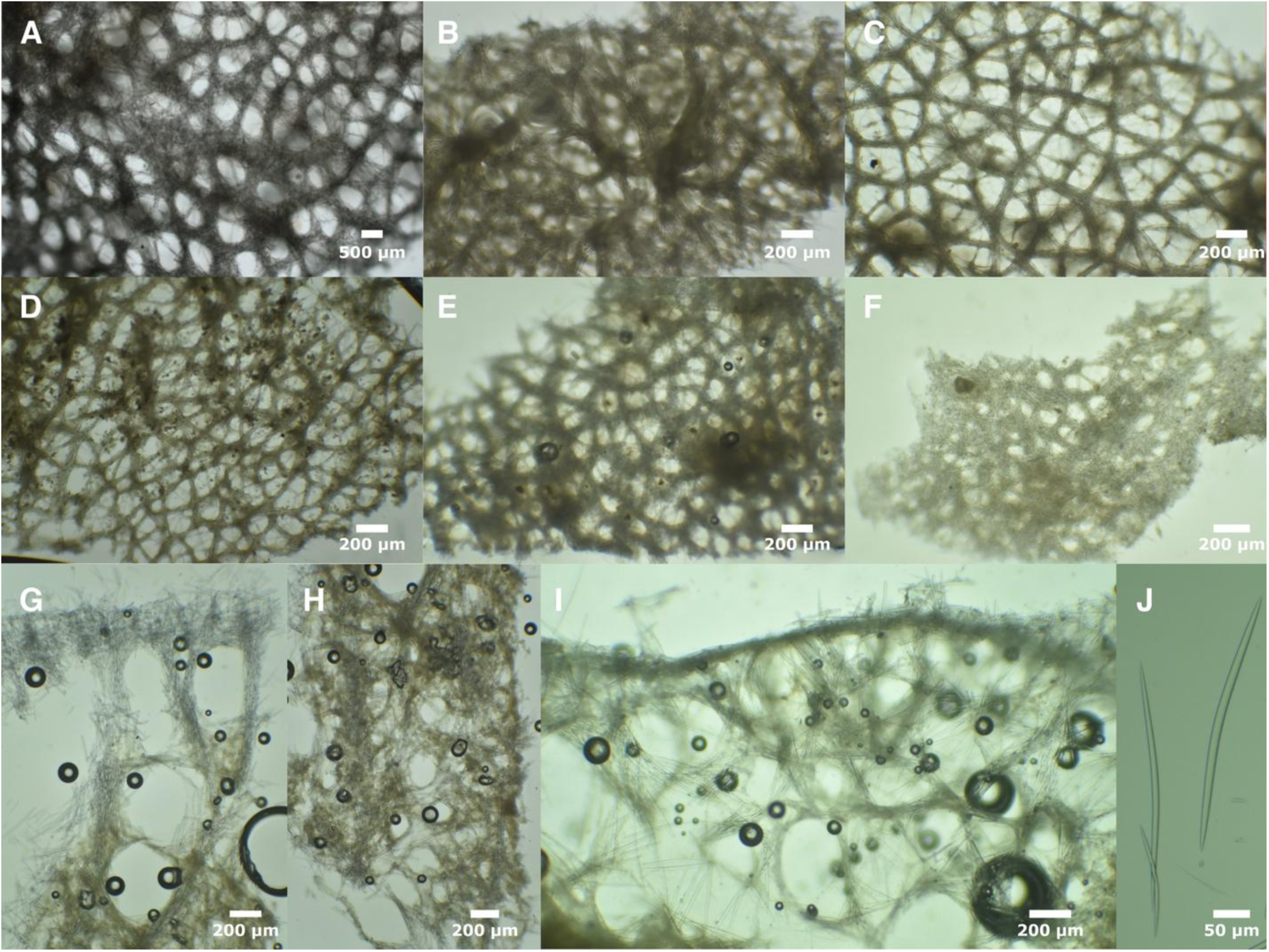
*Halichondria panicea* skeleton and spicules. Ectosomal skeletons: A (ESS80), B (TLT1430), C (FH12x38), D (Quoddy124), E (NIB608), F (TLT619). B is imaged from below to show the twisting columns of spicules connecting the ectosome to the choanosome in this sample. G: cross section at sponge surface, showing spicule columns and subectosomal spaces (TLT1430), H: cross section of same sample (TLT1430) farther towards the inferior, showing confused skeleton, I: cross section at sponge surface without large subectosomal spaces (TLT1370), J: oxeas (TLT1293).

## Spicules

Nearly exclusively oxeas, though rarely a spicule can be found that is modified into a style. Spicules have a typical halichondroid shape: evenly curved or with a slight central bend, thickest in the center and tapering gently to sharp points at both ends. Spicules have a large range in lengths in both the ectosome and the choanosome, but average 15.4% longer in the choanosome. Several samples (ESS11, 190424) had no difference in length by compartment (Figure S4). Dimensions are given for individual samples in table 1. When all samples are combined, spicules measure 114–272–499 x 2–8–16 μm, n=1502. Average values varied considerably across samples, with mean lengths from 202 to 358 μm, and mean widths from 5 to 10 μm.

**Table 1.** Spicule lengths of *Halichondria panicea*. Values given as min–mean–max in μm.

| Specimen | Ectosome | Choanosome |
| --- | --- | --- |
| Pacific Ocean |  |  |
| BHAK-00536 | 114–185–285 n=40 | 130–217–322 n=46 |
| BHAK-03453 | 153–204–256 n=40 | 170–241–322 n=40 |
| TLT619 | 130–270–451 n=40 | 199–334–442 n=40 |
| TLT1370 | 172–286–451 n=45 | 146–364–499 n=50 |
| TLT1430 | 147–233–353 n=96 | 162–299–367 n=98 |
| FH12x38 | 155–247–335 n=49 | 169–272–377 n=50 |
| FH12x39 | 134–247–357 n=42 | 169–291–367 n=47 |
| FH12x81 | 145–238–351 n=55 | 166–262–354 n=54 |
| Atlantic Ocean |  |  |
| NIB0608 | 176–257–322 n=49 | 187–281–365 n=51 |
| ZMA-6155 | 149–283–408 n=40 | 241–354–458 n=40 |
| 190424 | 178–273–347 n=40 | 196–267–342 n=40 |
| 180911 | 173–247–349 n=49 | 221–296–357 n=41 |
| Quoddy124 | 212–320–421 n=48 | 298–397–463 n=44 |
| ESS11 | 185–254–318 n=86 | 167–254–331 n=86 |

## Distribution and habitat

This species has been reported to occur from Iceland to the Mediterranean in the Northeast Atlantic, from Northern Canada to New England in the Northwest Atlantic, and possibly the Arctic and North Pacific (Erpenbeck and van Soest, 2002). DNA data is able to confirm only the Northern portion of this range in the Northeast Atlantic, including Iceland, Sweden, the United Kingdom, and the Netherlands. Genetic support in the Northwest Atlantic ranges from New Brunswick to New Hampshire. Sequenced samples from farther South in the Atlantic (Portugal, Connecticut, New York, Virginia) have thus far proved to be in the Hygeia or Bowerbanki species groups, but additional sampling could certainly extend this range. In the North Pacific, DNA-confirmed range is from the Bering Strait to Central California, with no DNA confirmation available from the Northwest Pacific. The species is rare or absent from shallow waters in Southern California, as the genotyping of over 100 *Halichondria* samples from this region found none to be *Ha. panicea*.

Previously, this species was described as common on rocks and brown algae in the intertidal and shallow subtidal zones, preferring exposed areas over sheltered bays, and extending down to a depth of 500 meters (Erpenbeck and van Soest, 2002). DNA-confirmed samples align with this description, although we cannot confirm a depth range beyond 29 meters. The species is common in exposed intertidal areas and the shallow subtidal zone but is rare or absent on human-made structures like floating docks in marinas, where *Ha. hygeia* sp. nov. and members of the Bowerbanki species group are prevalent.

## Remarks

As discussed in the genetic section, *Ha. panicea* from the Atlantic and Pacific basins form sister clades that are distinguishable at the genetic level and could potentially be considered separate species. However, if *Ha. panicea* occurs throughout the Arctic, it may link the populations across these basins. We conservatively retain both populations under the same species concept pending further data.

We lack genetic confirmation of *Ha. panicea* in the Northwest Pacific, but its presence there seems likely, as we have confirmed its distribution in the Aleutian Islands and the Bering Strait. This species has also been reported from various other global regions, including South America, New Zealand, and the tropical South Pacific (https://gbif.org). Several lines of evidence suggest that these records may actually pertain to other species, such as *Ha. hygeia* sp. nov., members of the Bowerbanki species group, or additional cryptic species that remain to be discovered. First, *Ha. panicea* seems less likely than those species to be experiencing a human-assisted range expansion for two reasons: 1) it does not seem to occur in human-modified habitats like marinas, and 2) genetic differentiation between Atlantic and Pacific populations make it likely that these populations do not share a common ancestor in the modern era.

Second, the species does not seem to tolerate warm temperate waters in Southern California, and is therefore unlikely to occur in shallow tropical waters.

Halichondria akesa sp. nov.

Figure 8

Material examined

Holotype: ESS07 (ARC 82019, CASIZ 245200), Brig’s Shoal, Eastern Shore Islands, (44.62781, - 62.93603), 25 m, 2021-07-19.

## Etymology

Name inspired by the Greek goddess Akeso, sister of Panakeia.

## Diagnosis

Yellow and green *Halichondria* with a tangential ectosomal skeleton developed as a thick mat of spicules and/or a lace-like network of spicules and spicule tracts. Choanosomal skeleton consists of prominent spicule tracts near the ectosome, transitioning to paucispicular tracts and spicules in confusion deeper in the choanosome. Spicules oxeas, ranging from 90–460 μm in length and 2–10 μm in width; slightly shorter in the ectosome than the choanosome. Diagnosis relative to similar species requires DNA characters, as follows:

COX1. No unique derived states. 28S. 621:T. N=1.

ND1. 39:T, 55:T. N=2.

## Morphology

The single specimen examined was thickly encrusting with prominent oscula. Yellow alive, white in ethanol.

## Skeleton

The single specimen examined has a tangential ectosomal skeleton which was a dense mat of spicules in some areas, and a network of tracts and single spicules surrounding oval-shaped open areas in other places. The ectosomal skeleton is supported by prominent spicule tracts that rise from the choanosome to fan out tangentially in the ectosome (Figure 8B, 8G). The choanosomal skeleton becomes more confused towards the interior, with some spicule tracts but many spicules in confusion. No apparent spongin.

**Figure 8.**
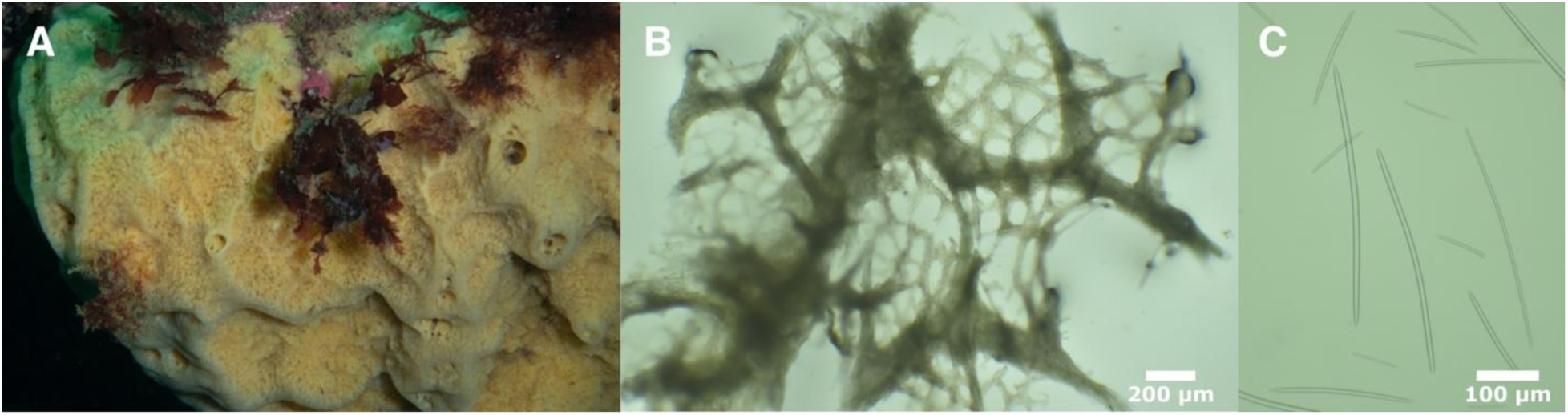
*Halichondria akesa*. A: field photos, B: tangential section showing ectosomal skeleton and subectosomal spicule tracts, C: oxeas. All photos from the holotype.

**Figure 9.**
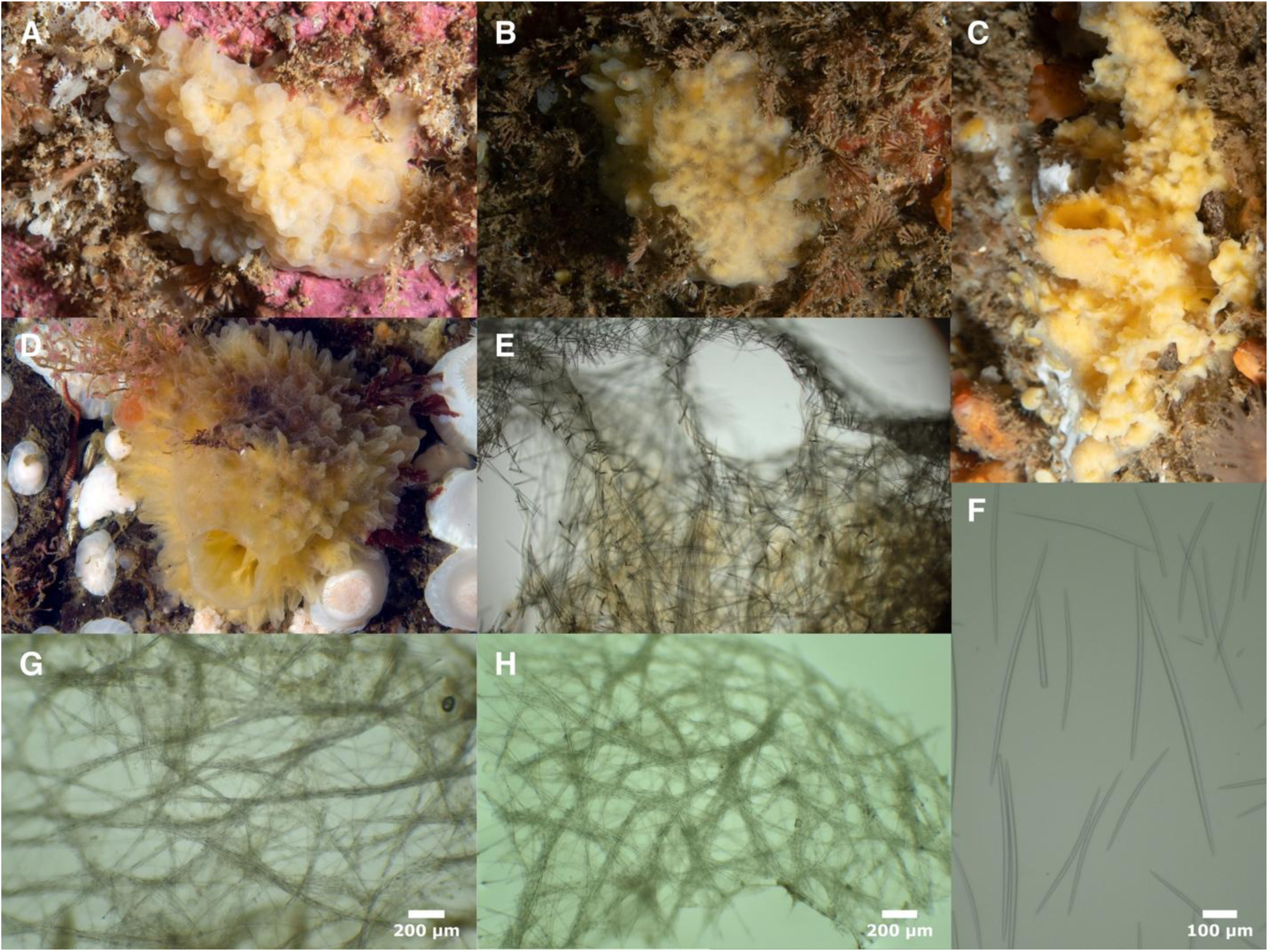
*Halichondria sitiens*. A (ESS44), B (ESS132), C (ESS153), D (RBC13, photo by Neil McDonald). E: cross-section showing choanosomal skeleton (ESS153), F: oxeas (RBC8). Tangential ectosomal skeletons: G (ESS130), H (ESS44).

## Spicules

Nearly exclusively oxeas, though rare styles can be found. Oxeas have the typical halichondroid shape: gently curved or with a slight central bend, thickest in center and tapering gently to sharp points at both ends. Spicules have a large range in lengths in both the ectosome and the choanosome, but averaged 9% longer in the choanosome. Ectosomal spicules 110–256–438 x 4–6–10 μm, n=44; choanosomal spicules 98–281–459 x 2–6–9 μm, n=44; all spicules together 98–268–459 x 2–6–10 μm, n=88. Spicules likely to be indistinguishable from *Ha. panicea*, though the minimum length was very slightly smaller than seen in any examined *Ha. panicea*.

## Distribution and habitat

Currently known from three samples, all from the far North Atlantic / Arctic: Eastern Shore Islands, Nova Scotia, Canada; Skútan, Northern Iceland; and Svalbard, Norway. Habitat preferences are likely similar to *H. panicea* – though all three samples were collected from the subtidal by divers, so it is possible this species does not occur in the intertidal.

## Remarks

Though we were only able to examine one sample of this new species, the published mitochondrial genome (MH756604) of another sample from Iceland is nearly identical. When the entire mitochondrial genome is aligned (coding and non-coding regions), these two samples differ at only 11 of 12,851 aligned sites (0.09%). In contrast, the average pairwise differences (Dxy) between the two *Ha. akesa* sp. nov. mitochondria and 9 sequenced *Ha. panicea* mitochondria is 0.8%, leading to an Fst of 0.745 across the mitochondrial genome between species. Furthermore, the ribosomal phylogeny and the phylogeny of concatenated mitochondrial genes confidently place true *Ha. panicea* sequences closer to *Ha. sitiens* than to *Ha. akesa* sp. nov. We therefore feel confident in describing these samples as a new species that is sympatric with *Ha. panicea* across the far North Atlantic. The sample from Nova Scotia was collected at the same time and place as three other samples that were found to be *Ha. panicea*, so these species are sympatric even at the scale of individual sites.

Assignment of the Svalbard sample to this species is based on only 610 bp of the cox1 gene (boldsystems.org sample ROPOR090-18). This sequence is, however, identical to the other *Ha. akesa* sp. nov. samples, and has several SNPs that differentiate it from *Ha. panicea* and other similar species.

Halichondria sitiens (Schmidt, 1870)

Figure 9

Material examined

ESS130, Faulker’s Shoal, Nova Scotia, (44.60733, -62.96168), 30 m, 2021-08-12; ESS132, Faulker’s Shoal, Nova Scotia, (44.60733, -62.96168), 30 m, 2021-08-12; ESS153, N. of Bull Rock, Nova Scotia, (44.65375, -62.90491), 30 m, 2021-08-14; ESS44, Bull Shoal, Nova Scotia, (44.63589, -62.89906), 28 m, 2021-07-21; RBC11, Knight Inlet, British Columbia, (50.68577, - 125.99457), 9 m, 2009-11-08; RBC13, Kuldekduma Island; Weynton Passage, British Columbia, depth not known, 2019-04-12; RBC8, Knight Inlet, British Columbia, (50.68500, -126.03667), 15-20 m, 2011-03-29.

## Diagnosis

Yellow, white, green, or slightly blueish *Halichondria* with numerous papillae decorating the surface. Scattered oscula usually visible, but often not at the ends of papillae. Tangential ectosomal skeleton developed as a web of meandering spicule tracts and individual spicules. Choanosomal skeleton consists of multispicular tracts and many spicules in confusion. Spicules oxeas, 130–1000 μm long, with averages per sample of 440–630 μm in length and 9–10 μm in width; usually shorter in the ectosome than the choanosome. Additional DNA characters that may be useful in identification are as follows: COX1. 754:A, 952:A. N=6. These characters differentiate *Ha. sitiens* from all other sequenced *Halichondria* except *Ha. panicea,* which is diagnosed relative to *Ha. sitiens* based on morphological characters such as spicule length and surface papillae.

28S. 2301:G. N=2. ND1. 97:C. N=5.

## Morphology

Thinly to thickly encrusting or growing as a semi-globular cushion. Surface is often replete with cone-shaped papillae that sometimes branch, but these are less apparent in small sponges. A few scattered oscula are usually visible, often large and prominent and sometimes atop oscular chimneys, but not found at the ends of the papillae. Color generally a pale, translucent yellow but surface can appear nearly white, and field photos of sponges that are likely this species have also documented pale green and slightly bluish individuals. White in ethanol.

## Skeleton

Tangential ectosomal skeleton is well-developed, but less dense and regularly-patterned than observed in *Ha. panicea*. Instead, the ectosomal skeleton is comprised of a mesh of meandering multispicular tracts and individual spicules, with spaces between them variable in size and shape (similar to species in the Bowerbanki group). Choanosomal skeleton typically halichondroid, with ill-defined spicule tracts and many spicules in confusion. No apparent spongin.

## Spicules

Exclusively oxeas with typical halichondroid shape: evenly curved or with a slight central bend, thickest in the center and tapering gently to sharp points at both ends. Spicules have a large range in lengths in both the ectosome and the choanosome, but average 14% longer in the choanosome. The difference between ectosome and choanosome is quite variable, with up to 30% longer or 5% shorter spicules in the choanosome. Dimensions are given for individual samples in table 2. When all samples are combined, spicules measure 138–531–1000 x 4–12–25 μm, n=616. Average values varied considerably across samples, with notably longer spicules seen in Pacific Canada (means 552–628 μm) than Atlantic Canada (means 448–531 μm).

**Table 2.** Spicule lengths of *Halichondria sitiens*. Values given as min–mean–max in μm.

| Specimen | Ectosome | Choanosome |
| --- | --- | --- |
| Pacific Ocean |  |  |
| RBC13 | 318–533–868 n=47 | 297–724–1000 n=47 |
| RBC11 | 335–582–892 n=40 | 332–648–964 n=40 |
| RBC8 | 308–505–844 n=41 | 295–597–893 n=43 |
| Atlantic Ocean |  |  |
| ESS130 | 312–498–623 n=53 | 325–455–619 n=59 |
| ESS153 | 270–466–625 n=40 | 302–518–698 n=40 |
| ESS44 | 138–398–680 n=43 | 332–497–659 n=43 |
| ESS132 | 258–468–728 n=40 | 370–563–723 n=40 |

## Distribution and habitat

Known from the Arctic, North Atlantic, and North Pacific, including Greenland, Svalbard, the White Sea, Alaska, and both coasts of Canada. Samples examined here were from British Columbia in the Pacific and Nova Scotia in the Atlantic, and these are the only samples with DNA data available. Previously described as occurring to 100 m depth (Erpenbeck and van Soest, 2002), but the samples examined here were all collected in the shallow subtidal by divers.

## Remarks

It is remarkable that *Ha. sitiens* is the closest relative of *Ha. panicea*, of all the species with DNA data available. Many of the species described here are morphologically indistinguishable from *Ha. panicea*, while *Ha. sitiens* has longer spicules and surface papillae that distinguish it. The presence of surface papillae has been used as a genus level trait in the past, later downgraded to the subgenus *Eumastia.* Given the close relationship between *Ha. panicea* (type species of subgenus *Halichondria*) and *Ha. sitiens* (type species of subgenus *Eumastia*), we recommend eliminating the subgenera.

In contrast to *Ha. panicea*, where fixed genetic differences were seen between Atlantic and Pacific populations, *Ha. sitiens* has nearly no genetic variation within or between populations at any of the sequenced loci.

Halichondria hygeia sp. nov.

Figure 10

Material examined

Holotype: TLT645 (CASIZ 245213), Santa Barbara Harbor, California, (34.40559, -119.68964), floating dock, 2020-02-16. Other samples: BHAK-03597 (UF 4157), Pruth Bay dock, Calvert Island, British Columbia, (51.65500, -128.12900), floating dock, 2017-08-08; FH12x108 (YPM 111876), Roche Harbor, San Juan Island, Washington, (48.60970, -123.15436), floating dock, 2012-05-23; FH12x114 (YPM 111877), Roche Harbor, San Juan Island, Washington, (48.60970, - 123.15436), floating dock, 2012-05-23; TLT1076 (SBMNH 718624), Santa Barbara Harbor, California, (34.40559, -119.68964), floating dock, 2021-06-01; TLT1417 (UCSB-IZC00069011), Santa Barbara Harbor, California, (34.40559, -119.68964), floating dock, 2023-09-01; TLT308 (UCSB-IZC00069012), Monterey Marina, California, (36.60797, -121.89288), floating dock, 2019-11-22; TLT504 (CASIZ 245212), Point Pinos, California, (36.63447, -121.93923), intertidal, 2019-11-24; TLT706 (SBMNH 718625), Oceanside Harbor, California, (33.20623, -117.39208), floating dock, 2021-01-09; TLT724 (CASIZ 245214), Ventura Harbor, California, (34.24801, -119.26550), floating dock, 2020-12-14; TLT742 (SBMNH 718626), Dana Point Harbor, California, (33.46090, - 117.69156), floating dock, 2021-01-09; TLT940 (UCSB-IZC00069013), Lazy Days, San Diego, California, (32.69415, -117.27110), 12-25 m, 2021-05-15; WMS19x5 (YPM 111878), West Meadow Beach, Stony Brook, New York, (40.92673, -73.14807), intertidal, 2019-05-19; WMS19x6 (YPM 111879), West Meadow Beach, Stony Brook, New York, (40.92673, -73.14807), intertidal, 2019-05-19; WMS19x7 (YPM 111880), West Meadow Beach, Stony Brook, New York, (40.92673, -73.14807), intertidal, 2019-05-19; WMS21x10 (YPM 111881), West Meadow Beach, Stony Brook, New York, (40.92673, -73.14807), intertidal, 2021-05-26; WMS21x19 (YPM 111882), West Meadow Beach, Stony Brook, New York, (40.92673, -73.14807), intertidal, 2021- 05-26; WMS21x6 (YPM 111883), West Meadow Beach, Stony Brook, New York, (40.92673, - 73.14807), intertidal, 2021-05-26; WMS21x7 (YPM 111884), West Meadow Beach, Stony Brook, New York, (40.92673, -73.14807), intertidal, 2021-05-26; WMS21x8 (YPM 111885), West Meadow Beach, Stony Brook, New York, (40.92673, -73.14807), intertidal, 2021-05-26; WMS21x9 (YPM 111886), West Meadow Beach, Stony Brook, New York, (40.92673, -73.14807), intertidal, 2021-05-26; WMS23x10 (YPM 111887), West Meadow Beach, Stony Brook, New York, (40.92673, -73.14807), intertidal, 2023-06-07; WMS23x9 (YPM 111888), West Meadow Beach, Stony Brook, New York, (40.92673, -73.14807), intertidal, 2023-06-07.

## Etymology

Name inspired by the Greek goddess Hygeia, sister of Panakeia.

## Diagnosis

Yellow *Halichondria* with well-developed tangential ectosomal skeleton manifested as a dense mat of spicules or a mesh of meandering multispicular tracts and individual spicules.

Choanosomal skeleton confused, sometimes accompanied by multispicular tracts. Spicules oxeas, 170–430 μm in length, with averages per sample of 250–350 μm in length and 5–10 μm in width. On average, no difference in length is seen between the ectosome than the choanosome. Diagnosis relative to similar species requires DNA characters, as follows: COX1. No unique derived states.

28S. 659:G, 670-671:GG, 675:A. N=24.

ND1. 877-880:AACC. State 877:A is unique compared to all close relatives, but homoplastic with *Halichondria loma.* The full sequence 877-880:AACC is not shared with any other sequenced *Halichondria*. N=14.

## Morphology

Thinly to thickly encrusting, sometimes with digitate tendrils. Oscula vary from not apparent, to slightly volcano-shaped with raised rims, to atop oscular chimneys. Color varies from pale yellow to golden; not seen in green. See Figure 10 for examples.

**Figure 10.**
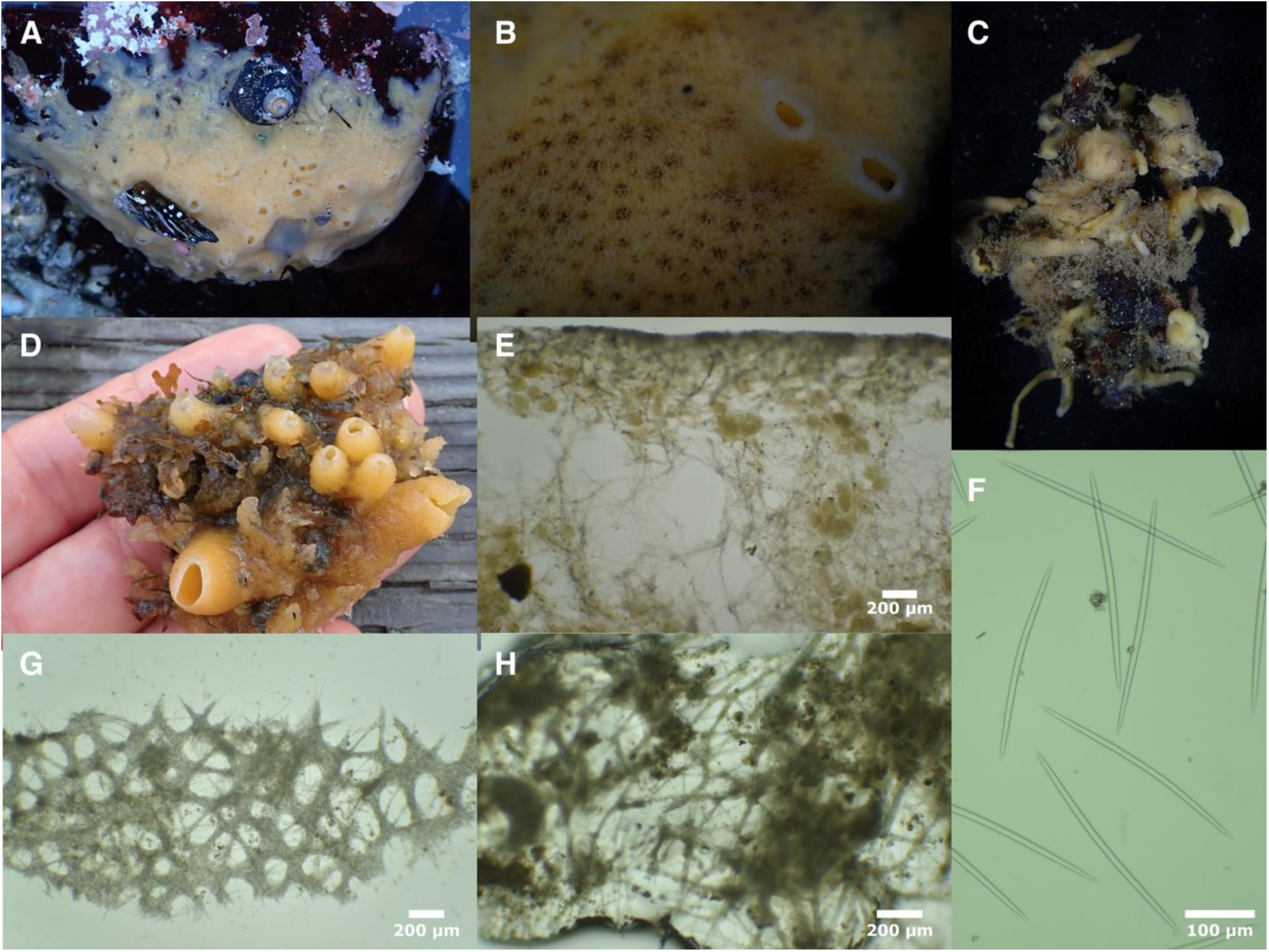
*Halichondria hygeia*. A & B (TLT504), C (TLT645), D (TLT706). E: cross-section showing choanosomal skeleton at sponge surface (TLT504), F: spicules (FH12x108). Tangential ectosomal skeletons: G (TLT742), H (TLT645).

## Skeleton

Tangential ectosomal skeleton can be sieve-like, with a dense mat of spicules surrounding oval-shaped open areas (Figure 10G), or more lace-like, with a web of spicule tracts and individual spicules delineating more irregularly shaped open areas (Figure 10H). This ectosomal skeleton is supported by poorly developed spicule tracts that delineate large subectosomal spaces, with the choanosomal skeleton becoming more confused towards the interior. No apparent spongin.

## Spicules

Exclusively oxeas with a typical halichondroid shape: evenly curved or with a slight central bend, thickest in the center and tapering to sharp points at both ends. Spicules have a large range in lengths, though slightly less range than seen in *Ha. panicea*, and spicules in the ectosome and the choanosome average the same length. Dimensions for five samples where ectosome and choanosome were measured separately are given in table 3. When spicules from all seven measured samples are combined, spicules measure 175–276–424 x 3–7–13 μm, n=561 for length, n=405 for width. Average values varied considerably across samples, with mean lengths 254–346 μm, and mean widths 5–10 μm.

**Table 3.** Spicule lengths of *Halichondria hygeia*. Values given as min–mean–max in μm.

| Specimen | Ectosome | Choanosome |
| --- | --- | --- |
| Pacific Ocean |  |  |
| FH12x108 | 284–343–386 n=40 | 236–349–424 n=40 |
| TLT645 | 194–259–313 n=40 | 216–269–312 n=40 |
| TLT742 | 190–267–301 n=40 | 193–259–319 n=43 |
| TLT706 | 188–284–343 n=42 | 232–279–328 n=43 |
| Atlantic Ocean |  |  |
| WMS21x9 | 246–283–336 n=40 | 204–256–302 n=44 |

## Distribution and habitat

In the Northeast Pacific, this species was found from Calvert Island, British Columbia to Oceanside, Southern California. In contrast to *Ha. panicea*, *Ha. hygeia* sp. nov. was found primarily on floating docks in sheltered marinas, but it was not limited to this habitat. Of 12 samples collected in the Northeast Pacific, 10 were on floating docks but one was in found in a wave-swept tide pool at Point Pinos (Monterey Bay, Central California), and one was found in subtidal rocky reef off Point Loma (extreme Southern California, collected between 12 and 25 m depth). The species is also found in the North Atlantic, as it was common on intertidal cobbles at West Meadow Beach, Stony Brook, Long Island, New York. Publicly available sequences of cox1 may indicate that the species is also found in Connecticut and Portugal (Genbank accessions MN448237, MZ487266, figure S1). These sequences are consistent with either *Ha. hygeia* sp. nov. or *Ha. californiana* sp. nov.; as *Ha. californiana* sp. nov. is thus far known only from California, these sequences are likely to be *Ha. hygeia* sp. nov., but confirmation would require sequencing 28S or ND1.

## Remarks

Two pieces of evidence suggest that *Ha. hygeia* sp. nov. has experienced a recent human-assisted range expansion. First, artificial substrates like floating docks — where this species is primarily found — are known to harbor many introduced species (Bulleri and Chapman, 2010; Glasby et al., 2007; Ruiz et al., 2000; Simkanin et al., 2012). Second, samples from New York and California are nearly identical across the mitochondrial genes and the ribosomal DNA (Figure 1). Other halichondriids have been shown to have limited larval dispersal (Maldonado, 2006; Xue et al., 2009), so together, these observations make it likely that the species has been moved around by human activity.

*Halichondria hygeia* sp. nov. and *Ha. panicea* are likely to be indistinguishable with respect to spicule and skeletal traits, but some differences between them are apparent. These include habitat (*Ha. hygeia* sp. nov. is common on floating docks in marinas, while *Ha. panicea* is common in the wave-exposed intertidal), southern range limit (*Ha. hygeia* sp. nov. was found in Southern California, while *Ha. panicea* was not found in Southern California), and color (*Ha. hygeia* sp. nov. is yellow while *Ha. panicea* can be yellow or green).

Halichondria californiana sp. nov.

Figure 11

Material examined

Holotype: TLT937 (CASIZ 245207), Six Fathoms, San Diego, California, (32.71000, -117.26860), 9- 18 m, 2021-05-15. Other samples: TLT1010 (UCSB-IZC00069005), Arroyo Quemado Reef, California, (34.46775, -120.11905), 7-11 m, 2021-07-22; TLT1093 (UCSB-IZC00069006), Arroyo Quemado Reef, California, (34.46775, -120.11905), 7-11 m, 2021-07-22; TLT160 (CASIZ 245204), Isla Vista Reef, California, (34.40278, -119.85755), 9-12 m, 2019-08-01; TLT256 (UCSB- IZC00069007), Isla Vista Reef, California, (34.40278, -119.85755), 9-12 m, 2019-08-01; TLT417 (CASIZ 245205), Coal Oil Point, California, (34.40450, -119.87890), 3-8 m, 2019-10-25; TLT428 (CASIZ 245206), Elwood Reef, California, (34.41775, -119.90150), 9-15 m, 2019-10-23; TLT812 (SBMNH 718620), Six Fathoms, San Diego, California, (32.71000, -117.26860), 9-18 m, 2021-05-15; TLT860 (SBMNH 718621), Cave Landings, California, (35.17535, -120.72240), intertidal, 2021-02-06; TLT921 (SBMNH 718622), Point Fermin, California, (33.70664, -118.28595), intertidal, 2021-03-10; TLT928 (UCSB-IZC00069008), Point Fermin, California, (33.70664, - 118.28595), intertidal, 2021-03-10; TLT934 (UCSB-IZC00069009), Six Fathoms, San Diego, California, (32.71000, -117.26860), 9-18 m, 2021-05-15.

## Etymology

Named for the state of California.

## Diagnosis

Yellow *Halichondria* with tangential ectosomal skeleton developed as a sieve-like mat of spicules or a lace-like network of spicule tracts. Choanosomal skeleton consists of poorly delineated multispicular tracts and spicules in confusion. Spicules oxeas, ranging from 90–290 μm in length, with averages per sample of 130–230 μm long and 3–7 μm wide. On average, no difference in length is seen between the ectosome than the choanosome. Diagnosis relative to similar species requires DNA characters, as follows:

COX1. No unique derived states.

28S. 1011:A, 1037:T, 1319:C. N=9–11 depending on position.

ND1. 214-217:AAAT. State 217:T is unique compared to all close relatives, but homoplastic with members of the Okadai group (Figure 1). The full sequence 214–217:AAAT is not shared with any other sequenced *Halichondria*. N=10.

## Morphology

Encrusting yellow sponges; oscula often small and flush with sponge surface, but can also occur on oscular chimneys. Color varies from pale, translucent yellow to golden, sometimes with red blotches; not seen in green. Beige in ethanol. See Figure 11 for examples.

**Figure 11.**
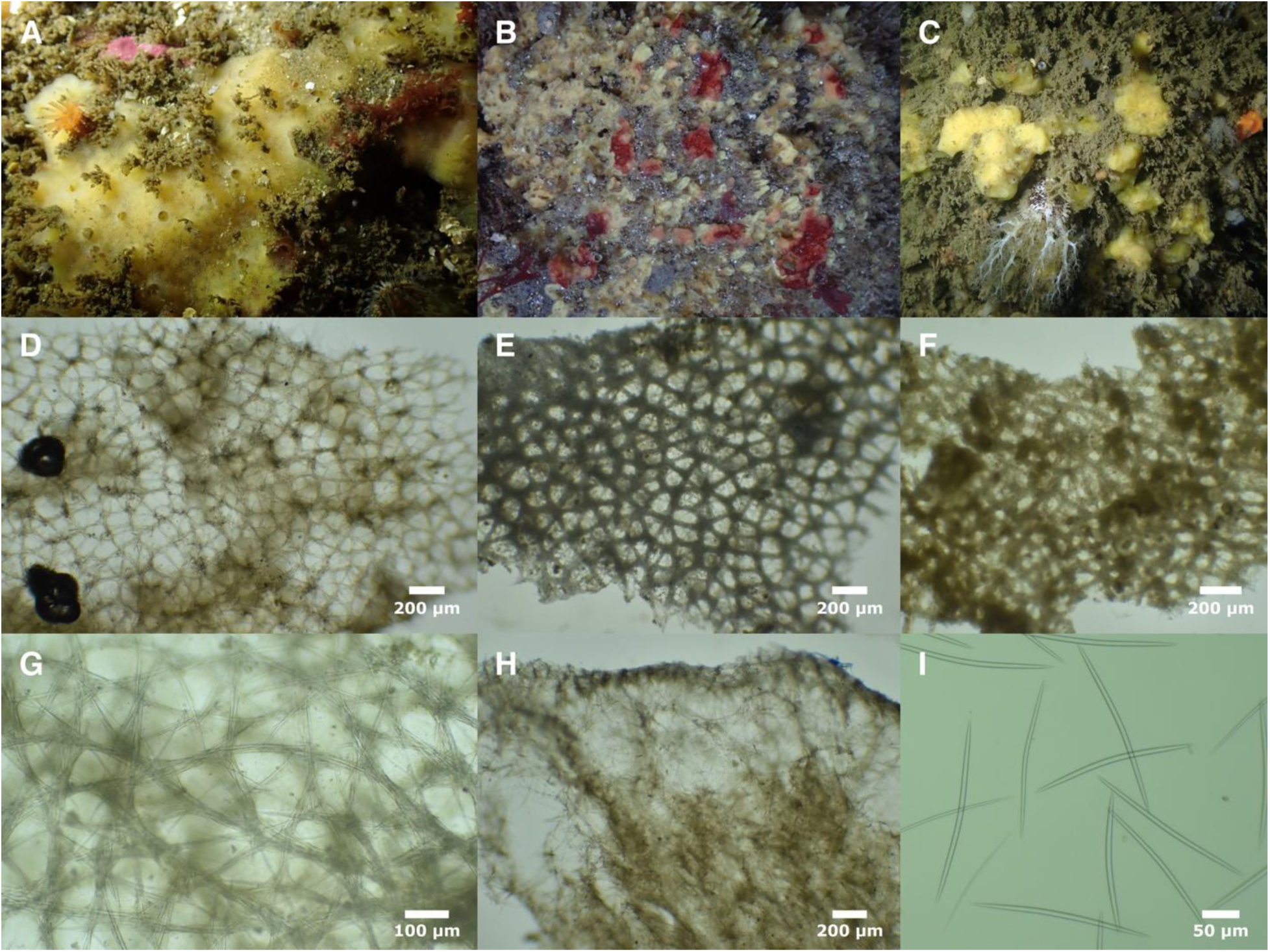
*Halichondria californiana*. Field photos: A (TLT937), B (TLT880), C (TLT1093). Ectosomal skeletons: D (TLT160), E (TLT937), F (TLT1010), G (TLT860). H: cross-section showing choanosomal skeleton at sponge surface (TLT160), I: spicules (TLT860).

## Skeleton

Tangential ectosomal skeleton can be sieve-like, with a dense mat of spicules, or more lace-like, with a web of spicule tracts and single spicules. Open areas are generally oval or polygonal and relatively regular in size. Choanosomal skeleton with poorly developed spicules tracts and many spicules in confusion. No apparent spongin.

## Spicules

Nearly exclusively oxeas, though rarely a spicule can be found that is modified into a style. Spicules have a typical halichondroid shape: evenly curved or with a slight central bend, thickest in the center and tapering gently to sharp points at both ends. Spicules have a large range in lengths, though slightly less range than seen in *Ha. panicea*, and spicules in the ectosome and the choanosome average the same length. Dimensions for the six samples where ectosome and choanosome were measured separately are given in table 4. When spicules from 10 measured samples are combined, spicules measure 99–194–286 x 2–5–9 μm, n=813 for length, n=733 for width. Average values varied considerably across samples, with means of 134–226 μm long and 3–7 μm wide.

**Table 4.** Spicule lengths of *Halichondria californiana*. Values given as min–mean–max in μm.

| Specimen | Ectosome | Choanosome |
| --- | --- | --- |
| TLT937 | 149–207–239 n=40 | 177–215–259 n=40 |
| TLT1010 | 113–136–172 n=51 | 99–132–161 n=51 |
| TLT921 | 121–177–199 n=40 | 145–180–208 n=40 |
| TLT256 | 107–146–164 n=40 | 122–155–190 n=41 |
| TLT860 | 187–235–274 n=40 | 190–236–286 n=59 |
| TLT428 | 167–213–258 n=40 | 166–218–266 n=40 |

The maximum length of 286 μm is notably shorter than some similar species, such as *Ha. hygeia* sp. nov. (424 μm), *Ha. panicea* (499 μm), and *Ha. cascadia* sp. nov. (444 μm) — though *Ha. columbiana* sp. nov. is similar (266 μm). Spicule size is insufficient to differentially diagnose all samples, however. For example, some individual *Ha. californiana* sp. nov. samples had average and maximum lengths exceeding some *Ha. hygiea* sp. nov. samples, and some *Ha. californiana* sp. nov. had average lengths exceeding some *Ha. panicea*.

## Distribution and habitat

Known only from California, from San Luis Obispo Bay in Central California to Point Loma in Southern California. Nearly allopatric with *Ha. panicea*: the two species were found growing side-by-side in a tidal pool near Avila Beach, California, but this was the southernmost location where *Ha. panicea* was found and the northernmost location for *Ha. californiana* sp. nov.. Like *Ha. panicea,* this species was found growing on rock or algae, in natural subtidal reefs and exposed intertidal areas, and not found on human structures in sheltered marinas. The deepest samples were found between 9 and 18 m.

## Remarks

In contrast to its closest known relative *Ha. hygeia* sp. nov., this species was found only in natural habitats, and not on human structures. It is also different from *Ha. hygeia* sp. nov. in that it was only found in California and thus could be endemic to the Northeast Pacific.

However, these two species cannot be differentiated at the commonly sequenced regions of cox1, so publicly available sequences from Connecticut and Portugal could be this species (Genbank accessions MN448237, MZ487266). The nudibranch *Doris montereyensis* was seen feeding on this species in multiple locations.

Halichondria cascadia sp. nov.

## Material examined

Holotype: FH12x92 (CASIZ 245208, YPM 111874), Eagle Cove, San Juan Island, Washington, (48.46128, -123.03155), intertidal, 2012-05-22. Additional sample: FH12x93 (YPM 111875), Eagle Cove, San Juan Island, Washington, (48.46128, -123.03155), intertidal, 2012-05-22.

## Etymology

Name inspired by the term "Cascadia region" which is sometimes used to refer to an area including Oregon, Washington, and British Columbia.

## Diagnosis

*Halichondria* with a tangential ectosomal skeleton developed as a mesh of meandering multispicular tracts and individual spicules. Choanosomal skeleton consists of spicules and spicule bundles in confusion. Spicules oxeas, 160–450 μm long, with averages per sample of 270–320 μm long and 8-9 μm wide. Infrequent styles and rare strongyles can be observed.

Some samples average shorter spicules in the ectosome than the choanosome. Diagnosis based primarily on DNA characters, as follows: COX1. 949:A. N=3. 28S. 2061:T. N=2. ND1. 847:A. N=2.

## Morphology

Known only from two small samples, so little is known about the morphology of this species. Preserved samples are white; color alive is yellow. Both samples have oscular chimneys approximately 1 cm in height.

## Skeleton

Tangential ectosomal skeleton a web of spicule tracts and individual spicules, with open areas irregular in size and shape (Figure 12B). Choanosomal skeleton of spicules and spicule bundles in confusion. No apparent spongin.

**Figure 12.**
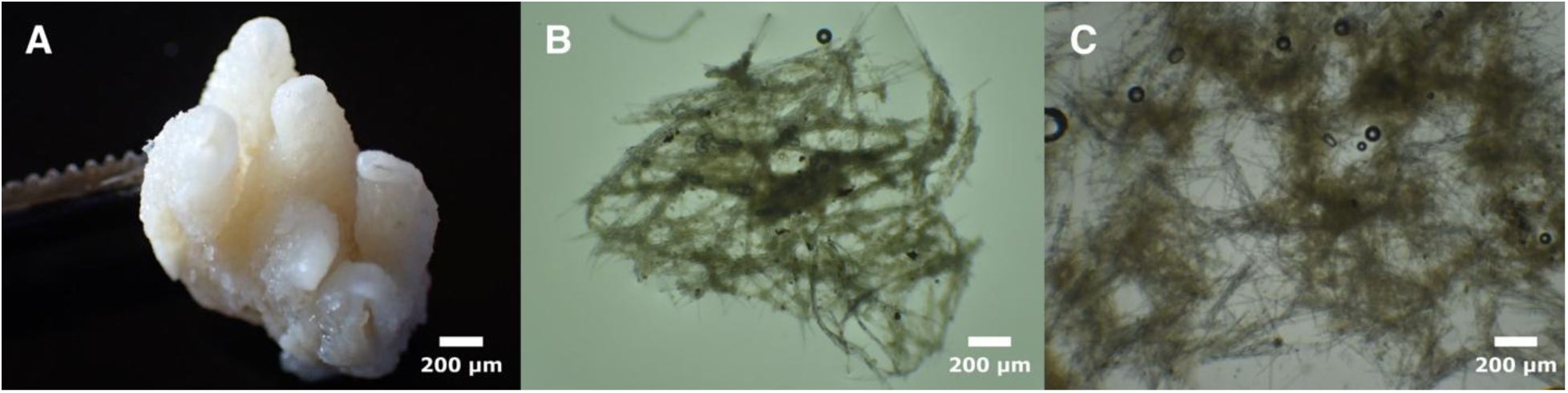
*Halichondria cascadia*. A: sample post-preservation (FH12x92), B: tangential ectosomal skeleton (FH12x93), C: choanosomal skeleton (FH12x92).

## Spicules

Oxeas with a typical halichondroid shape: evenly curved or with a slight central bend, thickest in the center and tapering to sharp points at both ends. Infrequent styles and rare strongyles can be observed. Spicules have a large range in lengths, with one sample having significantly shorter spicules in the ectosome while the other sample has no difference in length between ectosome and choanosome. When spicules from both samples are combined, spicules measure 162–294–444 x 3–8–12 μm, n=169. Table 5 shows dimensions for samples and domains separately.

**Table 5.** Spicule lengths of *Halichondria cascadia*. Values given as min–mean–max in μm.

| Specimen | Ectosome | Choanosome |
| --- | --- | --- |
| FH12x92 | 178–267–353 n=41 | 162–273–373 n=46 |
| FH12x93 | 199–287–280 n=42 | 260–353–444 n=40 |

## Distribution and habitat

Samples examined here were both collected from rocky intertidal habitats at Eagle Cove, San Juan Island, Washington. Publicly available sequence data suggests a third sample was found at Carl Moses Harbor, Captains Bay, Unalaska Island (Genbank accession MZ580608). Though only 658 bp of cox1 is available for this sample, it is identical to sample FH12x93.

## Remarks

The two samples examined here are small fragments. Though they were collected at the same location, they are genetically different from one another: they differ at 1.17% of bases in the nuclear ribosomal locus and at 0.6% of bases in the protein-coding mitochondrial genome. They form a well-supported clade in the phylogeny, with 98.5% bootstrap support, and are the sister group to the other three species in the Hygeia group (*Ha. hygeia* sp. nov.*, Ha. californiana* sp. nov.*, and Ha. columbiana* sp. nov.). They were collected at the same time and place as *Ha. panicea*, and *Ha. hygiea* sp. nov. was collected at a marina on the same island the following day, so this species is sympatric with several closely related *Halichondria.* Despite the limited sampling of this species, the strong support for *Ha. hygiea* sp. nov. and *Ha. californiana* sp. nov. as different species compels us to formally describe these samples in the hope that more can be learned about this enigmatic group.

Halichondria columbiana sp. nov.

Figure 13

Material examined

Holotype: BHAK-09466 (UF 4408), Port Reef, Hakai Pass, British Columbia, (51.697, -128.106), 28 m, 2019-05-28. Additional sample: BHAK-09479, Whidbey Point, Fitz Hugh Sound, British Columbia, (51.71700, -127.89200), 19 m, 2019-05-29.

## Etymology

Named for the British Columbia region of Canada.

## Diagnosis

*Halichondria* with tangential ectosomal skeleton developed as a chaotic mesh of meandering multispicular tracts and individual spicules. Choanosomal skeleton confused. Spicules oxeas, 150–270 μm long, with means per sample of 220–240 μm long and 8–9 μm wide. Diagnosis based primarily on DNA characters, as follows:

COX1. 973:C, 1012:C, 1102:G, 1207:A. N=2.

28S. No unique derived states.

ND1. 79:C, 88:C, 172:T, 340:G, 352:C, 733:G, 811:G. N=2.

## Morphology

Known only from two small samples, little is known about the morphology of this species, but they appear similar to encrusting *Ha. panicea*. Samples were yellow alive and are white in ethanol.

## Skeleton

Tangential ectosomal skeleton is a web of spicule tracts and individual spicules (Figure 13C). Choanosomal skeleton difficult to characterize in such small samples, but appears to be confused, as seen in all closely related species. No apparent spongin.

**Figure 13.**
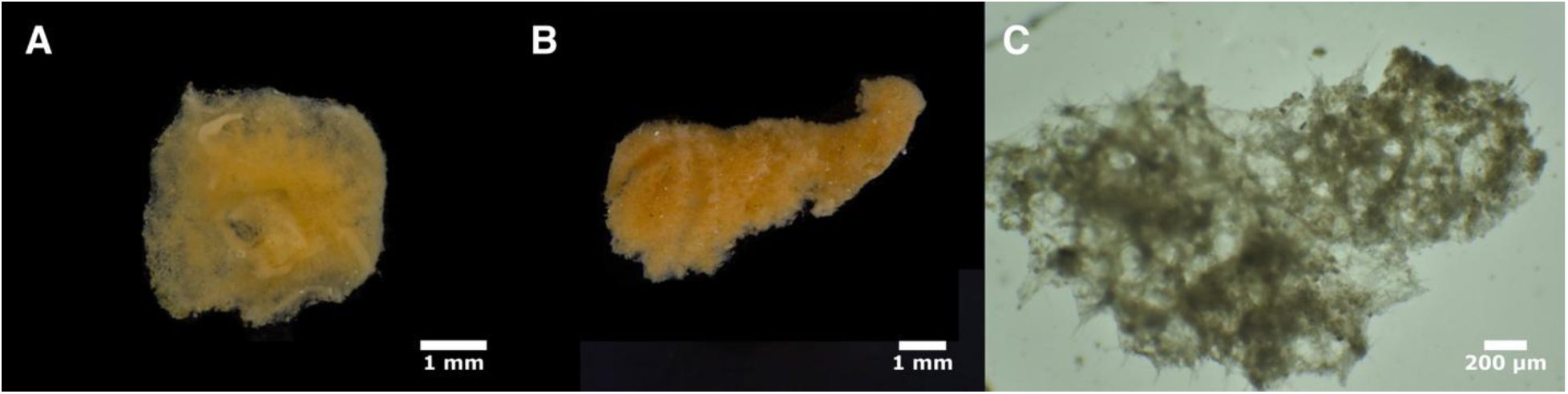
*Halichondria columbiana*. Photos of sponges after collection, before preservation: A (BHAK- 9466), B (BHAK-9479). C: tangential ectosomal skeleton (BHAK-9466).

## Spicules

Exclusively oxeas with a typical halichondroid shape: evenly curved or with a slight central bend, thickest in the center and tapering to sharp points at both ends. When spicules from both samples are combined, spicules measure 149–228–266 x 4–8–11 μm, n=151. Spicules from the ectosome and choanosome were measured separately for one sample and the sizes did not differ by compartment (the ectosome could not be located in the other small sample).

## Distribution and habitat

The two known samples were collected 15 km apart in British Columbia. Both were subtidal, between 19 and 28 m depth.

## Remarks

The two samples examined here are very small fragments. The samples are identical at the ribosomal locus and only slightly different at mitochondrial loci (sequence differences of 0.4% and 0.2% at ND1 and cox1, respectively). Contrastingly, they are genetically distinct from the other species in the Hygeia group: when compared to *Ha. hygeia* sp. nov., average pairwise differences are 1.0% (Fst=0.9) at the ribosomal locus and 1.2% (Fst=0.84) at cox1. Despite the limited sampling of this species, the strong support for *Ha. hygiea* sp. nov. and *Ha. californiana* sp. nov. as different species compels us to formally describe these samples in the hope that more can be learned about this enigmatic group.

Halichondria dokdoensis Kang, Kim & Kim, 2022

Figure 14

Material examined

TLT386, Oil Platform C, Santa Barbara, California, (34.33293, -119.63173), 21 m, 2019-08-05.

## Diagnosis

Yellow, reddish-purple, or orange *Halichondria* with tangential ectosomal skeleton developed as a lace-like network of spicule tracts surrounding oval-shaped open areas. Choanosomal skeleton with multispicular tracts supporting the ectosome and delineating large subectosomal spaces, with an increasingly confused skeleton towards the interior. Spicules oxeas, 130–310 μm long and 1–10 μm wide; slightly shorter in the ectosome than the choanosome. Diagnosis relative to similar species requires DNA characters, as follows:

COX1. 100:C, 247:G, 361:G, 844:C, 892:T. N=2-3 depending on position. 28S. 162:T, 195:C, 260:G, 584:C, 614:T, 671:T, 1660:G. N=1.

ND1. 286:C, 322:A, 346:G, 403:T, 769:G. N=2.

## Morphology

Samples from Korea were previously described as reddish-purple to orange with prominent oscular chimneys (Kang et al., 2022). A sample from California was yellow, with oscula nearly flush with the sponge surface, so color and shape appear to be variable, as seen in many other *Halichondria* species.

## Skeleton

Well-developed tangential ectosomal skeleton with a lace-like web of spicules and spicule tracts surrounding open spaces that are oval shaped and fairly consistently sized (Figure 14B). This ectosomal skeleton is supported by multispicular tracts that delineate large subectosomal spaces. The choanosomal skeleton becomes more confused towards the interior. No apparent spongin.

**Figure 14.**
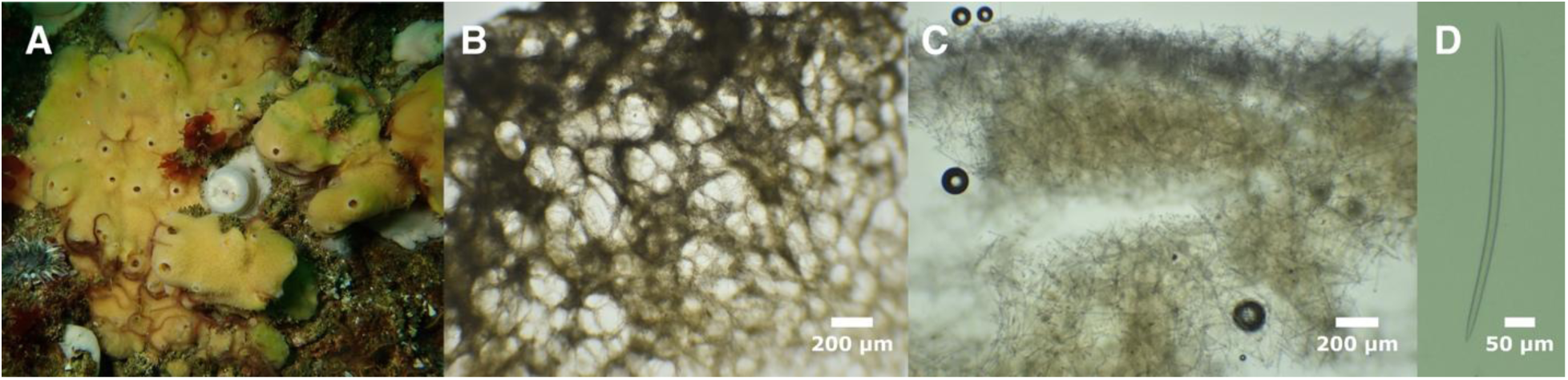
*Halichondria dokdoensis*. A: field photo, B: tangential ectosomal skeleton, C: cross-section showing choanosomal skeleton at sponge surface, D: oxea. All photos from TLT386.

## Spicules

Exclusively oxeas with a typical halichondroid shape: evenly curved or with a slight central bend, thickest in the center and tapering gently to sharp points at both ends (rare spicules occasionally found with a centrotylote swelling). The single sponge we examined had spicules 177–230–287 x 2–6–8 μm, n=146 for length, n=106 or width. When measured separately, spicules in the choanosome averaged 238 μm in length vs. 221 μm in the ectosome (p=0.001). Spicules from Korean samples previously reported as 135–310 x 1–8 μm (means and sample size not reported). Spicules were previously reported as having "size classes", with some being thin (1–2.5 μm) and some thick (5–8 μm). Of the 146 spicules we measured, only one would fall in the thin category, with 10 falling between thick and thin. The abundance of thin oxea is therefore variable and/or universally rare, perhaps because they are immature thick spicules.

## Distribution and habitat

The holotype location for this species is the Dokdo Islands, South Korea, with additional samples from mainland South Korea and Jeju Island. The sample examined here was found on an oil platform in the Santa Barbara Channel in Southern California, and a publicly available sequence matching this species was obtained near Goes, Zeeland, the Netherlands (see remarks). The California sample was subtidal (21 m), and though depths are not reported for the other samples, the Korean samples were also collected by SCUBA divers.

## Remarks

*Halichondria dokdoensis* was described based on morphological characters, but it is our view that it cannot be morphologically distinguished from *Ha. panicea*. Characters previously listed as distinguishing the species were color (but we show here that *Ha. dokdoensis* can be found in yellow) and larger spicules than *H. panicea* (but we show here that the spicules of *Ha. panicea* span a large size range encompassing that of *Ha. dokdoensis*). However, a mitochondrial genome sequence was previously published for *Ha. dokdoensis*, and we are able to use this sequence to "rescue" this previously published name (Kim et al., 2019). This sequence is nearly identical to the mitochondrial genome of our California sample: they differ by only 12 of 13,033 aligned base pairs (<0.1%). These samples are more closely related to *Hymeniacidon actites* than to any other sequenced *Halichondria*, and together these two species form a clade that is the sister to the combined Panicea and Hygeia species groups.

This species is likely to have experienced a recent human-mediated range expansion. Finding samples in both California and Korea could plausibly be due to a natural range across the North Pacific, though this is made less likely by the observation that it was only found (thus far) on an artificial structure in California. However, a publicly available sequence of cox1 indicates that the species is also likely found in the Netherlands (NLMAR324-20, boldsystems.org). Though this sequence is only 618 bp of the cox1 gene, it is identical to other *Ha. dokdoensis* and does not match any other sequenced species. This species is therefore likely to be introduced to multiple regions of the Northern Hemisphere, similar to other species described here.

Halichondria galea sp. nov.

Figure 15

Material examined

Holotype: TLT495 (CASIZ 245211), Point La Jolla, California, (32.85227, -117.27239), intertidal, 2020-02-08. Other samples: TLT496 (SBMNH 718623), Point La Jolla, California, (32.85227, - 117.27239), intertidal, 2020-02-08; TLT743 (UCSB-IZC00069010), Dana Point Harbor, California, (33.46090, -117.69156), floating dock, 2021-01-09.

## Etymology

Name inspired by the word "galleon", one of the ships of the Spanish armada in the age of sail.

## Diagnosis

Encrusting green, yellow, or brownish *Halichondria*, sometimes with a rugose surface. Tangential ectosomal skeleton is a sieve-like mat of spicules surrounding open areas that are generally oval-shaped and consistent in size. Choanosomal skeleton of multispicular tracts and abundant spicules in confusion. Spicules oxeas, 100–500 μm long, with averages per sample of 250–290 μm long and 5–6 μm wide. Spicules 10–20% longer in choanosome than ectosome, but the full range of spicules lengths is seen in both compartments. Diagnosis relative to similar species requires DNA characters, as follows:

COX1. 1069:A. N=8.

28S. 516:T, 520:G, 547:G, 1943:T, 2111:T. N=2–3 depending on position. ND1. No unique derived states.

## Morphology

Thinly to thickly encrusting with uneven thickness, leading to a lumpy, undulating, or rugose appearance. Oscula, when apparent, nearly flush with surface. Surface is a mix of yellow and green regions, or yellow, green, and brownish regions, with the interior yellow. See Figure 15 for examples.

**Figure 15.**
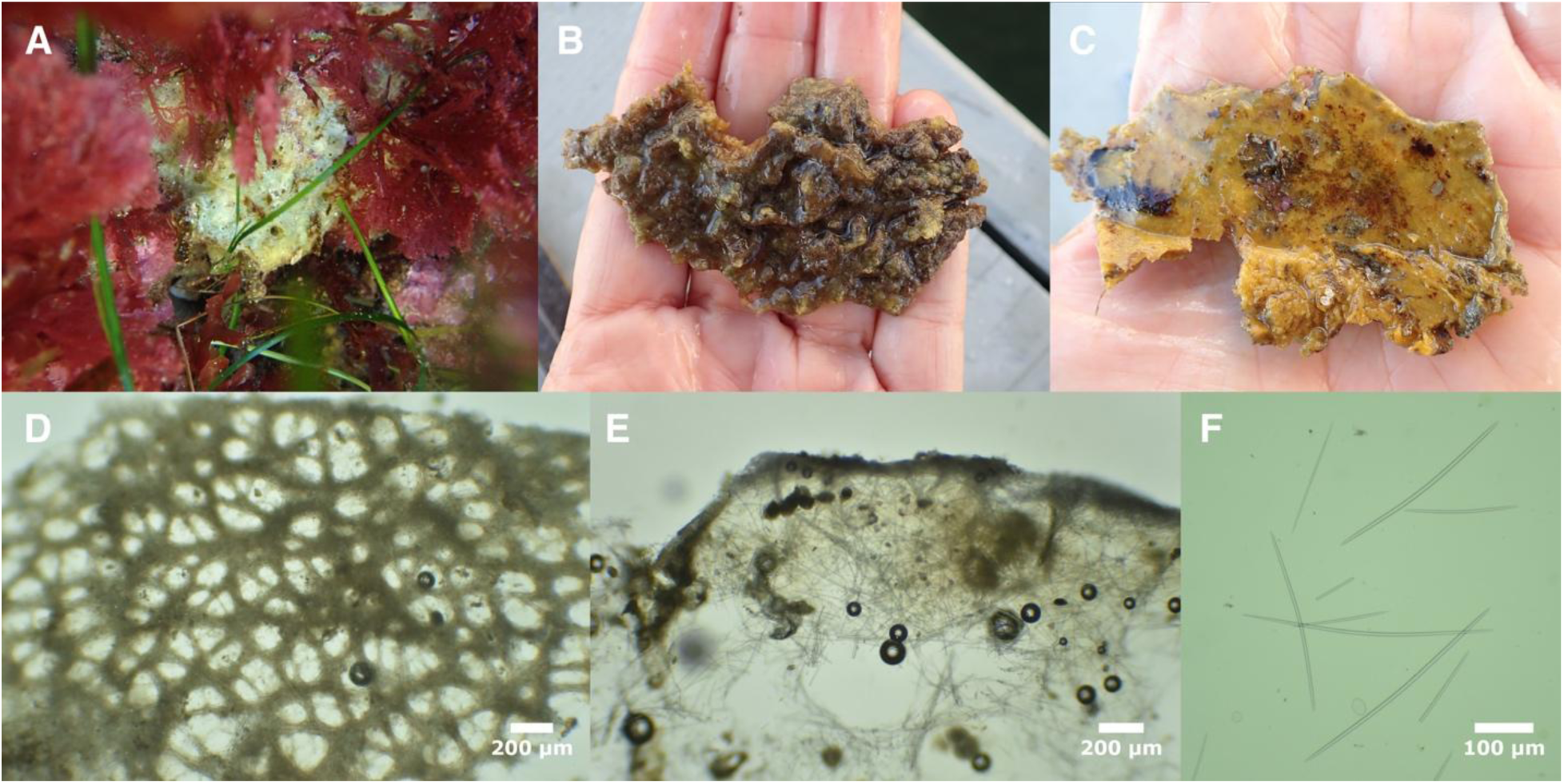
*Halichondria galea*. A (TLT495), B & C (TLT743, C shows underside that was attached to floating dock). D: tangential ectosomal skeleton (TLT496), E: cross-section showing choanosomal skeleton at sponge surface (TLT495), F: oxeas (TLT495).

## Skeleton

Tangential ectosomal skeleton a sieve-like mat of spicules surrounding oval-shaped open areas with fairly consistent size. Choanosomal skeleton with meandering multispicular tracts and many spicules in confusion. No apparent spongin.

## Spicules

Exclusively oxeas with a typical halichondroid shape: evenly curved or with a slight central bend, thickest in the center and tapering to sharp points at both ends. Like *Ha. panicea*, spicules have a very large range in lengths. Though spicules in the ectosome are 10–20% shorter than the choanosome, the full range of spicule lengths is seen in both compartments, with size distributions continuous but bimodal within each compartment. Dimensions for three samples where ectosome and choanosome were measured separately are given in table 6.

**Table 6.**
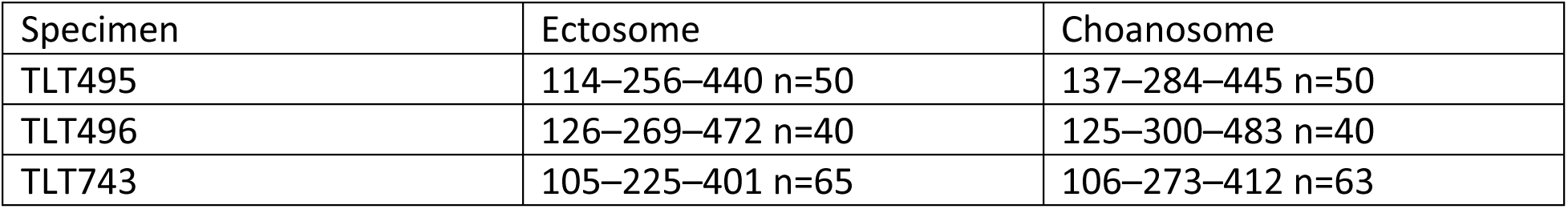
Spicule lengths of *Halichondria galea*. Values given as min–mean–max in μm.

| Specimen | Ectosome | Choanosome |
| --- | --- | --- |
| TLT495 | 114–256–440 n=50 | 137–284–445 n=50 |
| TLT496 | 126–269–472 n=40 | 125–300–483 n=40 |
| TLT743 | 105–225–401 n=65 | 106–273–412 n=63 |

When spicules from all samples are combined, spicules measure 105–265–483 x 2–5–10 μm, n=308. Average dimensions per sample were 249–270 μm long and 5–6 μm wide.

## Distribution and habitat

Known from China, Korea, and California. In the Eastern Pacific, this species is known from Orange and San Diego Counties in Southern California. We found it growing on a floating dock in Dana Point Harbor and in natural tidepools at Point La Jolla; it was also previously reported from Mission Bay, San Diego (Park et al., 2007). In the Western Pacific it has been collected from the Gulei Peninsula in China (Wang et al., 2016) and from rocky intertidal areas around the Korean Peninsula and islands of the Korean Strait (Park et al., 2007).

## Remarks

The first mitochondrial genome published for a *Halichondria* species was for a "*Halichondria sp.*" sponge sampled from Fujian Province, China (Wang et al., 2016). Sequencing of mitochondrial genes later documented this same genotype from the intertidal zone of mainland South Korea and nearby islands, and from Mission Bay, San Diego, California (Park et al., 2007). This sponge has remained undescribed, likely due to it being morphologically indistinguishable from *Ha. panicea*.

Our Southern California samples have mitochondrial genomes that are nearly identical with the published sequence from China (99.95% identical), confirming that we have recollected this same species from California. Consistent with previously published data, it is only distantly related to *Ha. panicea*, despite their morphological similarity. The clade we designate here as the Okadai species group contains *Ha. okadai* and *Ha. galea* sp. nov., which are sympatric in the Korean Strait, as well as the Caribbean species *Ha. magniconulosa* and *Ha. melanadocia,* and samples we collected from Pacific Panama that are likely from an undescribed species (Figure 3).

It is probable that *Ha. galea* sp. nov. has experienced a recent human-mediated range expansion, though the evidence is less compelling than for the other species reported here. It has only been found in the North Pacific thus far, and this could indicate a natural range connected through the Bering Strait. It has been found in artificial habitats like floating docks, but most samples reported to date have been collected from rocky intertidal areas that host fewer introduced species. However, finding nearly identical mitochondrial genomes in China and California makes it less likely that those populations are found at extreme ends of a wide distribution, and more likely that they share a recent common ancestor. When combined with the observation that many other *Halichondria* species have been moved around by human activity, we feel this species should be treated as a possible introduction across some part of its range (though which part remains to be determined).

Halichondria bowerbanki Burton 1930

Figure 16

Material examined

190615, Praia to Lumiar, Viana do Castelo, Portugal, (41.73540, -8.87578), 7.1 m, 2019-06-15; 190616, Canto Marinho, Viana do Castelo, Portugal, (41.72910, -8.88071), 12.5 m, 2019-06-16; NIB0309, The Dorn, Northern Ireland, (54.43459, -5.54536), intertial (boulders), 2021-04-28; NIB0612, Carnalea, Northern Ireland, (54.66844, -5.70801), intertidal (rocky shore), 2021-10- 27; NIB0614, Mount Stewart, Northern Ireland, (54.54482, -5.61017), intertidal (low shore boulders), 2021-04-29; TLT1052, Santa Barbara Harbor, California, (34.40559, -119.68964), floating dock, 2021-06-01; TLT1072, Santa Barbara Harbor, California, (34.40559, -119.68964), floating dock, 2021-06-01; TLT1087b, Santa Barbara Harbor, California, (34.40559, -119.68964), floating dock, 2021-06-01; TLT1357, Santa Barbara Harbor, California, (34.40559, -119.68964), floating dock, 2023-09-01; TLT1358, Santa Barbara Harbor, California, (34.40559, -119.68964), floating dock, 2023-09-01; TLT1418, Santa Barbara Harbor, California, (34.40559, -119.68964), floating dock, 2023-09-01; TLT234, Arroyo Quemado Reef, California, (34.46775, -120.11905), 7-11 m, 2019-07-29; TLT337, Saddleback Ridge, Santa Cruz Island, California, (34.03817, - 119.52470), 5-12 m, 2019-10-18; TLT342, Coal Oil Point, California, (34.40450, -119.87890), 3-8 m, 2019-08-30; TLT415, Coal Oil Point, California, (34.40450, -119.87890), 3-8 m, 2019-10-25; TLT490, Coal Oil Point, California, (34.40450, -119.87890), 5 m, 2020-07-17; TLT738, Marina del Rey, California, (33.97228, -118.44653), floating dock, 2021-01-09; TLT980, Santa Barbara Harbor, California, (34.40559, -119.68964), floating dock, 2021-06-01; TLT981, Santa Barbara Harbor, California, (34.40559, -119.68964), floating dock, 2021-06-01.

## Diagnosis

Yellow, green, or blue-green Halichondria with a tangential ectosomal skeleton developed as a web of spicule tracts and individual spicules, with open spaces generally triangular or polygonal rather than oval. Choanosomal skeleton confused, with a slight tendency to form multispicular tracts. Spicules oxeas, 180–550 μm long, with averages per sample of 300–430 μm long and 7– 12 μm wide; usually the same length in the ectosome and the choanosome. Diagnosis relative to similar species requires DNA characters, as follows:

COX1. 301:C, 484:A . N=12.

28S. 8:G, 848:T. 853:G, 865:C, 1060:C, 1937:C, 2032:A, 2327:A. N=13–20 depending on position.

ND1. 264:G, 684:A, 691:C, 739:T. N=13–14 depending on position.

## Morphology

Previously described as variable in shape, from encrusting to cushion-shaped, with oscular chimneys and/or thread-like tendrils, and sometimes highly ramose with anastomosing branches (Ackers et al., 2007; Burton, 1930; Evcen et al., 2023). Samples examined here, with identities confirmed by DNA sequencing, do not include any forms with branches, threads, or oscular chimneys, though the previously sequenced sample from Ireland (BELUM:Mc4003) had these features.

Sponges from sheltered marinas were thinly encrusting, usually green or blue-green but one sample was yellow. Samples from subtidal rocky reefs were usually large cushions, pale yellow, but one sample was encrusting and green. Oscula are not always apparent, but when they are, they are nearly flush with the sponge surface or with raised rims, and often accompanied by a translucent channel running up one side (Figure 16B). White or beige in ethanol.

**Figure 16.**
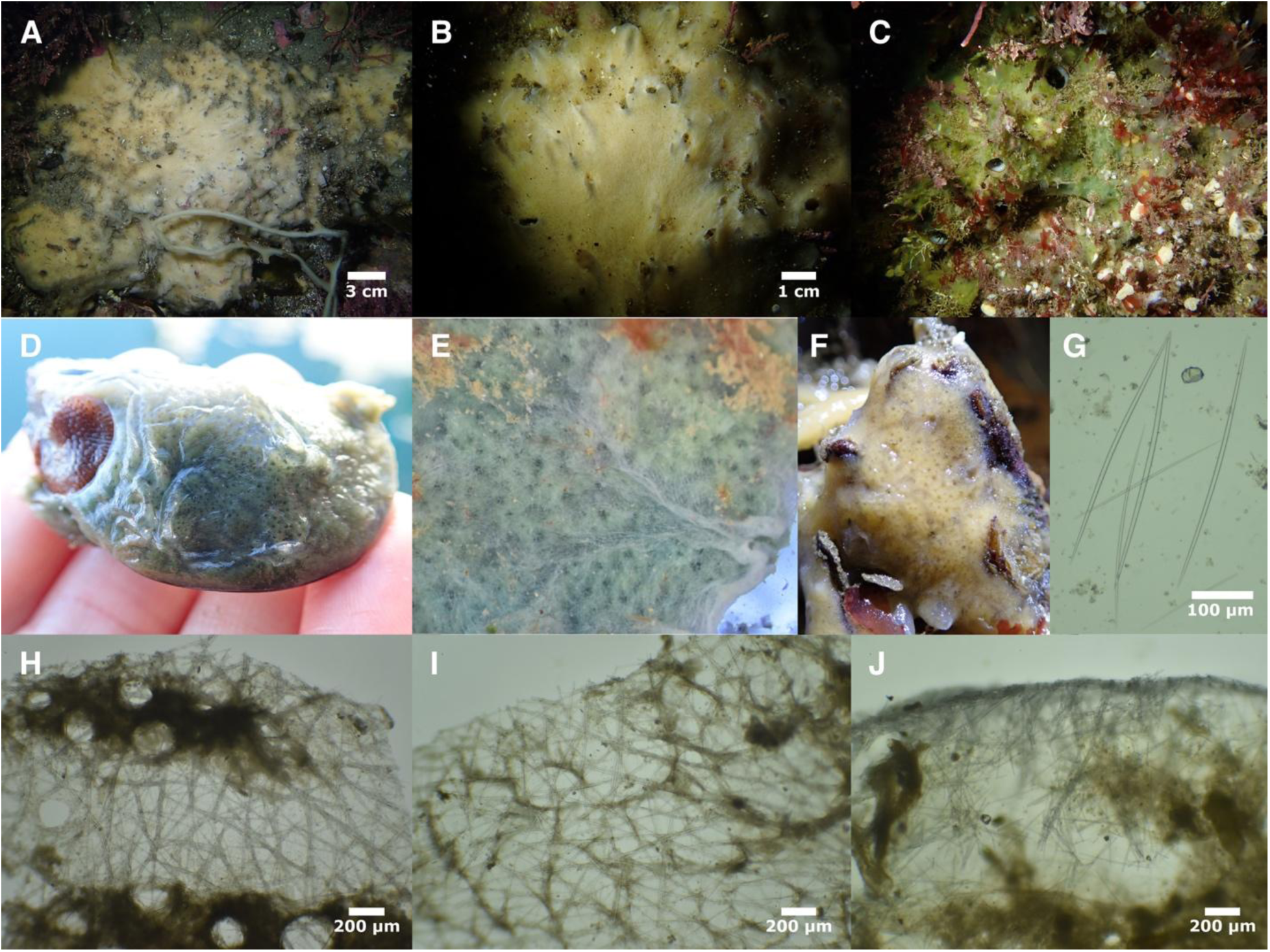
*Halichondria bowerbanki*. A & B (TLT490), C (TLT337), D (TLT738), E (TLT1052), F (TLT234). G: spicules (TLT234). Tangential ectosomal skeletons: H (TLT337), I (TLT234). J: cross-section showing choanosomal skeleton at sponge surface (TLT342).

## Skeleton

Tangential ectosomal skeleton is a web of spicule tracts and individual spicules, with open spaces generally triangular or polygonal rather than oval. The choanosomal skeleton is confused, with many free spicules, but meandering multispicular tracts are usually evident. No apparent spongin.

## Spicules

Nearly exclusively oxeas, though rarely a spicule can be found that is modified into a style. Spicules have the typical halichondroid shape: evenly curved or with a slight central bend, thickest in the center and tapering gently to sharp points at both ends. Spicules have a large range in lengths in both the ectosome and the choanosome, with no average difference between compartments (though individual samples varied from 5% shorter to 7% longer in the choanosome). Dimensions for 10 samples where ectosome and choanosome were measured separately are given in table 7. When spicules from all 16 samples with data are combined, spicules measure 185–379–547 x 2–8–17 μm, n=1312 for length, n=1092 for width. Average values varied considerably across samples, 307–423 μm long, and 7–12 μm wide.

**Table 7.** Spicule lengths of *Halichondria bowerbanki*. Values given as min–mean–max in μm.

| Specimen | Ectosome | Choanosome |
| --- | --- | --- |
| Pacific Ocean |  |  |
| TLT1052 | 273–356–454 n=45 | 203–359–410 n=40 |
| TLT342 | 350–433–490 n=40 | 291–411–528 n=41 |
| TLT490 | 340–412–477 n=40 | 265–430–506 n=40 |
| TLT980 | 231–333–384 n=40 | 249–323–371 n=46 |
| TLT981 | 251–322–389 n=40 | 213–323–379 n=40 |
| TLT234 | 228–424–539 n=97 | 196–422–547 n=96 |
| Atlantic Ocean |  |  |
| NIB0512 | 185–307–365 n=40 | 206–306–399 n=40 |
| NIB0614 | 292–389–467 n=45 | 320–416–491 n=45 |
| NIB0309 | 263–362–488 n=96 | 210–370–472 n=96 |
| 190615 | 334–408–469 n=40 | 249–391–507 n=40 |

## Distribution and habitat

This species was previously described as occurring throughout Europe, both coasts of North America, and the Korean Peninsula (de Voogd *et al*. 2024) However, many of these samples are likely to be members of the cryptic species newly described here, so the confirmed range of this species is limited to samples with genetic data. European samples with DNA confirmation are from Ireland, the Netherlands, Portugal, and the Black Sea. Confirmed Pacific samples were found in a small number of Southern California locations. Samples previously sequenced from Argentina expand the known range of this species into the Southern Hemisphere (Figure S1).

This distribution is likely a result of accidental human transport, and the species seems likely to be confirmed from additional locations in the future.

Previous publications have stated that this species is less tolerant of tidal exposure than *Ha. panicea*, but more tolerant of silted and brackish conditions (Ackers et al., 2007; Hartman, 1958; Vethaak et al., 1982). DNA-confirmed samples are consistent with this description, as the species was found in sheltered marinas and subtidal reefs, but intertidal samples were verified only in Ireland. Samples were collected on floating docks in several marinas, but *Ha. urca* sp. nov.*, Ha. zabra* sp. nov., and *Ha. hygeia* sp. nov. were much more common and widespread on floating docks than *Ha. bowerbanki.* Subtidal samples were collected in natural rocky reef habitat, up to 12 m deep, usually growing on holdfasts of *Macrocystis pyrifera*. Though not quantitative, our observations of this species in the California kelp forest demonstrate substantial differences in abundance through time. The species was among the most abundant sponges at two Southern California locations near Santa Barbara (Coal Oil Point and Arroyo Quemado Reef) in 2019 and 2020, but could not be relocated in those areas in 2023 or 2024.

## Remarks

Though *Ha. panicea* and *Ha. bowerbanki* cannot always be distinguished without genetic data, they do have average differences in morphological characters and habitat. However, *Ha. bowerbanki* is much harder to distinguish from other *Halichondria* such as *Ha. zabra* sp. nov.*, Ha. urca* sp. nov., and *Ha. pinaza* sp. nov., and many previous reports of *H. bowerbanki* are likely to be conflated with these species. In California, for example, ramose *Halichondria* from harbors and marinas have long been attributed to *Ha. bowerbanki*, but we found *Ha. bowerbanki* to be uncommon in California marinas, and found it to be thinly encrusting in those habitats when present.

It is likely that *Ha. bowerbanki* is introduced in some parts of its range. The presence of this species in the North Atlantic and North Pacific could potentially be explained by a natural range that includes the Arctic, though this seems less likely than for *Ha. panicea* because 1) genetic variation does not seem to be associated with geographic distance, 2) the species is found in human-modified habitats where introduced species are common, and 3) samples confirmed thus far suggest a less cold-tolerant range than for *Ha. panicea* (in the Pacific, where our sampling is most extensive, it has been found only in Southern California so far). Moreover, several *Ha. bowerbanki* were collected from Punta Verde, a beach in San Antonio Bay, Argentina, by Marianela Gastaldi (*pers. comm.*). Sequencing of the cox1 locus confirms these specimens are *Ha. bowerbanki* (MZ487239, MZ487236, MZ487221). It is highly unlikely that this population was founded by natural means.

Halichondria urca sp. nov.

Figures 17 & 18

## Material examined

Holotype: TLT1087 (CASIZ 245218), Santa Barbara Harbor, California, (34.40559, -119.68964), floating dock, 2021-06-01. Other samples: 15492 (BULA-0171), Cabrillo Beach, California, (33.70800, -118.28500), intertidal, 2019-08-20; 15925 (BULA-0236), Port of Long Beach, California, (33.72865, -118.23638), 10 m, 2019-08-20; MSH23x2 (YPM 111898), Mount Sinai Harbor, New York, (40.96345, -73.03646), 1 m, 2023-08-16; NIB0610 (BELUM.Mc2024.7), Carrickfergus Marina, Northern Ireland, (54.71028, -5.81237), floating dock, 2021-10-24; TLT1002 (SBMNH 718631), Farnsworth Bank, California, (33.34380, -118.51650), 21-27 m, 2021-07-10; TLT1050 (UCSB-IZC00069017), Santa Barbara Harbor, California, (34.40559, -119.68964), floating dock, 2021-06-01; TLT1051 (SBMNH 718632), Santa Barbara Harbor, California, (34.40559, -119.68964), floating dock, 2021-06-01; TLT1053 (UCSB-IZC00069018), Santa Barbara Harbor, California, (34.40559, -119.68964), floating dock, 2021-06-01; TLT1058 (UCSB- IZC00069019), Santa Barbara Harbor, California, (34.40559, -119.68964), floating dock, 2021- 06-01; TLT1060 (UCSB-IZC00069020), Santa Barbara Harbor, California, (34.40559, -119.68964), floating dock, 2021-06-01; TLT1062 (UCSB-IZC00069021), Santa Barbara Harbor, California, (34.40559, -119.68964), floating dock, 2021-06-01; TLT1063 (SBMNH 718633), Santa Barbara Harbor, California, (34.40559, -119.68964), floating dock, 2021-06-01; TLT1065 (UCSB- IZC00069022), Santa Barbara Harbor, California, (34.40559, -119.68964), floating dock, 2021- 06-01; TLT1066 (UCSB-IZC00069023), Santa Barbara Harbor, California, (34.40559, -119.68964), floating dock, 2021-06-01; TLT1068 (SBMNH 718634), Santa Barbara Harbor, California, (34.40559, -119.68964), floating dock, 2021-06-01; TLT1069 (UCSB-IZC00069024), Santa Barbara Harbor, California, (34.40559, -119.68964), floating dock, 2021-06-01; TLT1070 (UCSB- IZC00069025), Santa Barbara Harbor, California, (34.40559, -119.68964), floating dock, 2021- 06-01; TLT1071 (UCSB-IZC00069026), Santa Barbara Harbor, California, (34.40559, -119.68964), floating dock, 2021-06-01; TLT1073 (SBMNH 718635), Santa Barbara Harbor, California, (34.40559, -119.68964), floating dock, 2021-06-01; TLT1075 (UCSB-IZC00069027), Santa Barbara Harbor, California, (34.40559, -119.68964), floating dock, 2021-06-01; TLT1078 (UCSB- IZC00069028), Santa Barbara Harbor, California, (34.40559, -119.68964), floating dock, 2021- 06-01; TLT1081 (SBMNH 718636), Santa Barbara Harbor, California, (34.40559, -119.68964), floating dock, 2021-06-01; TLT1083 (SBMNH 718637), Santa Barbara Harbor, California, (34.40559, -119.68964), floating dock, 2021-06-01; TLT1085 (SBMNH 718638), Santa Barbara Harbor, California, (34.40559, -119.68964), floating dock, 2021-06-01; TLT534 (UCSB- IZC00069029), Point La Jolla, California, (32.85227, -117.27239), intertidal, 2020-02-08; TLT590 (UCSB-IZC00069030), Mission Bay: Quiviera Basin, California, (32.76421, -117.23829), floating dock, 2020-02-08; TLT591 (SBMNH 718639), Mission Bay: Quiviera Basin, California, (32.76421, -117.23829), floating dock, 2020-02-08; TLT593 (CASIZ 245219), Santa Barbara Harbor, California, (34.40559, -119.68964), floating dock, 2020-02-16; TLT594 (UCSB-IZC00069031), Santa Barbara Harbor, California, (34.40559, -119.68964), floating dock, 2020-02-16; TLT661 (UCSB-IZC00069032), Stearn’s, California, (34.41023, -119.68563), 3-5 m, 2020-08-25; TLT715 (UCSB-IZC00069033), Ventura Harbor, California, (34.24801, -119.26550), floating dock, 2020- 12-14; TLT720 (SBMNH 718640), Mission Bay: South Shores Boat Launch, California, (32.76406, -117.21750), floating dock, 2021-01-10; TLT733 (CASIZ 245220), Marina del Rey, California, (33.97228, -118.44653), floating dock, 2021-01-09; TLT740 (CASIZ 245221), Marina del Rey, California, (33.97228, -118.44653), floating dock, 2021-01-09; TLT747 (SBMNH 718641), San Diego Bay: Shelter Island Marinas, California, (32.71094, -117.23423), floating dock, 2021-01- 10; TLT764 (SBMNH 718642), Newport Bay: Balboa Peninsula, California, (33.60375, - 117.90052), intertidal (on piling), 2021-01-09; TLT766 (SBMNH 718643), Newport Bay: Balboa Peninsula, California, (33.60375, -117.90052), floating dock, 2021-01-09; TLT926 (UCSB- IZC00069034), Mission Bay: Quiviera Basin, California, (32.76421, -117.23829), floating dock, 2021-05-15; TLT951, Santa Barbara Harbor, California, (34.40559, -119.68964), floating dock, 2021-05-24; TLT951B (UCSB-IZC00069035), Nudi Wall, California, (32.69872, -117.27580), 18-28 m, 2021-05-16; TLT960 (UCSB-IZC00069036), Ventura Harbor, California, (34.24801, -119.26550), floating dock, 2021-05-24.

## Etymology

Named for the urca, one of the ships of the Spanish armada in the age of sail.

## Diagnosis

Yellow *Halichondria* with a tangential ectosomal skeleton developed as a web of spicule tracts and individual spicules, or as a sieve-like mat of spicules surrounding oval-shaped open spaces. Spicules oxeas, 100–440 μm long, with averages per sample of 230–360 μm long and 5–10 μm wide; usually very slightly longer in the choanosome than the ectosome. Diagnosis relative to similar species requires DNA characters, as follows:

COX1. 72:T, 93:A, 128:G, 139:C, 247:C, 310:C, 313:T, 364:T. 397:G, 412:T, 475:T, 538:G, 559:G, 634:T, 709:C, 823:G, 880:C, 979:C, 1040:T, 1041:G, 1102:T, 1120:T, 1150:A, 1321:T, 1432:G. N=9–21 depending on position. 28S. 693:A, 704:G, 829:C, 846:A, 884:T, 1359:G, 1373:C, 1656:G, 2331:G, 2102:C, 2328:T. N=9– 40 depending on position. ND1. 10:G, 121:C, 204:C, 228:C, 364:T, 448:G, 543:A, 658:T, 838:T. N=9–15 depending on position.

## Morphology

Samples discovered to date were bright yellow to yellowish brown, and did not contain any traces of green. Shape is variable from thin encrustations to semi-globular cushions. Oscula can be flush with surface, with raised rims, on volcano-shaped protrusions, or atop oscular chimneys (Figure 17). Sometimes has tendril-like projections. Translucent areas usually present, including channels leading up the sides of oscula, as often seen in *Ha. bowerbanki*. When removed from the substrate or torn, brooded bright yellow larvae may be present internally and where the sponge contacted the substrate (Figure 17F). In Santa Barbara Harbor, Southern California, larvae were seen in one of two sponges collected in February, a single sponge collected in May, and 5 of 20 sponges collected in June.

**Figure 17.**
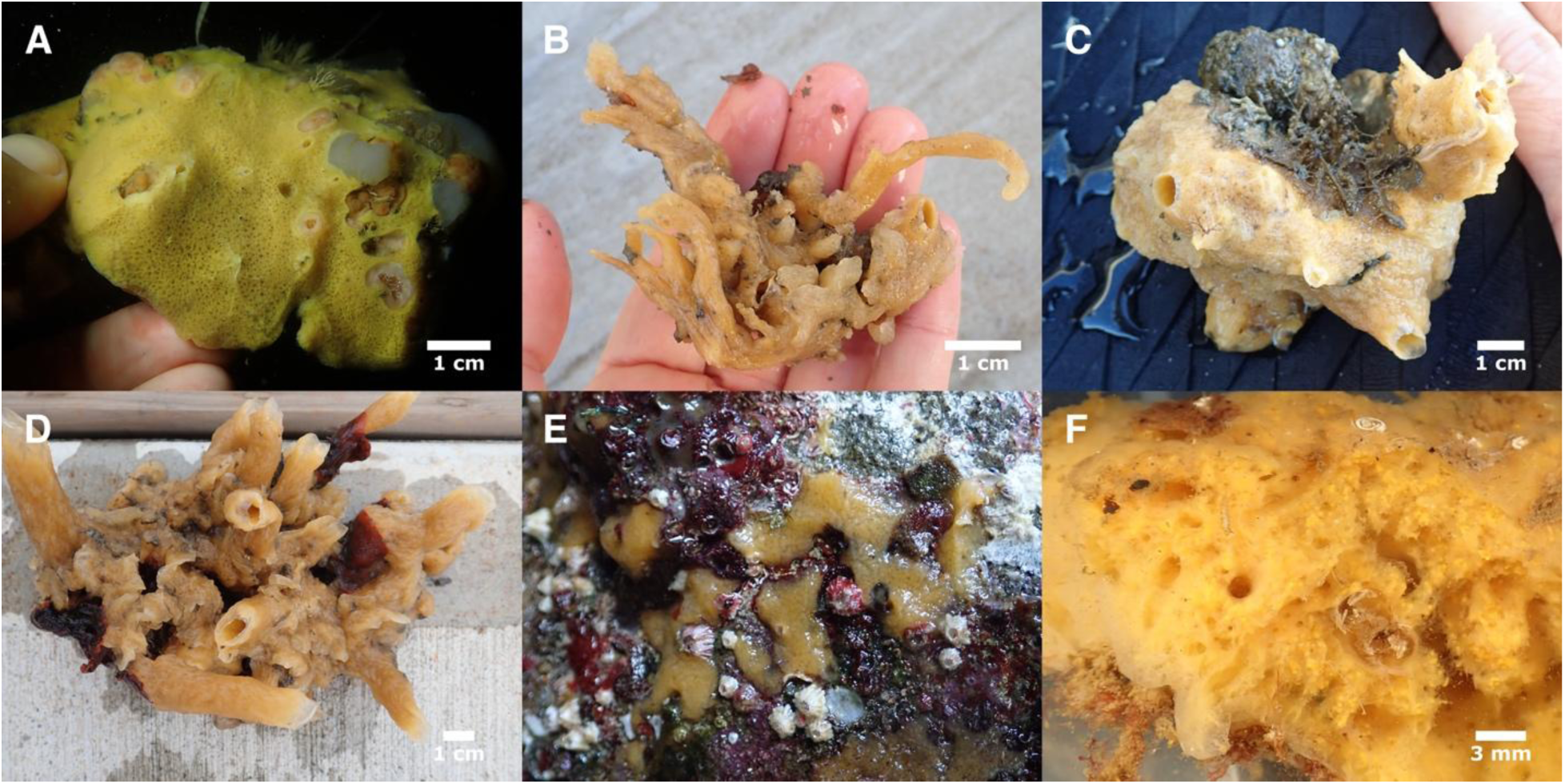
*Halichondria urca*. A (TLT 661), B (TLT740), C (TLT747), D (TLT1063), E (TLT534), F (TLT1069, sponge torn to show brooded larvae inside). A (subtidal) & E (intertidal) are in situ, others are pictured after removal from floating docks, before preservation.

## Skeleton

The tangential ectosomal skeleton is usually developed as a web of long meandering spicule tracts and single spicules, with open spaces between them irregular in size and shape.

However, some samples have more regular open spaces that are triangular and polygonal, and two samples had a mat of spicules that surrounded oval-shaped openings (Figure 18C).

**Figure 18.**
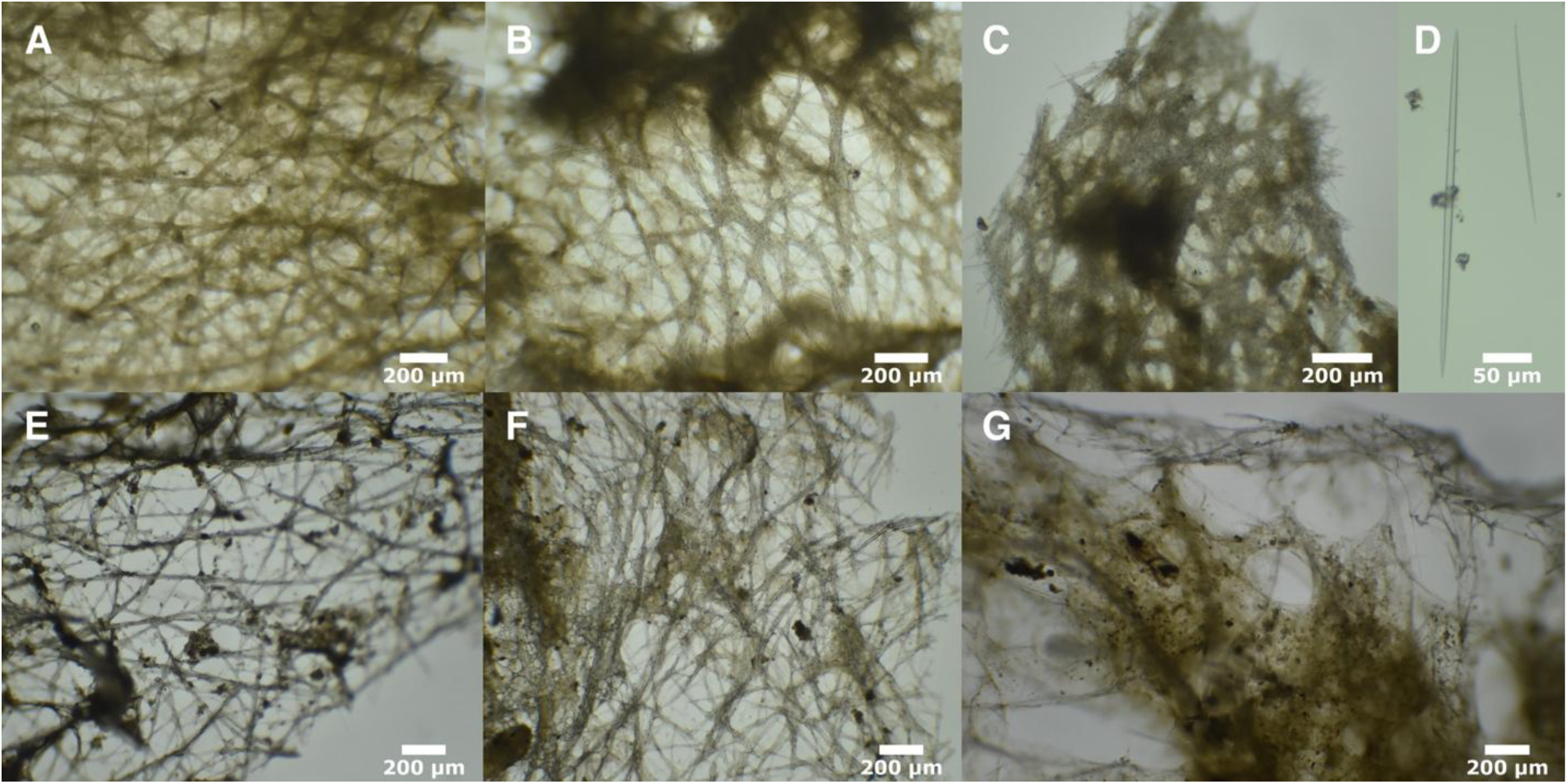
*Halichondria urca* skeleton and spicules. Tangential ectosomal skeletons: A (TLT926), B (TLT661), C (TLT534), E (TLT591), F (TLT594). D: spicules (TLT733). G: cross-section showing choanosomal skeleton at sponge surface (TLT594).

Choanosomal skeleton typically halichondroiid, with meandering spicules tracts and spicules in confusion, without apparent spongin.

## Spicules

Nearly exclusively oxeas, though rarely a spicule can be found that is modified into a style. Spicules have a typical halichondroid shape: evenly curved or with a slight central bend, thickest in the center and tapering to sharp points at both ends. Spicules have a large range in lengths in both the ectosome and the choanosome, with a very slight tendency to be longer in the choanosome (average of 3% longer across samples). Table 8 shows dimensions for 6 samples where ectosome and choanosome were measured separately. When all 19 samples with measurements are combined, spicules are 107–308–435 x 1–7–14 μm, n=1047 for length, n=863 for width. Average values varied across samples, with means of 238–359 μm long, 5–10 μm wide.

**Table 8.** Spicule lengths of *Halichondria urca*. Values given as min–mean–max in μm.

| Specimen | Ectosome | Choanosome |
| --- | --- | --- |
| Pacific Ocean |  |  |
| TLT926 | 264–315–368 n=50 | 273–327–362 n=50 |
| TLT951 | 241–296–335 n=40 | 242–320–364 n=40 |
| TLT1051 | 277–308–361 n=40 | 241–310–359 n=42 |
| TLT661 | 183–254–315 n=50 | 181–239–296 n=50 |
| TLT715 | 211–315–365 n=42 | 288–332–391 n=42 |
| Atlantic Ocean |  |  |
| NIB0610 | 259–346–416 n=40 | 316–372–425 n=40 |

## Distribution and habitat

Known from the intertidal to at least 28 m depth, but this species is much more common on floating docks in sheltered marinas than in other habitats investigated. Exceptions include encrusting samples found exposed at low tide at Point La Jolla and on a rocky reef in the Point Loma kelp forests, both near San Diego, California. Other samples found outside of marinas were on man-made debris: entangled trash 3 m below the Santa Barbara Wharf and an entangled rope at 21 m depth in the Farnsworth Offshore State Marine Conservation Area at Catalina Island, California.

Pacific samples are all from Southern California. Atlantic samples are from New York, Virginia, and Ireland. It was the most abundant species of *Halichondria* found in marinas in Southern California, where 48% of *Halichondria* (34 of 71 samples) were this species (sampling strategies were aimed at maximizing diversity rather than estimating relative abundance, so this number is likely an underestimate). *Halichondria urca* sp. nov. was not found at higher latitudes in the Northeast Pacific: only *Ha. zabra* sp. nov., *Ha. pinaza* sp. nov., and *Ha. hygeia* sp. nov. were collected in marinas from Central California to British Columbia, suggesting that *Ha. urca* sp. nov. may be less cold-tolerant than those species (though sampling in these areas was more limited than in Southern California).

## Remarks

This species is often morphologically indistinguishable from other members of the Bowerbanki group, but there were some features that made it more likely a sample would prove to be *Ha. urca* sp. nov. when genotyped. These include large size (semiglobular *Halichondria* ∼10 cm across, from floating docks, have consistently proved to be this species) and color (*Ha. urca* sp. nov. has not yet been found with any traces of green color).

It is possible that *Ha. urca* sp. nov. is introduced in some parts of its range. Its strong association with artificial substrate is consistent with an introduced species, as is finding samples in both the North Atlantic and North Pacific. The genetic differences seen between the Atlantic and Pacific are intriguing, however, and it is possible that additional sampling will reveal some of all Atlantic samples to be from an additional undescribed species.

Halichondria zabra sp. nov.

Figures 19 & 20

## Material examined

Holotype: TLT1064 (CASIZ 245222), Santa Barbara Harbor, California, (34.40559, -119.68964), floating dock, 2021-06-01. Other samples: TLT1077 (SBMNH 718644), Santa Barbara Harbor, California, (34.40559, -119.68964), floating dock, 2021-06-01; TLT1082 (UCSB-IZC00069037), Santa Barbara Harbor, California, (34.40559, -119.68964), floating dock, 2021-06-01; TLT1086 (UCSB-IZC00069038), Santa Barbara Harbor, California, (34.40559, -119.68964), floating dock, 2021-06-01; TLT1317, Santa Barbara Harbor, California, (34.40559, -119.68964), floating dock, 2023-09-01; TLT612 (CASIZ 245223), Jack London Square Marina, California, (37.79371, - 122.27757), floating dock, 2020-11-15; TLT623 (UCSB-IZC00069039), Jack London Square Marina, California, (37.79371, -122.27757), floating dock, 2020-11-15; TLT624 (SBMNH 718645), Jack London Square Marina, California, (37.79371, -122.27757), floating dock, 2020- 11-15; TLT712 (UCSB-IZC00069040), Ventura Harbor, California, (34.24801, -119.26550), floating dock, 2020-12-14; TLT722 (SBMNH 718646), Ventura Harbor, California, (34.24801, - 119.26550), floating dock, 2020-12-14; TLT737 (CASIZ 245224), Marina del Rey, California, (33.97228, -118.44653), floating dock, 2021-01-09; TLT751 (UCSB-IZC00069041), San Diego Bay: Shelter Island Marinas, California, (32.71094, -117.23423), floating dock, 2021-01-10; TLT756 (CASIZ 245225), Mission Bay: South Shores Boat Launch, California, (32.76406, -117.21750), floating dock, 2021-01-10; TLT757 (SBMNH 718647), Rainbow Harbor, California, (33.76184, - 118.19111), floating dock, 2021-01-09; TLT759 (UCSB-IZC00069042), Rainbow Harbor, California, (33.76184, -118.19111), floating dock, 2021-01-09; TLT760 (SBMNH 718648), Rainbow Harbor, California, (33.76184, -118.19111), floating dock, 2021-01-09; TLT761 (UCSB- IZC00069043), Rainbow Harbor, California, (33.76184, -118.19111), floating dock, 2021-01-09; TLT780 (SBMNH 718649), San Diego Bay: Shelter Island Marinas, California, (32.71094, - 117.23423), floating dock, 2021-01-10; TLT815 (SBMNH 718650), Ventura Harbor, California, (34.24801, -119.26550), floating dock, 2021-05-24; TLT874 (UCSB-IZC00069044), Tomales Bay Resort & Marina, California, (38.10792, -122.86237), floating dock, 2021-02-08; TLT967 (UCSB- IZC00069045), Ventura Harbor, California, (34.24801, -119.26550), floating dock, 2021-05-24; TLT978 (UCSB-IZC00069046), Ventura Harbor, California, (34.24801, -119.26550), floating dock, 2021-05-24; TLT979 (SBMNH 718651), Ventura Harbor, California, (34.24801, -119.26550), floating dock, 2021-05-24.

## Etymology

Named for the zabra, one of the ships of the Spanish armada in the age of sail.

## Diagnosis

Yellow or green *Halichondria* with a tangential ectosomal skeleton developed as a web of spicule tracts and individual spicules, with open spaces generally triangular or polygonal rather than oval. Choanosomal skeleton with spicule tracts and spicules in confusion. Spicules oxeas, 180–570 μm long, with averages per sample of 290–410 μm long and 6–12 μm wide; little-to-no difference in length in the ectosome and the choanosome. Diagnosis relative to similar species requires DNA characters, as follows:

COX1. No unique derived states.

28S. 4–5:GG. State 4:G is unique compared to all close relatives, but homoplastic with members of the Okadai group. The full sequence 4–5:GG is not shared with any other sequenced *Halichondria*. 831–832:GG. State 831:G is unique compared to all close relatives, but homoplastic with *H. dokdoensis.* The full sequence 831–832:GG is not shared with any other sequenced *Halichondria*. N=6.

ND1. 238:A. N=14.

## Morphology

Samples discovered to date were yellow, green, or a mix of the two; some yellow samples were very pale and nearly white (Figure 19F). Except in thinly encrusting samples, green pigment was more pronounced on the external surfaces of the sponge. Found as thin or thick encrustations, all with oscula elevated to various degrees: as volcano-shape protrusions, or atop oscular chimneys. Tendril-like projections may also be present, and these and oscular branches are sometimes anastomosing (Figure 19C). Translucent channels sometimes present, including those leading up one side of oscular protrusions, as often seen in other members of the Bowerbanki group. In Santa Barbara Harbor, Southern California, brooded bright yellow larvae were seen in one of three sponges collected in June (Figure 19G).

**Figure 19.**
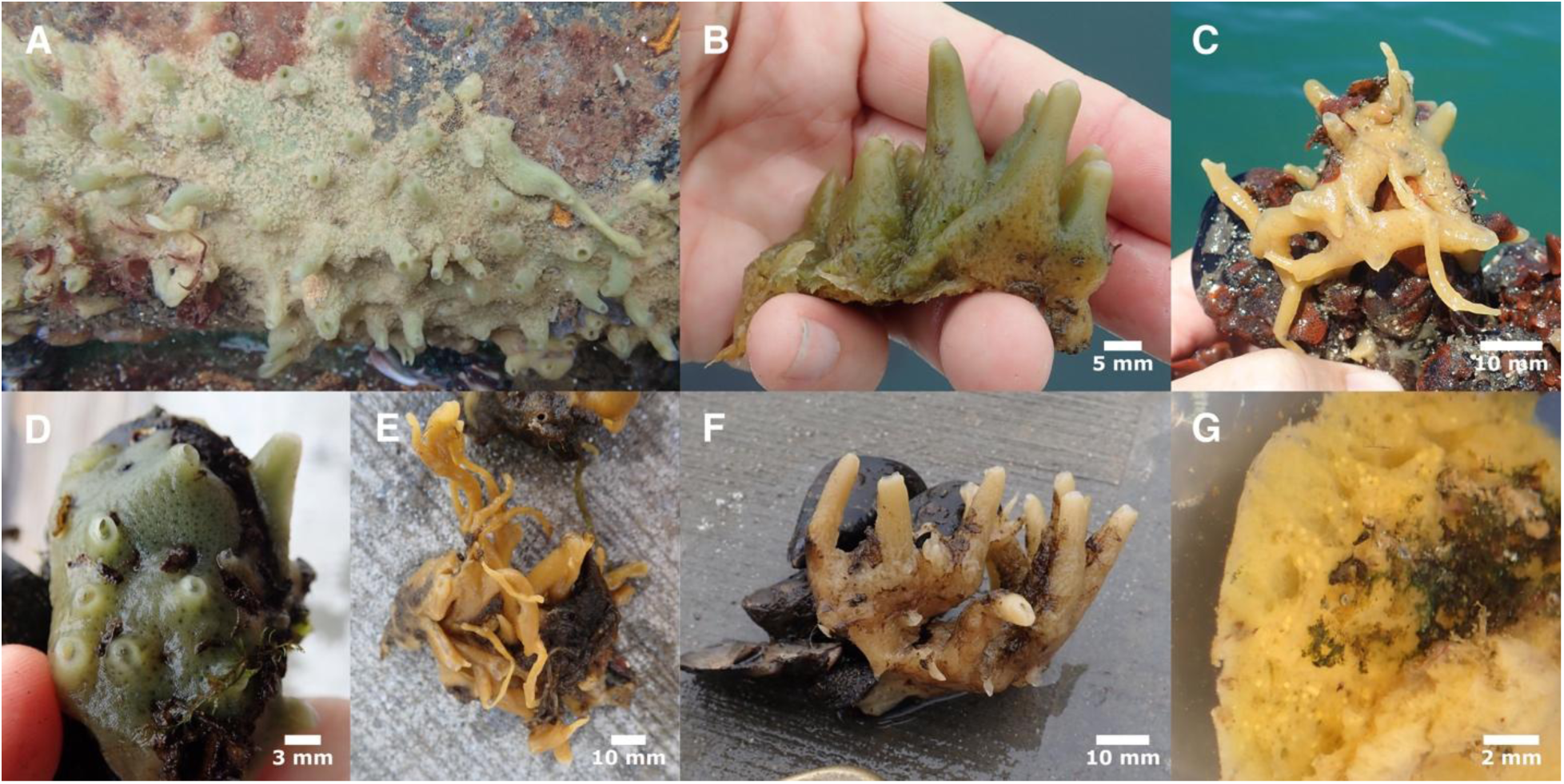
*Halichondria zabra*. A (TLT1317), B (TLT1077), C (TLT978), D (TLT759), E (TLT624), F (TLT761), G (TLT1086, sponge torn to show brooded larvae inside). A is in situ, others are pictured after removal from floating docks, before preservation.

**Figure 20.**
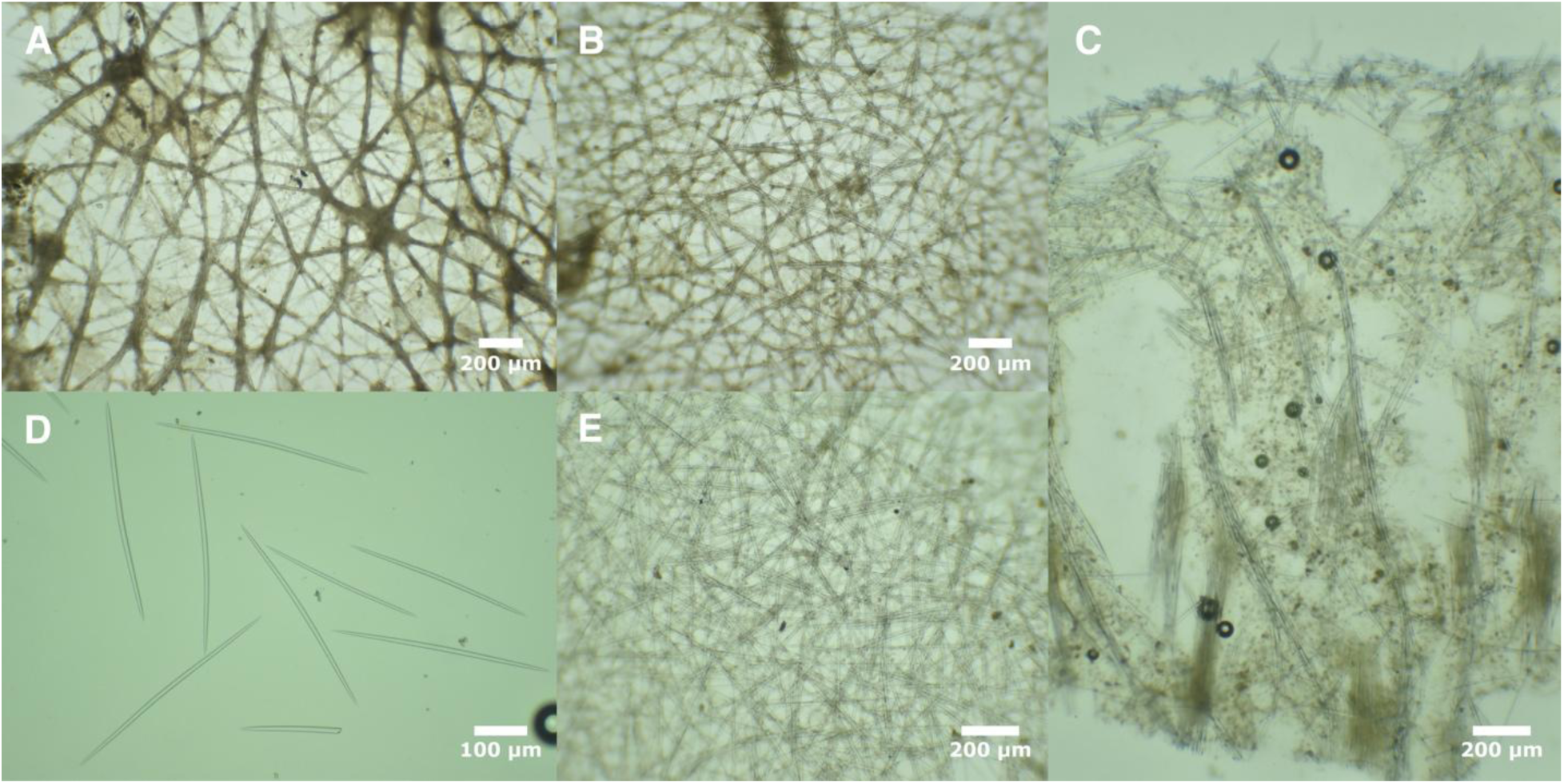
*Halichondria zabra* skeleton and spicules. Tangential ectosomal skeletons: A (TLT751), B (TLT624), E (TLT760). C: cross-section showing choanosomal skeleton at sponge surface (TLT623), D: spicules (TLT712).

## Skeleton

Tangential ectosomal skeleton with a web of spicule tracts and single spicules, with open spaces fairly regular in size and tending towards triangular or polygonal shapes. Choanosomal skeleton typically halichondroid, with vague spicules tracts and spicules in confusion, without apparent spongin.

## Spicules

Exclusively oxeas with a typical halichondroid shape: evenly curved or with a slight central bend, thickest in the center and tapering to sharp points at both ends. Spicules have a large range in lengths in both the ectosome and the choanosome, with little average difference in length between compartments. Table 9 shows dimensions for 5 samples where ectosome and choanosome were measured separately. When all 17 samples with measurements are combined, spicules measure 186–347–566 x 3–9–15 μm, n=1080 for length, n=912 for width. Average values varied across samples, with means of 294 to 407 μm long and 6 to 12 μm wide.

**Table 9.**
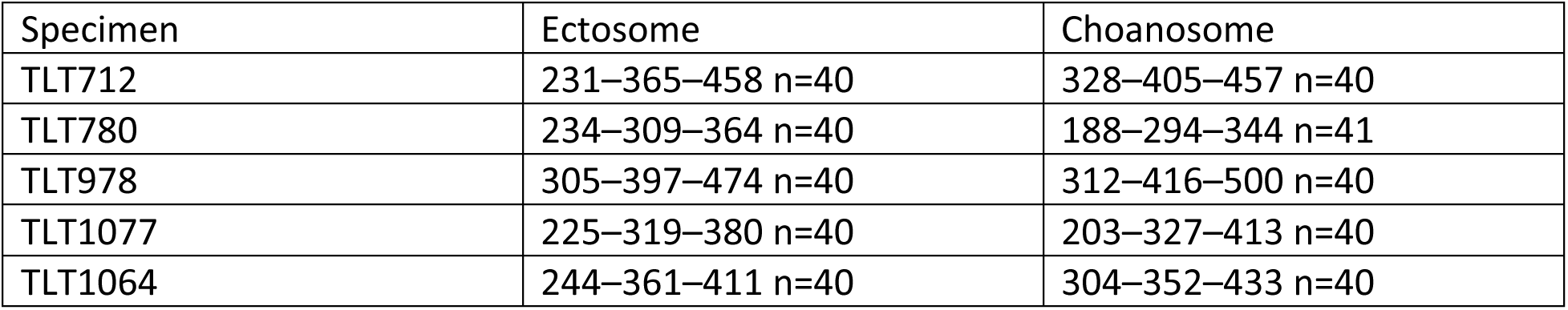
Spicule lengths of *Halichondria zabra*. Values given as min–mean–max in μm.

## Distribution and habitat

This species is known only from floating docks in California marinas, where it was found from Tomales Bay (the most northern site investigated) to San Diego Bay (the most Southern site investigated). It was the second most common *Halichondria* on docks in Southern California, where 19 of 71 *Halichondria* (27%) were this species (sampling strategies were aimed at maximizing diversity rather than estimating relative abundance, so this number is very approximate). Only *Ha. zabra* sp. nov. and *Ha. pinaza* sp. nov. have thus far been found on floating docks in Northern California, where they were at similar abundance (with much less sampling than Southern California).

## Remarks

Though this species has only been found in California thus far, we predict it will be found in additional regions as more samples are sequenced. Its strong association with floating dock habitat, where many introduced species are found, make it likely that it has been unintentionally introduced. Previously published cox1 sequences from New Hampshire, New York, Virginia, and Germany are consistent with a possible Atlantic distribution (see supplementary data), but could also be from *Ha. pinaza* (these two species cannot be differentiated at the most commonly sequenced region of this gene).

Halichondria pinaza sp. nov.

Figure 21

Material examined

Holotype: TLT869 (CASIZ 245217), Sausalito Yacht Harbor, California, (37.85930, -122.48044), floating dock, 2021-02-08. Other samples: NIB0603 (BELUM.Mc2024.5), Ballycastle Marina, Northern Ireland, (55.20687, -6.23962), floating dock, 2021-10-16; NIB0604 (BELUM.Mc2024.6), Ballycastle Marina, Northern Ireland, (55.20687, -6.23962), floating dock, 2021-10-16; TLT1056 (CASIZ 245216), Santa Barbara Harbor, California, (34.40559, -119.68964), floating dock, 2021-06-01; TLT1080 (UCSB-IZC00069016), Santa Barbara Harbor, California, (34.40559, -119.68964), floating dock, 2021-06-01; TLT873 (SBMNH 718629), Tomales Bay Resort & Marina, California, (38.10792, -122.86237), floating dock, 2021-02-08; TLT892 (SBMNH 718630), Home Bay, Drake’s Estero, California, (38.06934, -122.91711), intertidal, 2021-02-08; WMS21x22 (YPM 111896), West Meadow Beach, Stony Brook, New York, (40.92673, -73.14807), intertidal (rocks), 2021-05-26; WMS23x1 (YPM 111897), West Meadow Beach, Stony Brook, New York, (40.92673, -73.14807), intertidal (rocks), 2023-06-07.

## Etymology

Named for the pinaza, one of the ships of the Spanish armada in the age of sail.

## Diagnosis

Yellow or green *Halichondria* with a tangential ectosomal skeleton developed as a web of multispicular tracts and individual spicules, with open spaces tending towards triangular or polygonal shapes rather than ovals. Choanosomal skeleton with spicule tracts and spicules in confusion. Spicules oxeas, ranging from 90–370 μm in length, with averages per sample of 264– 308 μm in length and 6–10 μm in width; little-to-no difference in length in the ectosome and the choanosome. Diagnosis relative to similar species requires DNA characters, as follows:

COX1. No unique derived states.

28S. 4:T, 628:A, 669:T, 786:A, 803:C, 998:T. N=7–9 depending on position.

ND1. No unique derived states.

## Morphology

All samples found to date were encrustations with protruding oscular chimneys (Figure 21). Occurs in yellow, green, or both. Tendril-like projections not seen. Translucent channels sometimes present, including those leading up one side of oscular protrusions, as often seen in other members of the Bowerbanki group. In Santa Barbara Harbor, California, brooded bright yellow larvae were seen in two sponges collected in June (Figure 21E).

**Figure 21.**
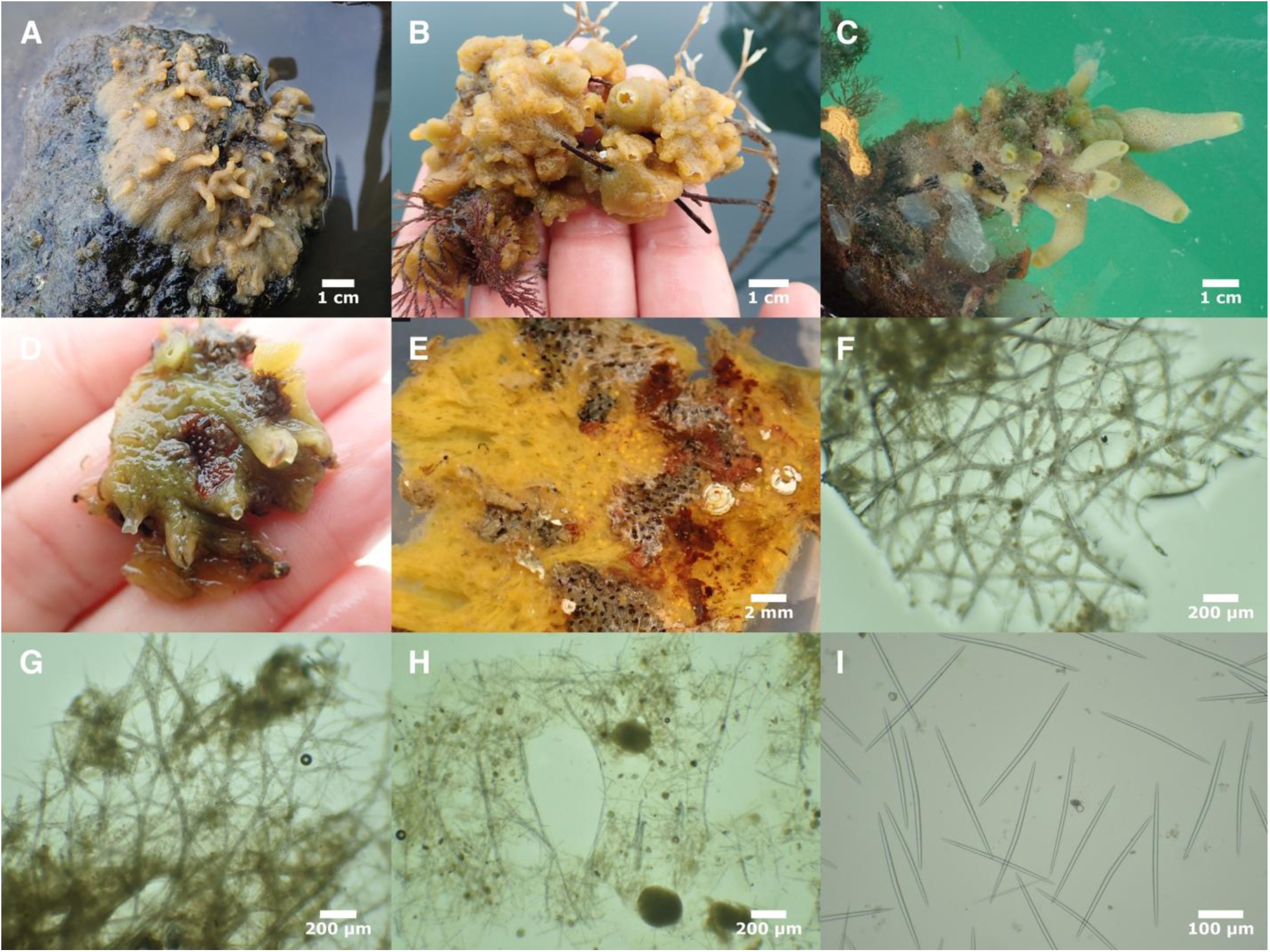
*Halichondria pinaza*. A (TLT892), B (TLT1080), C (TLT869), D & E (TLT1056, E shows underside where sponge attached to dock, with brooded larvae visible). Tangential ectosomal skeletons: F (NIB0603), G (NIB0604). H: cross-section showing choanosomal skeleton at sponge surface (TLT1080), I: spicules (TLT869).

## Skeleton

Tangential ectosomal skeleton with a web of spicule tracts and single spicules, with open spaces tending towards triangular or polygonal shapes (Figure 21F and 21G). Choanosomal skeleton typically halichondroid, with vague spicule tracts and spicules in confusion, without apparent spongin.

## Spicules

Nearly exclusively oxeas, though one sample (TLT892) had some aberrant spicules including styles, centrotylote oxeas, and oxeas with x-shaped cross pieces at one end. Oxeas have the typical halichondroid shape: evenly curved or with a slight central bend, thickest in the center and tapering to sharp points at both ends. Spicules have a large range in lengths in both the ectosome and the choanosome, with a very slight tendency to be longer in the choanosome (average of 3% longer across samples). Table 10 shows dimensions for 3 samples where ectosome and choanosome were measured separately. When all 6 samples with measurements are combined, spicules are 92–288–365 x 1–7–12 μm, n=432. Average values varied across samples, with means of 264 to 308 μm long, 6 to 10 μm wide.

**Table 10.** Spicule lengths of *Halichondria pinaza*. Values given as min–mean–max in μm.

| Specimen | Ectosome | Choanosome |
| --- | --- | --- |
| Pacific Ocean |  |  |
| TLT1056 | 158–272–311 n=55 | 236–281–330 n=57 |
| Atlantic Ocean |  |  |
| NIB0603 | 268–296–343 n=40 | 265–310–351 n=40 |
| NIB0604 | 261–305–357 n=41 | 179–302–346 n=40 |

## Distribution and habitat

This species is thus-far known from California, New York, and Ireland. All samples were found in sheltered bays and marinas. All were on floating docks except one sample on the underside of an intertidal boulder. It may be less tolerant of warm water than congeners such as *Ha. bowerbanki* and *Ha. zabra* sp. nov., as California samples were found from Drake’s Estero in Northern California to Santa Barbara in Southern California, but not in sites farther South. Of all the *Halichondria* collected on floating docks in Southern California, only 2 of 71 (3%) were this species; in contrast, *Ha. zabra* sp. nov. and *Ha. pinaza* sp. nov. had similar abundance in Northern California (though sampling effort was more limited there).

## Remarks

This species is likely introduced in some portion of its range. First, it was found primarily on floating docks, which are known to harbor many introduced species (Bulleri and Chapman, 2010; Glasby et al., 2007; Ruiz et al., 2000; Simkanin et al., 2012). Second, samples with very similar DNA sequences were found in California, New York, and Ireland, despite this species likely having limited larval dispersal (Maldonado, 2006; Xue et al., 2009). Sequences from the cox1 locus from samples collected by others from New Hampshire, New York, Virginia, and Germany may indicate this species is also present in those areas, but these could also be *Ha. zabra* sp. nov. (these two species cannot be differentiated at the most commonly sequenced region of this gene).

*Halichondria loma* Turner & Lonhart 2023 Material examined

Holotype: TLT1225 (CASIZ 236654 & UCSB-IZC00048445), North Monastery Beach, California, (36.52647, -121.92730), 12-29 m, 2021-09-22. Paratypes: TLT1012, Acropolis Street (Pt. Pinos), California, (36.64183, -121.93060), 9-18 m, 2021-08-09; TLT1024, Butterfly House, California, (36.53908, -121.93520), 9-20 m, 2021-08-10; TLT1130, Fire Rock Pescadero Point, California, (36.55898, -121.95110), 10-22 m, 2021-08-10; TLT514, Goalpost, San Diego, California, (32.69438, -117.26860), 12-15 m, 2020-02-08; TLT936, Six Fathoms, San Diego, California, (32.71000, -117.26860), 9-18 m, 2021-05-15; TLT976, Wreck of the Ruby E, California, (32.76680, -117.27620), 18-25 m, 2021-05-16; TLT989, Inner Pinnacle, California, (36.55910, - 121.96630), 10-18 m, 2021-08-10.

## Diagnosis

Yellow *Halichondria* with a tangential ectosomal skeleton developed as a web of mulitspicular tracts and individual spicules, with open spaces tending towards triangular or polygonal shapes, or as a lace-like mat of spicules surrounding oval-shaped open areas. Choanosomal skeleton with spicule tracts and spicules in confusion. Spicules oxeas, 290–650 μm long, with averages per sample of 460–550 μm long and 5–16 μm wide. This species is diagnosed relative to *H. sitiens* by a lack of papillae, and from all other *Halichondria* examined here by having longer spicules However, this difference is slight (the longest *Ha. bowerbanki* sample averaged only 39 μm shorter), so diagnosis is facilitated by DNA characters, as follows: COX1. No unique derived states. 28S. 5:A, 13:G, 46:G, 621:G, 674:T, 894:C, 2091:T, 2198:G. N=6–8 depending on position. ND1. 199:C, 205:G, 209:G, 220:A, 231:G, 248:T, 250:T, 700:G, 880:A. N=7.

## Remarks

This species was recently described by Turner and Lonhart (2023), and no additional samples have been discovered or examined since. Our contribution to the understanding of this species comes from DNA data: we used Illumina sequencing to increase the genetic data available for 7 of the 8 previously known samples, and used that data to locate DNA characters that aid in diagnosis.

This species is currently the closest known relative of the Bowerbanki group (Figure 1). In contrast to the species in that group, *Ha. loma* has not been found in sheltered bays or on human structures, and there is no indication that it may be introduced in California.

Genus *Hymeniacidon* Bowerbank, 1858

Encrusting or lobate Halichondriidae with small styles (<500 μm) for megascleres. Ectosomal tangential skeleton of intercrossing bundles end single megascleres. Choanosomal skeleton with ascending vague bundles and many loose, confusedly arranged megascleres (paraphrased from Erpenback and Van Soest, 2002). Approximately 50 species are currently considered valid (de Voogd et al., 2024).

Hymeniacidon perlevis (Montagu, 1814)

Figure 22

Synonyms

Spongia perlevis Montagu 1814

*Hymeniacidon sinapium* de Laubenfels 1930

The World Porifera Database lists 27 additional synonyms (de Voogd, et al., 2024)

## Material examined

CASIZ 180246, Newport Bay (*Hy. sinapium* type location), California, depth not known, 2006-09- 14; NIB0611, Carnalea, Bangor, Northern Ireland, (54.66844, -5.70801), intertidal (rocky shore crevices), 2021-10-27; NIB0615, Mount Stewart, Strangford Lough, Northern Ireland, (54.54482, -5.61017), intertidal (boulders), 2021-04-29; NIB0616, Portaferry, Strangford Lough, Northern Ireland, (54.38226, -5.55393), intertidal (boulders), 2021-10-10; TLT1054, Santa Barbara Harbor, California, (34.40559, -119.68964), floating dock, 2021-06-01; TLT1057, Santa Barbara Harbor, California, (34.40559, -119.68964), floating dock, 2021-06-01; TLT1059, Santa Barbara Harbor, California, (34.40559, -119.68964), floating dock, 2021-06-01; TLT1067, Santa Barbara Harbor, California, (34.40559, -119.68964), floating dock, 2021-06-01; TLT1074, Santa Barbara Harbor, California, (34.40559, -119.68964), floating dock, 2021-06-01; TLT1079, Santa Barbara Harbor, California, (34.40559, -119.68964), floating dock, 2021-06-01; TLT1265, Zebra Cove, California, (34.01000, -119.44000), 6-13 m, 2022-06-17; TLT129, Carpinteria Reef, California, (34.39163, - 119.54169), 3-6 m, 2019-07-31; TLT247, Carpinteria Reef, California, (34.39163, -119.54169), 3- 6 m, 2019-07-31; TLT595, Coal Oil Point, California, (34.40450, -119.87890), 3-8 m, 2020-07-17; TLT643, Santa Barbara Harbor, California, (34.40559, -119.68964), floating dock, 2020-02-16; TLT648, Santa Barbara Harbor, California, (34.40559, -119.68964), floating dock, 2020-02-16; TLT651, Coal Oil Point, California, (34.40450, -119.87890), 3-8 m, 2020-07-17; TLT658, Lechuza Point, California, (34.03437, -118.86146), intertidal, 2020-12-14; TLT730, Lechuza Point, California, (34.03437, -118.86146), intertidal, 2020-12-14; TLT736, Marina del Rey, California, (33.97228, -118.44653), floating dock, 2021-01-09; TLT762, Rainbow Harbor, California, (33.76184, -118.19111), floating dock, 2021-01-09; TLT79, Tajigus, California, (34.46279, - 120.10185), 3-8 m, 2019-07-01; TLT823, Elwood Beach, California, (34.42395, -119.90604), intertidal, 2020-12-29.

## Diagnosis

Yellow, green, orange, or red-orange *Hymeniacidon,* thinly or thickly encrusting, frequently with oscular chimneys, tendrils, or other fistulae. Ectosomal skeleton developed as a disorganized network of tangential spicule bundles, with some paratangential and protruding from the sponge surface. Choanosomal skeleton with multispicular tracts and spicules in confusion.

Spicules styles; in the Northeast Pacific, they range from 110–420 μm in length, with averages per sample of 170–290 μm in length and 4–6 μm in width. Diagnosed relative to all other *Hymeniacidon* in the Northeast Pacific, except *Hy. fusiformis*, by having a large range (>100 μm) in lengths within any one sample, with some spicules in excess of 300 μm. Diagnosis relative to *Hy. fusiformis* based on thinner, non-fusiform spicules. It is not well established if this species can be diagnosed morphologically in all other regions where it occurs, so DNA confirmation of clade membership is recommended.

## Morphology

Yellow, green, orange, or red-orange encrusting sponges, frequently with erect projections such as oscular chimneys and digitate tendrils. Dark red and pinkish forms reported from Europe and southern Africa (Ackers et al., 2007; Samaai et al., 2022). In silted habitats, it is often partially buried, with only vertical projections protruding from sediment. Translucent surface channels sometimes present, but it generally has a more opaque surface than *Halichondria* in the Panicea and Bowerbanki groups. Sometimes containing brooded yellow larvae.

## Skeleton

Ectosomal skeleton is more disorganized than *Ha. panicea*, but similar in that it is developed as a tangential mesh of spicule bundles (Figure 22E). Bundles of spicules also protrude from this mesh to create a microscopically hispid surface. Choanosomal skeleton with wispy spicule tracts and spicules in confusion (Figure 22F). No apparent spongin.

**Figure 22.**
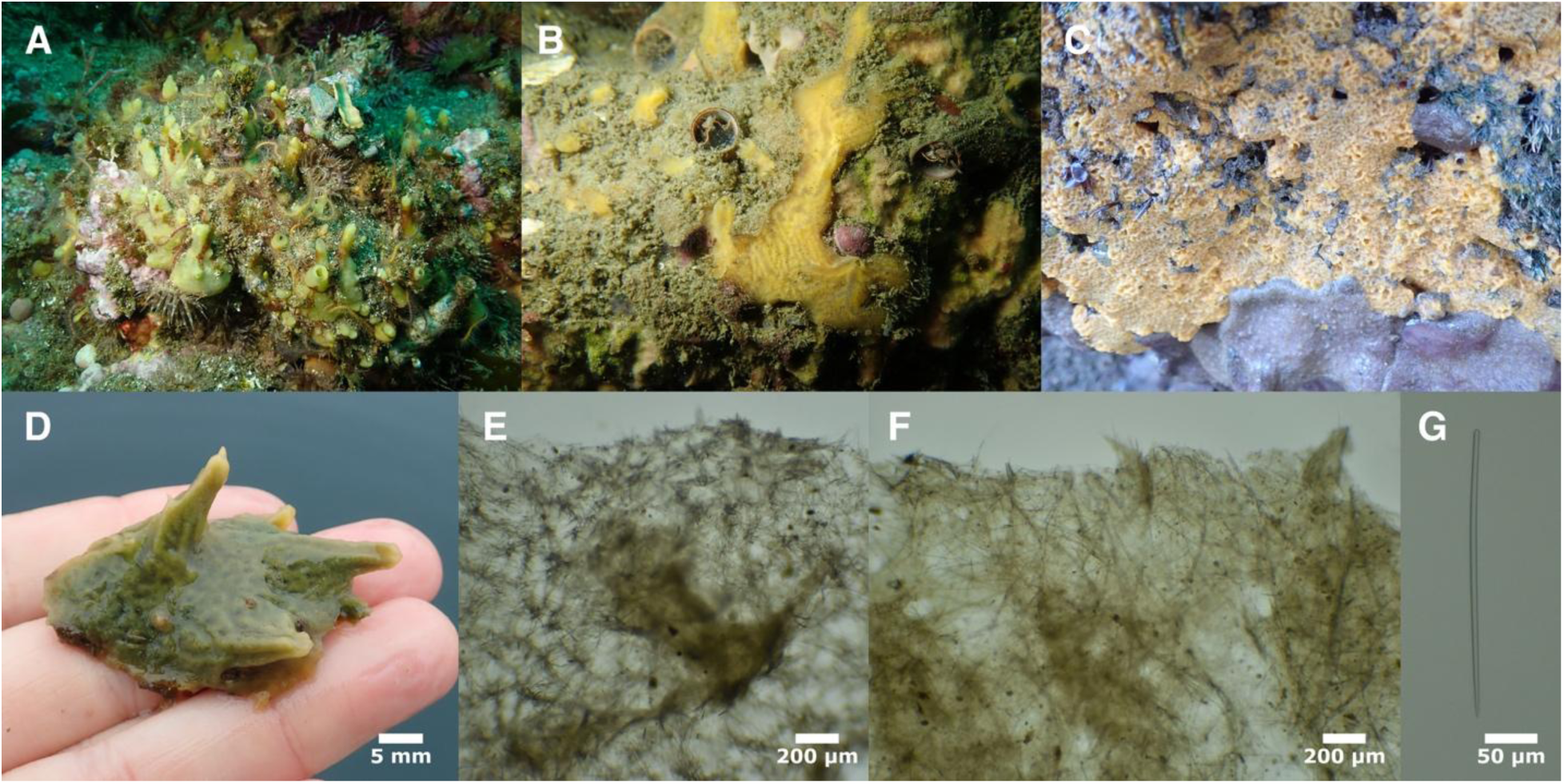
*Hymeniacidon perlevis*. A (TLT1265), B (TLT550), C (TLT730), D (TLT1067). A-C are in situ, with A and B subtidal and C exposed at low tide. D is removed from a floating dock. E: tangential ectosomal skeleton (TLT129). F: cross-section showing choanosomal skeleton at sponge surface (TLT129). G: typical style (TLT1057).

## Spicules

Exclusively styles, usually with uniform thickness but a significant minority have faint swelling at or near the head (Figure 22G). When data from 18 samples from California are combined, spicules measure 116–242–415 x 2–5–9 μm, n=660 for length, n=645 for width. Mean lengths per sponge varied from 177 to 289 μm, and mean widths from 4 to 6 μm. Sponges from other regions are reported to be similar in length but sometimes thicker (Harbo et al., 2021).

## Distribution and habitat

In the Northeastern Pacific this species is known from Ladysmith Harbor, British Columbia to Southern California. It is present in many other regions as well, including China, Korea, Europe, Brazil, Argentina, and South Africa (Samaai et al., 2022; Turner, 2020). Future DNA sequencing will likely confirm its presence in other regions such as Peru (Cóndor-Luján et al., 2023) and possibly New Zealand (Bergquist, 1970).

This species has a broad environmental niche, thriving in the intertidal zone, the shallow subtidal, areas with high levels of siltation, and brackish bays and estuaries. It is abundant on human structures in marinas but can also be found in relatively undisturbed natural habitats, such as those found at the Channel Islands in Southern California.

## Remarks

Turner (2020) compiled genetic and morphological data from many previous studies to argue that this species has a worldwide distribution. Here, we provide additional genetic data consistent with that conclusion. A complete mitochondrial sequence was previously published from Korea (Jun et al., 2015), and we find it to be nearly identical to mitochondrial genomes from California and Ireland.

Eleven samples were collected in Santa Barbara Harbor from April to July 2021, and checked for brooded larvae. None were seen in the 10 samples collected between April 21st and June 1st, but they were present in the final sample collected July 1st.

Hymeniacidon actites (Ristau, 1978)

Figure 23

## Synonyms

Leucophloeus actites Ristau 1978

## Material examined

Holotype: USNM 24526, Horeshoe Cove, Bodega Head, California, intertidal, 1977-01-06. Other samples: RBC1, Execution Rock Cave, Bamfield, British Columbia, (48.83170, -125.17830), not recorded, 2010-08-10; TLT1158, CRABS, California, (36.55377, -121.93840), 10-17 m, 2021-09-21; TLT227, Arroyo Quemado Reef, California, (34.46775, -120.11905), 7-11 m, 2019-07-29; TLT454, Middle Reef, Whaler’s Cove, Point Lobos, California, (36.52172, -121.93894), 6-15 m, 2019-11-23; TLT847, Cave Landings, California, (35.17535, -120.72240), intertidal, 2021-02-06; TLT856, Cave Landings, California, (35.17535, -120.72240), intertidal, 2021-02-06.

## Diagnosis

Encrusting yellow *Hymeniacidon* with partially translucent ectosome and no tassels or tendril-like projections. Spicules exclusively styles 150–200 μm long, with less than 75 μm difference between shortest and longest spicules in any one sample. Well-developed sieve-like tangential ectosomal skeleton reminiscent of *Ha. panicea*. Much more genetically similar to *Ha. panicea* than other species of *Hymeniacidon*.

## Morphology

Encrusting sponges, bright yellow to pale yellow-tan alive, with partially translucent ectosome. Beige or white preserved. Oscula usually apparent but small and flush with sponge surface.

General morphology indistinguishable from sympatric *Halichondria* such as *Ha. californiana* sp. nov. See Figure 23 for examples.

**Figure 23.**
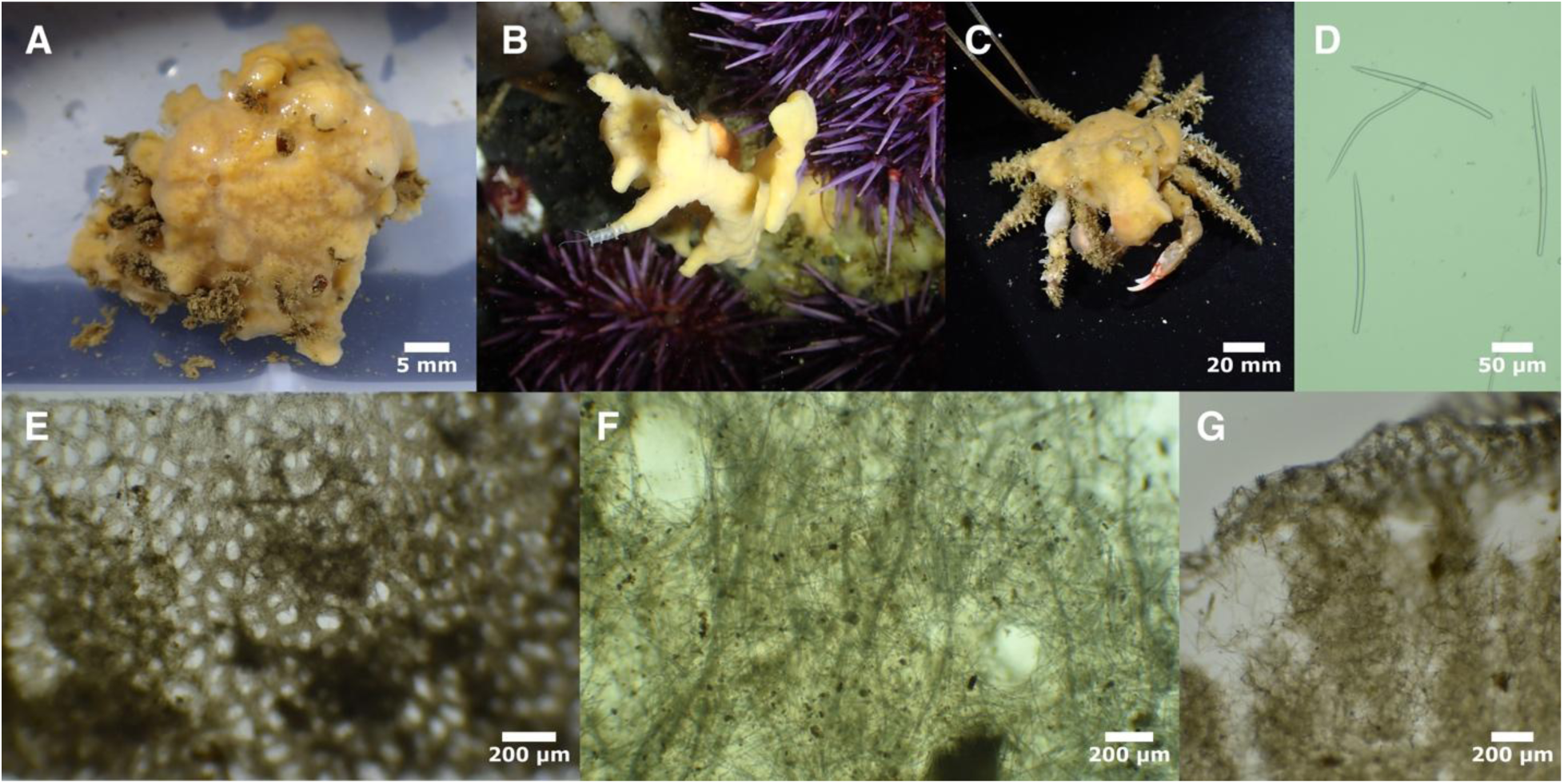
*Hymeniacidon actites*. A (TLT227), B (TLT1158), C (TLT454). Spicules: D (USNM 24526). E: tangential ectosomal skeleton (TLT227). F & G: cross-sections showing choanosomal skeleton (TLT227; ectosomal skeleton is visible at the top of G).

## Skeleton

Tangential ectosomal skeleton generally sieve-like, with a dense mat of spicules surrounding open areas that are mostly circular or oval shaped (Figure 23E). Choanosomal skeleton with many spicules in confusion, but wavy, meandering tracts can be found in some samples. No apparent spongin.

## Spicules

Exclusively styles, generally straight or evenly curved. Some samples contain spicules with very slight swelling at spicule heads or with multiple bends that make spicules somewhat sinuous. When all samples are combined, spicules measure 139–179–237 x 2–5–8 μm, n=250. Mean values per sample 156 to 199 μm long, 4 to 6 μm wide. Spicules from holotype: 168–196–237 x 4–6–8 μm, n=43.

## Distribution and habitat

Occurs in wave-exposed rocky intertidal habitats and shallow rocky reefs; not known from bays, harbors, or artificial structures. Known range previously limited to Northern and Central California (Lee et al., 2007), but is now extended North to British Columbia and South to Southern California. One sample was found on a *Loxorhynchus* decorator crab.

## Remarks

This species can be differentiated from *Hy. kitchingi* and *Hy. pierrei* sp. nov. due to the absence of tendrils and tufts, and from *Hy. perlevis* by color, surface patterning, smaller spicules, and a narrower range in spicule lengths. It is harder to distinguish from *Hy. ungodon* de Laubenfels 1932. As described below, the spicules and choanosomal skeletons in the two holotypes are likely indistinguishable. Indeed, the British Columbian sample examined here (RBC 018-00195- 002) was previously identified as *Hy. ungodon* by William Austin. We assign this sample, and the others we collected, to *Hy. actites* based on two apparent differences between the species: ectosomal skeleton and color (Ristau, 1978).

Regular sieve-like ectosomal patterning was reported in the original *Hy. actites* description and was seen in samples we examined (including the British Columbian sample). The *Hy. actites* holotype fragment examined by us was too small to investigate the skeleton, but the holotype was previously examined by Willard D. Hartman, and online images of his sections show this patterning as well (Yale Peabody Museum, sample IZ.074262). In contrast, the original description of *Hy. ungodon* states it has a rugose surface with a dense mat of ectosomal spicules in confusion (de Laubenfels, 1932). Absence of sieve-like ectosomal patterning in the *Hy. ungodon* holotype is not conclusive evidence of these species being distinct, as other samples we examined that show a sieve-like pattern in some portions of the ectosome will show a disordered mat in other places. *Hymeniacidon ungodon* is also described as having contrasting "mahogany-brown ectosome over yellowish-drab endosome", and this was called-out in the original description as being "characteristic and unique in the genus" (de Laubenfels, 1932). Color is known to be highly variable, so it is unclear if this trait is a legitimate reason to separate these species either. Genetic data from samples matching the original description of *Hy. ungodon* are therefore needed to resolve whether these two species are truly different, but for now, we assign these samples to *Hy. actites*.

Hymeniacidon ungodon de Laubenfels 1932

## Material examined

Holotype: USNM 22061, Point Lobos, Carmel, California, intertidal, 1930-07-12.

## Diagnosis

Encrusting sponges with mahogany-brown ectosome over yellowish-drab endosome and no tassels or tendril-like projections. Ectosome densely packed with spicules in confusion. Spicules exclusively styles 160–230 μm in length.

## Morphology

We were unable to locate any new samples that we could definitively assign to this species, and examined only a small fragment of the holotype. The original description states that they are encrusting sponges 1 cm thick, with a coarsely rugose surface; mahogany-brown ectosome over yellowish-drab endosome (de Laubenfels, 1932).

## Skeleton

The original description states that the ectosome is "densely packed with spicules in confusion". Only a small piece of ectosome could be located on the holotype fragment we examined, and it was a mat of tangential spicules in confusion, matching the previous description. The choanosomal skeleton was previously described as confused with vague ascending tracts, like other species of *Hymeniacidon*.

## Spicules

Exclusively styles, generally curved and of consistent thickness. 161–206–227 x 4–7–8 μm, n=50.

## Distribution and habitat

The holotype is from the intertidal zone in Central California, and was the only sample vouchered by de Laubenfels; the original description states that he has "seen this species several times in collections made by students" but the locations are not stated (de Laubenfels, 1932). Samples identified by others as this species have ranged from British Columbia to Southern California, from the intertidal to the subtidal (Lee et al., 2007). We examined many of these samples; the only one from Southern California is likely *Hy. perlevis*, and the others were either assigned to *Hy. actites* or were of uncertain affinity.

## Remarks

When Ristau (1978) described *Hy. actites,* he stated that it differed from *Hy. ungodon* in "coloration, growth form, and disposition of the ectosomal skeleton". We have been unable to locate any fresh samples matching the coloration described for *Hy. ungodon,* and all samples we examined had ectosomal skeletons matching *Hy. actites*. DNA was successfully amplified from only one sample previously identified as *Hy. ungodon*, from British Columbia, and it matched sequences from *Hy. actites*. Genetic data from samples matching the original description of *H. ungodon* are needed to resolve whether these two species are truly different.

Hymeniacidon kichingi (Burton, 1935)

Figure 24

TLT154, Elwood Reef, California, (34.41775, -119.90150), 9-15 m, 2019-05-15; TLT378, Coal Oil Point, California, (34.40450, -119.87890), 3-8 m, 2019-10-25; TLT412, Elwood Reef, California, (34.41775, -119.90150), 9-15 m, 2019-10-23; TLT424, Elwood Reef, California, (34.41775, - 119.90150), 9-15 m, 2019-10-23; TLT491, Coal Oil Point, California, (34.40450, -119.87890), 7 m, 2020-07-17; TLT597, Coal Oil Point, California, (34.40450, -119.87890), 3-8 m, 2020-07-17; TLT820, Platform Holly, California, (34.38993, -119.90656), 3-21 m, 2021-05-12; TLT941, Lazy Days, San Diego, California, (32.69415, -117.27110), 12-25 m, 2021-05-15; TLT977, Wreck of the Ruby E, California, (32.76680, -117.27620), 18-25 m, 2021-05-16.

## Diagnosis

Encrusting or cushion-shaped sponges, pale yellow, bright yellow, or beige shaded with pink or purple. Surface bearing many thin tassel-like extensions: in Californian samples these have a distinctive "frayed rope" appearance that differentiates this species from all others known from the region except *Hy. pierrei* sp. nov.. Styles are the only spicule, 110–260 μm long and 1–8 μm wide. Choanosomal skeleton is typically halichondroid, with meandering spicule tracts and spicules in confusion; differentiated ectosomal skeleton lacking. Diagnosis relative to *Hy. pierrei* sp. nov. requires DNA characters, as follows:

COX1. 718:T, 742:A, 769:G785:C, 786:T, 792:C, 796:C, 820:G, 937:C, 949:A, 1011:C, 1030:C, 1099:C, 1108:A, 1186:C, 1369:A, 1372:T, 1405:A, 1423:A, 1444:G, 1447:G, 1455:G, 1480:G, 1489:A, 1492:C. N=5–8 depending on position. 28S. 190:T, 206:G, 286:G, 471:G, 525:T, 529:T, 536:A, 540:C, 552:C, 558:A, 579:A, 603:T, 637:A, 641:T, 649:C, 660:G, 682:T, 697:T, 720:T, 789:C, 793:T, 797:C, 820:G, 826:A, 852:C, 859:T. N=5– 9.

ND1. Not assessed.

## Morphology

Form varies from thin encrustations to semiglobular cushions. California samples were replete with tassels: digitate projections that appear frayed due to profusely branching tips. All but the smallest samples in the British Isles are also covered in thin tassels, but available images do not clearly show the "frayed ends" appearance (Ackers et al., 2007). California samples pale to bright yellow; British Isles samples greyish-beige, with a tinge of pink or purple. Translucent surface channels, pores, and small scattered oscula visible on some living samples.

## Skeleton

Choanosomal skeleton typically halichondroid, with meandering multispicular tracts and spicules in confusion. No special ectosomal skeleton is apparent, though some spicules and spicule tracts are tangential in the surface of the sponge. Spicule tracts pierce the sponge surface at tassel tips. No apparent spongin.

## Spicules

Thin styles, straight or slightly bent. Most are of consistent thickness until tapering to a point, but a significant minority have very slight swelling at the head, with at least one sample (TLT491) containing some tylostyles. Some are slightly thinner at the head, and therefore weakly fusiform. The original description notes that tips are telescope-like, in that they taper in a stepwise fashion. This feature was subtly present in Californian samples, but only in some spicules; a more recent publication found it to be present on only about 10% of spicules in samples from the British Isles as well (Ackers et al., 2007). When data from 7 California samples is pooled, spicules measure 114–216–260 x 1–4–8 μm, n=192, very similar to reports on European samples (Ackers et al., 2007). Mean values per sponge were 206 to 223 μm long, 4 to 6 μm wide.

## Distribution and habitat

Previously known from just below the intertidal zone in the British Isles and Brittany. Samples examined here were found in 5 locations in Southern California, all subtidal, from 3 to 25 m depth. Two of these locations were artificial substrates (an oil platform and a scuttled ship), and three were natural reefs where the sponge was attached to rock.

## Remarks

This species was not previously known outside of the British Isles and Brittany, so it was surprising to find samples matching its description in Southern California. California samples are genetically identical to a 2085 bp sequence of the 28S ribosomal locus that was previously published from Ireland (Morrow et al., 2012; Thacker et al., 2013). This sequence includes the hypervariable "C2" barcoding region, making it highly likely that these samples are the same species. It also seems probable that the species has experienced a human-assisted range expansion, as a natural circumboreal distribution would likely result in some genetic divergence between these distant locations. This species can therefore be added to the growing list of Halichondriidae potentially introduced by human activity.

This species is not closely related to the type species *Hy. perlevis*, and instead forms a distinct clade with *Hy. pierrei* sp. nov., *Hy. fusiformis*, an unidentified *Hymeniacidon* from the North Atlantic, and *Semisuberites cribrosa* (Figure 1). Published sequences of cox1 indicate additional samples in this clade, identified as belonging to a variety of genera, with a Hawaiian sample identified as *Hy. gracilis* being the only additional sample identified as a *Hymeniacidon* (Figure S1).

Hymeniacidon pierrei sp. nov.

Figure 25

Material examined

Holotype: TLT599 (CASIZ 245233), Coal Oil Point, California, (34.40450, -119.87890), 6.7 m, 2020-07-17. Other sample: TLT531 (SBMNH 718663), Coal Oil Point, California, (34.40450, - 119.87890), 3-8 m, 2020-06-30.

## Etymology

Named for Christoph Pierre, marine naturalist and Director of Marine Operations at the University of California, Santa Barbara.

## Diagnosis

Encrusting *Hymeniacidon* bearing many thin tassel-like extensions. Styles are the only spicule, 100–230 μm long and 1–7 μm wide. Choanosomal skeleton is typically halichondroid, with meandering spicule tracts and spicules in confusion; differentiated ectosomal skeleton lacking. The combination of tassel-like extensions, no differentiated ectosomal skeleton, and very small, thin styles in a small size range differentiate this species for all others save *Hy. kitchingi*.

Diagnosis relative to *Hy. kitchingi* requires DNA characters, as follows: COX1. 486:C, 502:C, 598:C, 727:G, 792:A, 886:C, 1034:C. N=1.

28S. 170:T, 181:T, 182:T, 203:A, 262:T, 277:C, 278:A, 377:A, 463:G, 500:A, 512:C, 523:T, 530:T, 535:A, 551:T, 552:A, 559:T, 562:T, 603:G, 606:T, 608:C, 609:A, 611:G, 615:A, 623:A, 625:A, 642:C, 654:G, 672:T, 682:G, 704:C, 720:C, 769:A, 771:A, 778:A, 789:T, 792:A, 795:G, 796:A, 818:A, 819:C, 827:T, 841:T, 850:T, 884:G, 894:C. N=2.

ND1. Not assessed.

## Morphology

Thinly encrusting sponges replete with tassels: digitate projections that appear frayed due to profusely branching tips (Figure 25A & 25B). Translucent surface channels, pores, and small scattered oscula visible on living samples. Pale yellow alive, white when preserved.

**Figure 24.**
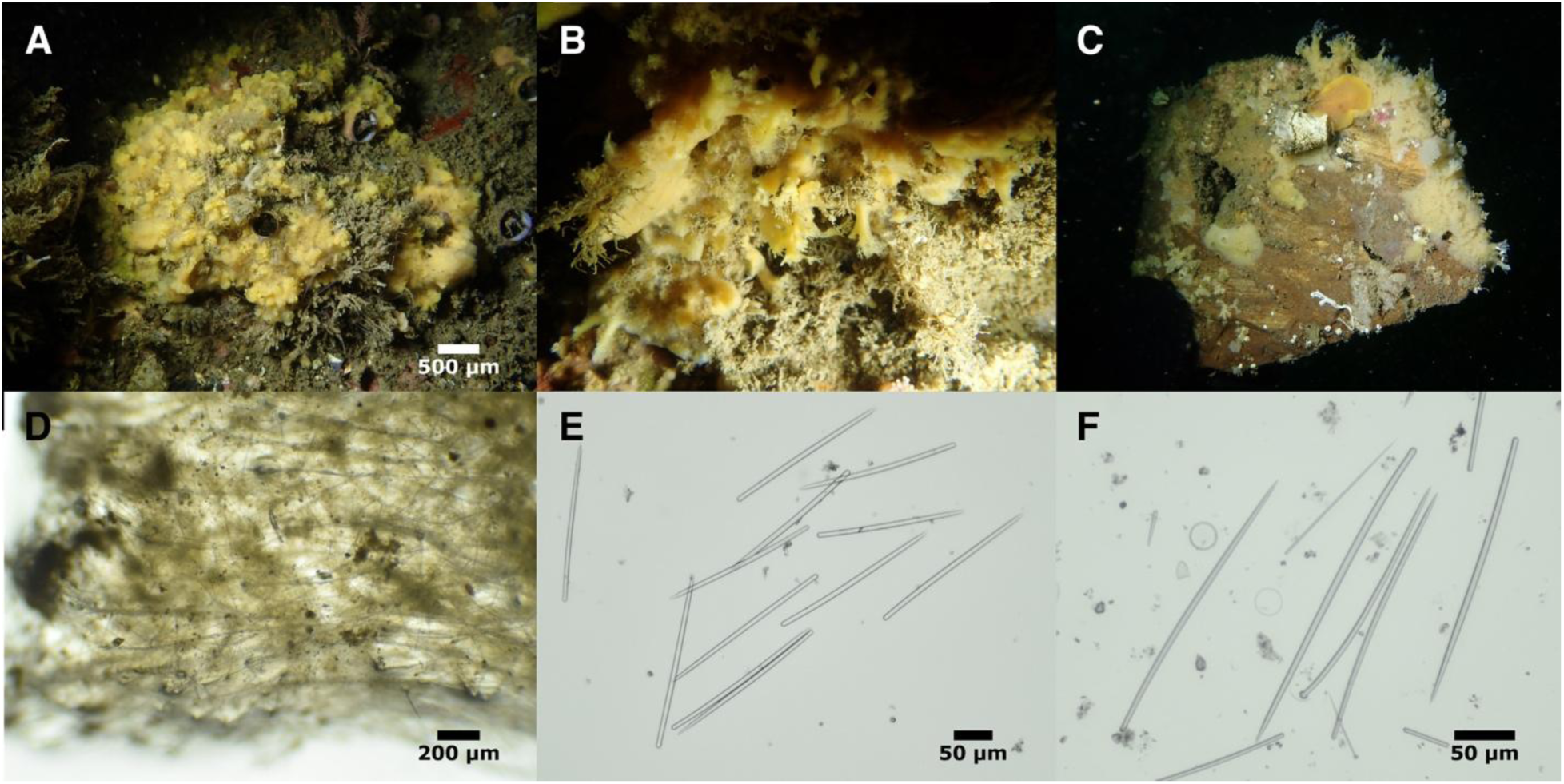
*Hymeniacidon kitchingi*. A (TLT597), B (TLT154), C (TLT412). D: cross-section showing choanosomal skeleton at sponge surface (TLT154). Spicules: E (TLT154), F (TLT491).

**Figure 25.**
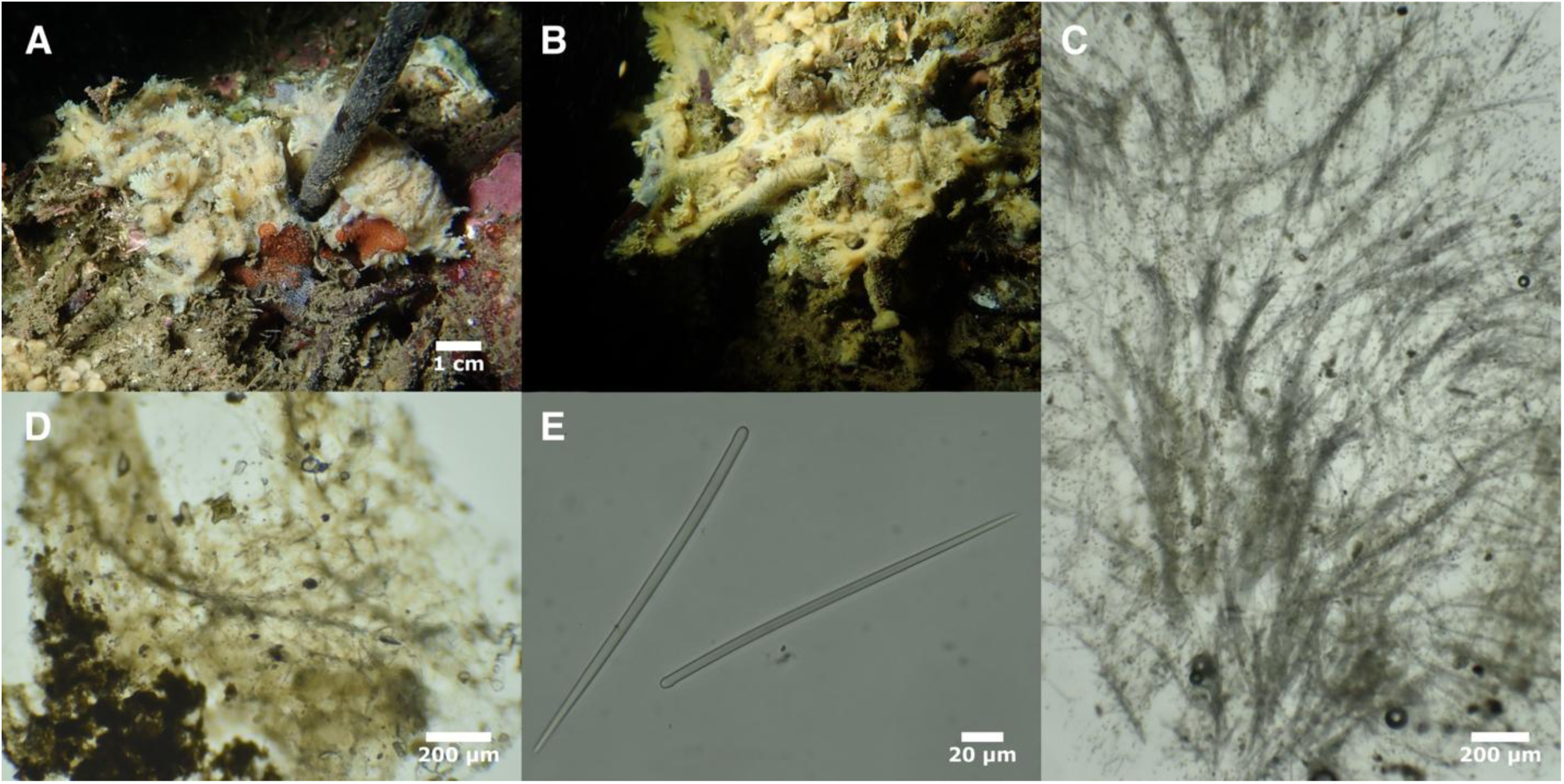
*Hymeniacidon pierrei*. A (TLT599), B (TLT531). C: cross-section showing choanosomal skeleton in tuft (TLT531). D: tangential spicules and tracts in ectosome (TLT599). Spicules: E (TLT531).

## Skeleton

Choanosomal skeleton typically halichondroid, with meandering multispicular tracts and spicules in confusion. Branching and somewhat plumose spicule tracts are well developed in tassels, where they pierce the sponge surface at tassel tips. No special ectosomal skeleton is apparent, though some spicules and spicule tracts are tangential in the ectosome. No apparent spongin.

## Spicules

Thin styles, straight or slightly bent. Most are of consistent thickness until tapering to a point, but a minority have very slight swelling at the head. Like *H. kitchingi*, the tips of some spicules taper in a subtly stepwise fashion. When data from both samples is pooled, spicules measure 109–187–229 x 1–4–7 μm, n=121. Means per sponge were 183 and 199 μm in length and 4 and 5 μm in width.

## Distribution and habitat

This species is only known from the shallow (3–8 m) subtidal at Coal Oil Point, Santa Barbara, California. It was found on the decaying holdfasts of dead *Macrocyctis pyrifera*.

## Remarks

*Hymeniacidon kitchingi* was discovered in California in 2019 at Coal Oil Point and nearby Elwood Reef. An additional tasseled *Hymeniacidon* was collected on a return trip to Coal Oil Point in 2020, but when it was sequenced, it was very genetically distinct from *Hy. kitchingi*. An additional trip later in 2020 yielded both genotypes, collected side-by-side. With 13% sequence divergence (Fst = 0.97) at the 28S locus and 10% sequence divergence (Fst = 0.32) at the cox1 locus, these sequences are clearly from a different species. Indeed, at cox1, *Hy. pierrei* sp. nov. are more closely related to samples collected in Hawaii (Genbank accession KY565326, identified as *Hy. gracilis*) and Australia (Genbank accession KJ620398, unidentified) than to *Hy. kitchingi*.

It is unclear whether *H. pierrei* sp. nov. is native to California or introduced. It was found on a natural reef that is now part of a marine protected area, but this site was previously highly disturbed, as it housed a pier and oil derricks as part of the Elwood Oil Field. This area is one of two places that the (possibly introduced) *Hy. kitchingi* has been found, so it seems possible that *Hy. pierrei* sp. nov. is introduced as well.

Hymeniacidon fusiformis Turner & Lonhart 2023

## Material examined

Holotype: TLT1194 (CASIZ236657 & UCSB-IZC00048452), Inner Carmel Pinnacle, Carmel, California, (36.55852, -121.96820), 10–24 m, 2021-09-22. Additional sample: FH12x28 (YPM 111899), Bell Island (West Face), Washington, (48.59667, -122.98150), 15 m, 2012-05-18.

## Diagnosis

Thickly encrusting white or yellow sponges, sometimes with irregularly projecting fingers and lobes, but not thin tentacle-like extensions or tassels. Styles are the only spicule, slightly fusiform, 150–410 μm in length and 4–19 μm in width. Maximum style length and large range in spicule lengths differentiate this species from sympatric *Hymeniacidon* except *Hy. perlevis*, while the wide range in spicule widths is unique in this geographic region. Choanosomal skeleton is typically halichondroid, with spicule tracts and spicules in confusion. The ectosomal skeleton is developed as a sieve-like mesh surrounding triangular open spaces or as a dense unstructured mat of tangential spicules.

## Spicules

Exclusively styles; most are very slightly fusiform, so that they are thicker in the center than near the head of style. Length and width are both highly variable; distributions are continuous but strongly bimodal, despite not showing any differences in size in the ecotosome vs. the choanosome. Spicules in the holotype measure 186–281–409 x 4–10–18 μm (n=345), with length modes 210 and 350 μm and width modes 9 and 14 μm. The new sample from Washington is similar, 153–270–396 x 6–11–19 μm (n=80), with length modes 260 and 330 μm and width modes 8 and 13 μm.

## Distribution and habitat

Known from two samples, the first from a natural subtidal pinnacle in Carmel Bay, Central California, 10–24 m depth, the second from a submerged rock wall, Bell Island, Washington, 15 m depth.

## Remarks

This species was recently described by Turner & Lonhart (2023) from a single sample. Here we report additional genetic data from this sample. We used Illumina sequencing to completely sequence the nuclear ribosomal locus; coverage was insufficient to assemble the mitochondrial genome, but we were able to extract 777 bp of the cox1 locus. We also report the discovery of one additional sample. This new sample conforms well to the morphological description of the holotype, and is only 0.4% divergent at the 28S locus.

The closest relative to this species in our molecular phylogenies is a *Hymeniacidon* sp. we collected and sequenced from the Bay of Fundy, in the Canadian North Atlantic (sample Quoddy115 in Figures 1 & 3). This sample is 1.2% divergent at the cox1 locus and 0.4% divergent at the 28S locus when compared to Pacific samples, which could indicate a species-level difference or not. However, it is also has much shorter and thinner spicules that are not fusiform, and these combined morphological and genetic differences indicate it is likely a different, possibly undescribed, species. These species fall in a larger clade including *Hy. kitchingi*, *Hy. pierrei* sp. nov., *S. cribrosa* and samples from Genbank identified as belonging to a variety of other genera (Figure S1).

*Hymeniacidon globularis* Ott, McDaniel & Humphrey, 2024

## Material examined

None

## Diagnosis

Light yellow globular sponges with a prominent central oscule; subtylostyles with elliptical heads are the only spicule. Distinguished from all other *Hymeniacidon* in the region by "puffball" shape, large styles (300–450 μm long and 5–13 μm wide), and having elliptical subtylostyles as the primary spicule shape.

## Distribution and habitat

Known only from the subtidal, 17-24 m, in British Columbia.

## Remarks

This species was recently described by Ott, McDaniel & Humphrey (Montagu 1814), and is included here for completeness. See their description for additional details.

Genus *Semisuberites* Carter, 1877

Funnel-shaped sponges arising from a stalk. Choanosomal skeleton of branching spicule tracts run mostly parallel to the sponge surface, but give rise to tracts that branch off to intersect the sponge surface. These flare to form bouquets that pierce the surface of the sponge. Some spicules are also confused throughout the choanosome and tangentially arranged within the ectosome. Spicules styles and subtylostyles. Genus is monotypic.

*Semisuberites cribrosa* (Miklucho-Maclay, 1870)

Figure 26

Synonyms

Veluspa polymorph*a* var. *cribrosa* Miklucho-Maclay, 1870

*Semisuberites arctica* Carter, 1877

*Stylissa stipitata* de Laubenfels, 1961

The World Porifera Database lists 22 additional synonyms (de Voogd, et al., 2024)

## Material examined

BHAK-08809, Nalau Pass, British Columbia, (51.784, -128.101), 30 m, 2019-05-25; BHAK-09371, Wedgeborough Point, British Columbia, (51.647, -127.956), 20 m, 2019-05-27; BHAK-10379, Wedgeborough Point, British Columbia, (51.647, -127.956), 25 m, 2019-05-27.

## Diagnosis

Soft, fragile, funnel-shaped sponges arising from a stalk. Styles are the only spicule, slightly fusiform, 110–450 μm long 1–12 μm wide (styles reportedly up to 600 μm long and 16 μm wide in other regions). Prominent spicule tracts run parallel to the surface of the sponge, branching off to intersect the sponge surface, where they form spicule brushes in the ectosome.

## Morphology

Funnel or trumpet-shaped sponges arising from stalks that attach to the substrate with root-like structures (Figure 26B). Stalk may branch and give rise to multiple funnels (Figure 26A). Sponges soft and fragile, with distal lip often frayed (Figure 26B). Velvety surface. Tan alive, white in ethanol.

**Figure 26.**
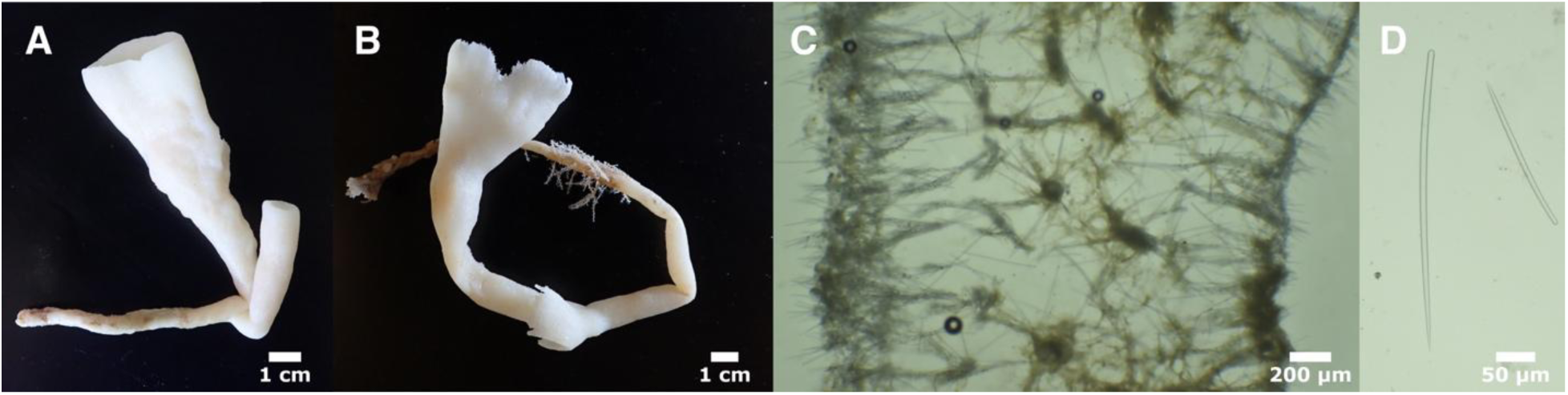
*Semisuberites cribrosa*. A (BHAK-08809), B (BHAK-09371). C: cross-section showing choanosomal skeleton near sponge rim, with external surface on the left and internal surface on the right; round structures in center are cross-sections of spicule tracts (BHAK-08809). Spicules: E (BHAK- 10379).

## Skeleton

Branching spicule tracts run mostly parallel to the sponge surface, but give rise to tracts that branch off to intersect the sponge surface. These flare to form bouquets that pierce the surface of the sponge. Some spicules are also confused throughout the choanosome and tangentially arranged within the ectosome (Figure 26C).

## Spicules

Nearly exclusively styles (Figure 26BD), straight or evenly curved, some subtylote but many simple; rare strongyles occasionally seen. Distributions of length were bimodal or trimodal for each sponge, but continuous with unclear size classes. When spicules from three samples are combined, they measure 118–285–446 x 1–8–12 μm, n=260 for length, n=253 for width. The combined length distribution is trimodal, with modes at approximately 210, 310, and 390 μm. Means per sponge were 259 to 301 μm in length and 7 to 8 μm in width.

## Distribution and habitat

Known from the Arctic, North Atlantic, and North Pacific, from 20–289 m depth (Dinn et al., 2020). The Southernmost Atlantic specimens are known from the Gulf of St. Lawrence (Dinn et al., 2020), while the Southernmost Pacific specimens are known from the San Juan Islands, Washington (de Laubenfels, 1961).

## Remarks

The choanosomal skeletons of halichondriid sponges are often described as "confused", but this term can also be applied to the taxonomy of many species in this group. This species is no exception. Populations of this species have been described as members of various families and genera; in the Northeast Pacific, these sponges were first described as *Stylissa stipitata* (de Laubenfels, 1961); it now seems likely that this name is a junior synonym of *Semisuberites cribrosa*. The samples we examined here agree with the recent morphological description of samples from the North Atlantic, though Atlantic samples had slightly larger spicules (Dinn et al., 2020).

The genus *Semisuberites* is monotypic and has been difficult to place. It was assigned to the Suberitidae by Topsent (1928); Van Soest and Hajdu (2002) noted similarities with *Hymeniacidon* but tentatively placed the genus in the Esperiopsidae (Order Poescilosclerida).

Van Soest (2016) later proposed that this genus is more likely to be in the order Suberitida. Here we present the first genetic data for this species, which clearly places it within the Suberitida. Based on morphological data and genetic similarity to some *Hymeniacidon* species, we propose placement in the Halichondriidae. The upright brushes of styles in the ectosome are unusual for this family, but there do appear to be some tangential spicules in the ectosome as well.

## Genus Topsentia Berg, 1899

Halichondriidae with a densely confused choanosomal skeleton and a crust-like tangential or paratangential arrangement of spicules in the ectosome. The ectosome is difficult to detach due to limited subdermal cavities. Spicules oxeas or modifications, in a large size range, including smaller spicules concentrated at the surface (paraphrased from Erpenback and Van Soest, 2002). Approximately 36 species are currently considered valid (de Voogd et al., 2024).

*Topsentia fernaldi* (Sim and Bakus, 1986)

Figure 27

Synonyms

Oxeostilon fernaldi Sim & Bakus, 1986

## Material examined

Holotype: USNM 33629, Bird Rock, Santa Catalina Island, California, (33.45000, -118.41667), 46 m, 1968-11-20. Other samples: TLT1134, Honeymooners, California, (36.50390, -121.94100), 10-20 m, 2021-09-22; TLT1261, Honeymooners, California, (36.50390, -121.94100, 10-20 m, 2021-09-22.

## Diagnosis

Thickly encrusting, irregular masses, yellow or chocolate brown exteriorly with yellow interiors. Sponges are hard and incompressible due to dense spiculation. The ectosomal skeleton is a crust of tangential spicules in a sieve-like pattern or a disordered mat. The choanosome is very dense with disordered spicules, with many oriented vertically near the surface, their tips piercing the ectosome. Spicules are oxeas in a large size range, 160–1170 μm long and 4–35 μm wide. Distribution of length is strongly bimodal, with modes near 250 and 850 μm. Spicules average 50% shorter in the ectosome, where most spicules are less than 600 μm long, but the full size-range is present in the choanosome.

## Morphology

The holotype was previously described as a thickly encrusting yellow mass, firm but friable, with a crusty surface texture and sparse oscula. Newly collected samples were thick, irregular encrustations, chocolate brown on the surface and pale yellow interiorly. Their interiors were full of bryozoan skeletons and worm tubes which are presumed to have been overgrown by the sponge, and the exterior was heavily fouled with hydroids, bryozoans, and other organisms.

Oscula were few, small, scattered, and flush with surface. Samples are white in ethanol.

## Skeleton

The original description states only that spicules are "distributed in an irregular fashion". We examined only a small portion of the holotype, and did not locate ectosomal regions. The choanosomal skeleton was confused, with a higher density of spicules than *Ha. panicea*, but a lower density than the recently collected samples from Central California.

Freshly collected samples from Central California had tangential ectosomal skeletons developed in some places as a sieve-like pattern around open spaces that were oval or polygonal, and in other places as a dense mat of discorded spicules. The choanosome was densely populated with spicules in confusion. Spicules became somewhat less confused towards the ectosome, where the majority are oriented vertically, with the final ones piercing the surface to create microscopic hispidity. No apparent spongin.

## Spicules

Nearly exclusively curved oxeas, with rare styles or strongyles sometimes present. Distributions of length were strongly bimodal, but continuous distributions made it difficult to define size classes. Spicules from the holotype measure 224–568–1129 x 7–15–31 μm, n=115; length modes at about 250 μm and 850 μm. When data from the holotype are combined with the two newly collected samples, spicules measure 168–511–1161 x 4–15–35 μm, n=499. See table 11 for data from each sample individually. When spicules are isolated from the ectosome and choanosome separately, the full range of lengths is seen in the choanosome, but ectosomal spicules are mostly from the shorter half of the length distribution (ectosome mean length 330 μm vs. choanosome mean length 614 μm).

**Table 11.** Spicule lengths of *Topsentia fernaldi*. Values given as min–mean–max in μm.

| Specimen | Ectosome | Choanosome |
| --- | --- | --- |
| TLT1134 | 177–406–880 n=45 | 218–648–1021 n=46 |
| TLT1261 | 168–282–1161 n=70 | 194–592–974 n=70 |
| *Topsentia* aff. *fibrosa*: |  |  |
| RBC10 | 288–463–977 n=95 | 300–667–1020 n=72 |

The holotype was originally described as having small oxeas, large oxeas, styles, and strongyles, with "styles and especially strongyles uncommon". Our spicule preparation from the holotype found only 2 styles, one tylostyle, and no strongyles, out of several hundred spicules, so we believe these to be rare aberrations (one style was also seen in another sample). We also do not consider oxeas to be present in size classes, because the length distribution is continuous, but the modes of the length distribution are very similar to the means of the size classes previously reported (239 and 813 μm).

## Distribution and habitat

Described from a single sample collected at Catalina Island, in Southern California, at 46 m. Two additional samples were recently collected in Carmel Bay in Central California, between 10-20 m. Both locations were natural rocky reefs.

## Remarks

The two sponges collected from Central California fit well into the current concept of *Topsentia*: they have a well-developed tangential ectosomal spicule crust that is difficult to remove, a densely confused choanosomal skeleton, and oxeas in a large size range, with smaller spicules at the sponge surface (Erpenbeck and van Soest, 2002). However, we have reservations in assigning them to *T. fernaldi.* The original description for this species is sparse, but it was described as "moderately dense, firm but friable", rather than stony and incompressible, as the newly collected samples are. The small fragment of the holotype we examined did appear to be less densely spiculated, consistent with this difference. However, the spicules from the new samples are very similar to the *T. fernaldi* holotype, and DNA extractions from the holotype were not successful. We therefore assign the new samples to this species, but would not be surprised that future collections yield new material that would result in splitting them.

We also examined a sponge from British Columbia, previously identified by William Austin as *Topsentia* aff. *fibrosa*. This sponge was morphologically indistinguishable from the Central California samples (table 11), but DNA extractions were again unsuccessful. It is therefore possible that *T. fernaldi* ranges farther North, but additional sampling of this genus in the Northeast Pacific is badly needed. *Topsentia fibrosa* is an Arctic species that may range into British Columbia, but has not been confirmed there. The original description of *T. fibrosa* is limited, but does state that samples are dark brown after preservation in spirit (Fristedt, 1887), while all the samples examined here were white after preservation in ethanol.

The presence of *T. fernaldi* within the Panicea species group is very surprising (Figure 1).

While much more distantly related species are morphologically indistinguishable from *Ha. panicea*, the spicule dimensions and density make this species quite distinct. A few other *Topsentia* species have genetic data available, and some appear to fall within entirely different orders of demosponges (Pankey et al., 2022; Redmond et al., 2013). These results serve to emphasize how evolutionary liable spicule size, shape, and density are in this clade.

Topsentia disparilis (Lambe, 1893)

## Synonyms

*Halichondria disparilis* Lambe, 1893

## Material examined

None.

## Description

First described from a single massive, flat sponge, 35 x 28 mm across and 11 mm thick. Spicules are exclusively or nearly exclusively oxeas, but in a very wide range of lengths, with two size-classes (or perhaps modes in a continuous distribution). Large oxeas originally described as 438–1287 μm long, averaging 13 μm thick (Lambe, 1893); a later report states they are 900 x 20 μm (de Laubenfels, 1953). Short oxeas reported as 130 x 8 μm (de Laubenfels, 1953) or 91 x 4 (Lambe, 1893). Choanosomal skeleton densely confused; ectosomal skeleton variously reported as a tangential reticulation of oxeas (Lambe, 1893) or bristling with erect spicules (de Laubenfels, 1953). The latter report of this species also notes it has scattered areas covered in low, small tubercles less than 1 mm high (de Laubenfels, 1953).

## Distribution and habitat

The holotype was collected at 73 m near Comox, Vancouver Island, British Columbia, and additional samples were reported from 38–45 m near Point Barrow, AK (de Laubenfels, 1953).

## Remarks

We did not examine any material from this species, but it is included here for completeness. Published descriptions do not clearly differentiate this species from *T. fibrosa* (Fristedt, 1887), which is also known from the Arctic, and may be a senior synonym. As stated in the *T. fernaldi* section above, further revision of *Topsentia* diversity in the Northeast Pacific is badly needed, but likely requires freshly collected material and additional DNA data.

Genus *Axinyssa* Lendenfeld, 1897

Halichondriidae lacking an ectosomal tangential skeleton. Choanosomal skeleton confused interiorly but with bundles near the periphery that protrude and form surface conules. (paraphrased from Erpenback and Van Soest, 2002). Approximately 28 species are currently considered valid (de Voogd et al., 2024).

Axinyssa piloerecta sp. nov.

Figure 28

Material examined

Holotype: TLT1007 (CASIZ 245235 & UCSB-IZC00069001) and additional sample TLT1132 (SBMNH 718614), both collected at Yellowbanks, Santa Cruz Island (33.99432, -119.51910), 24-30 m, 2021-09-15.

## Etymology

From the word piloerection, the process that causes skin to form "goosebumps" and hair to stand up, creating a pattern similar to the surface of this sponge.

## Diagnosis

Yellow cushion-shaped sponges with widely-spaced surface conules. The combination of a smooth, slick-looking surface and conules with protruding spicule bundles is distinctive in the region. This contrasts with the sympatric *Obruta collector* Turner & Lonhart 2024, which has similar spicules but lacks raised bumps and is hispid over the entire surface. The large oxeas (530–1480 x 5–33 μm) and the significant fraction of spicules that are styles (18%) further distinguish this species from others in the region.

## Morphology

Cushion-shaped irregular mounds, pale yellow when alive and beige in ethanol. The surface is covered in raised projections, 1–3 mm apart, 1–3 mm in height, with spicules projecting an additional 0.5 mm. When alive, the holotype was approximately 60 x 35 mm across, 24 mm in maximum thickness; sampled portion approximately 35 x 35 x 24 mm. A distinct surface membrane covers the sponge, creating a smooth and slick-looking surface except where spicules protrude from raised conules.

## Skeleton

A dense, confused mass of spicules internally, with no apparent organization. Spicules become roughly parallel towards the periphery, pointing outwards, and form vague plumose tracts.

Some of these tracts end in tufts of spicules that pierce the surface of the sponge. No ectosomal skeleton. Spongin not apparent, but body of sponge is collagenous and dissolves very slowly in bleach, in contrast to all other Halichondridae described here, which dissolved very rapidly.

## Spicules

Styles and oxeas, with some intermediates; one strongyle also present. Oxeas are curved or bent, usually near center, sometimes with weak centrotylote swelling (Figure 28F). Dimensions highly variable, but without clear size categories; 533–957–1476 x 5–24–33 μm (n=78). Styles are less common than oxeas (18% of spicules), but with similar dimensions. 606–1178–2166 x 5–17–27 μm (n=17).

**Figure 27.**
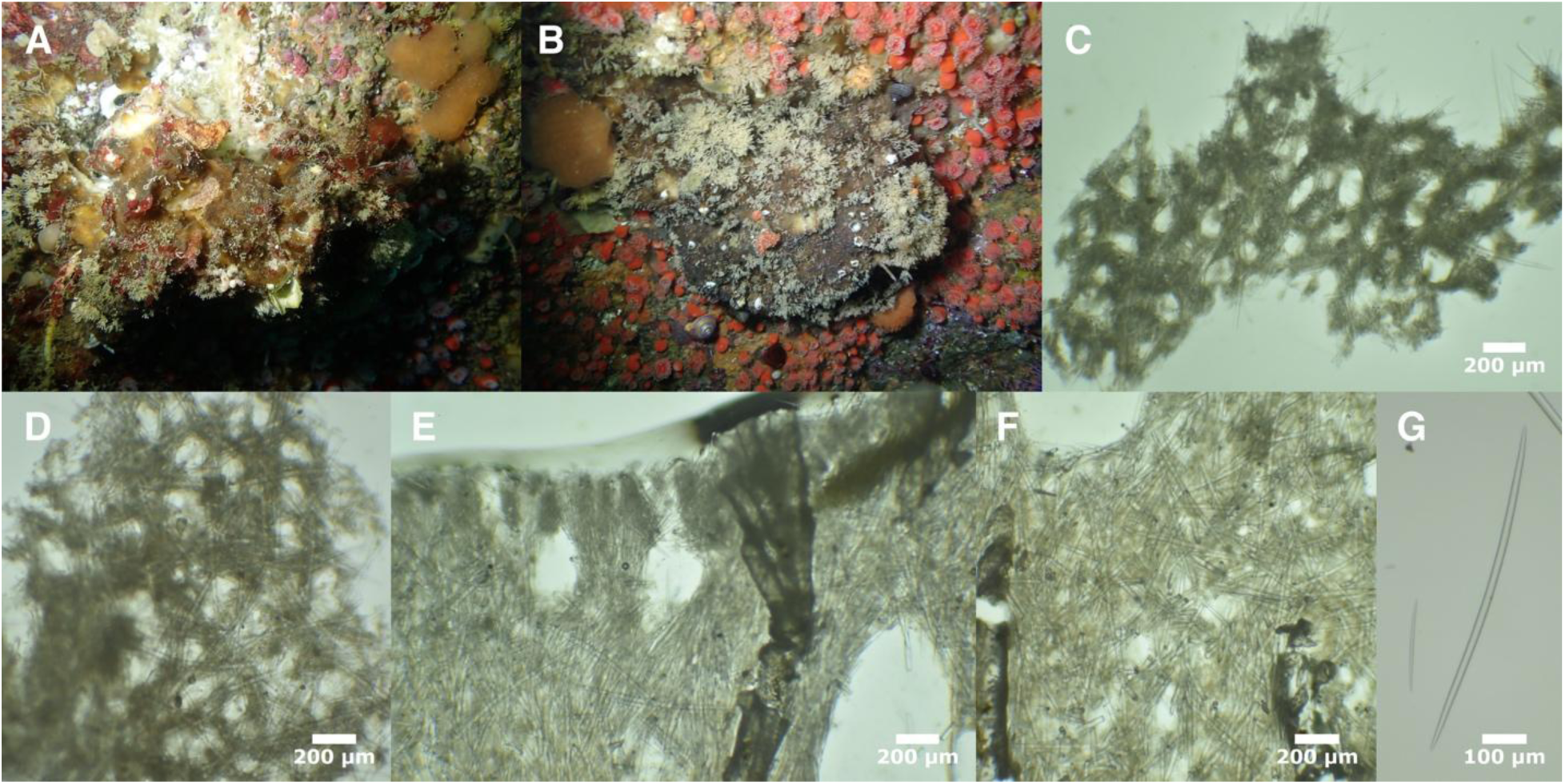
*Topsentia fernaldi*. A (TLT1134), B (TLT1261). C & D: tangential ectosomal skeleton (TLT1261). Cross-sections show choanosomal skeleton at sponge surface (E) and farther towards the interior (F), TLT1261. G: long and short oxeas from holotype, USNM 33629.

**Figure 28.**
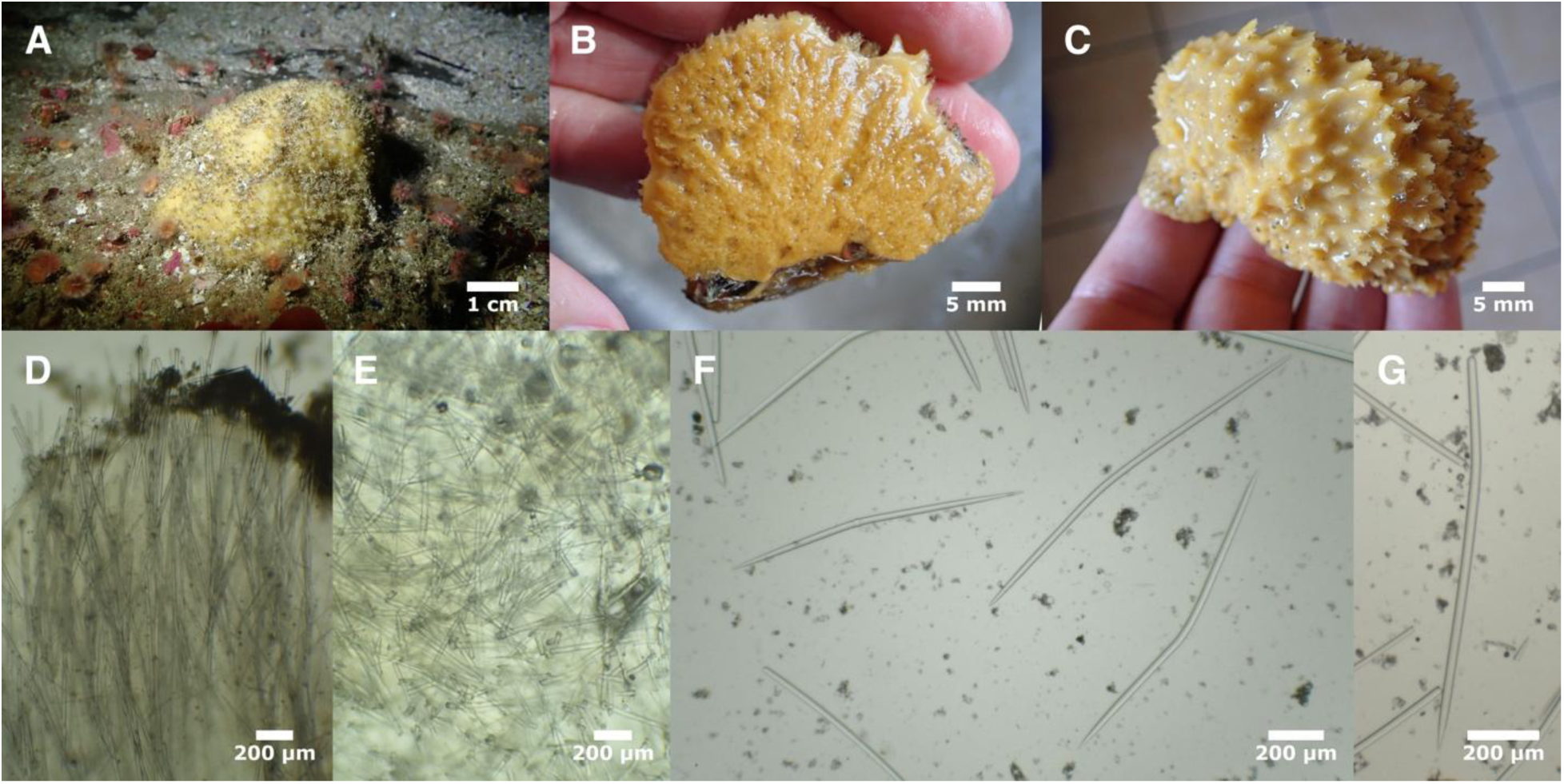
*Axinyssa piloerecta*. Sample shown in situ (A) and cross-sectioned after collection, before preservation (B). C: sample TLT1132 after collection, before preservation. Cross-sections show choanosomal skeleton at sponge surface (D) and farther towards the interior (E). F: example oxeas, G: example style. All photos from TLT1007 except C.

## Distribution and habitat

Found at only a single location: a rocky reef near Yellowbanks, Santa Cruz Island. The site is a rocky outcrop, typical of other rocky reefs in the region, but most other sites searched in the Northern Channel Islands have been in less than 20 m of water, while this site was 24-30 m in depth. The species appeared to be common at the site at the time of collection.

## Remarks

The morphology of this species places it firmly within the *Axinyssa* (Erpenbeck and van Soest, 2002). Like the genus *Topsentia*, the genus *Axinyssa* is highly polyphyletic, with some species, including the type species, falling among the order Bubarida and other species falling in the order Biemnida (Morrow and Cárdenas, 2015; Morrow et al., 2013; Pankey et al., 2022; Thacker et al., 2013). Genetic data from this species was previously reported as placing it clearly in the order Axinellida (Turner and Pankey, 2023). We describe it here because in the current taxonomy, this genus remains within the Halichondriidae, but we expect future taxonomic revision to move this species to the Axinellida.

Two species of *Axinyssa* are known from the North Pacific: *A. tuscara* (Ristau, 1978) and *A. isabela* (Carballo and Cruz-Barraza, 2008). The new species is distinguished from these species by its long spicules and high fraction of spicules that are styles. These same features distinguish it from species known from the temperate Atlantic.

Axinyssa tuscara (Ristau, 1978)

## Synonyms

*Axinomimus tuscaru*s Ristau, 1978

## Material examined

Holotype: USNM 24527, Carmel River Beach Point, Carmel, California, 7 m, 1973-04-10.

## Diagnosis

Thickly encrusting dark brown sponges that remain brown after preservation in ethanol. Soft and compressible. Choanosomal skeleton of vertical spicule tracts that pierce sponge and project up to 1.5 mm beyond sponge surface, connected by a confused reticulation of individual spicules. No special ectosomal skeleton. Spicules oxeas only, longer and considerably thicker than sympatric *Halichondria*, 355–552 x 10–22 μm.

## Morphology

Previously described as dark brown exteriorly and dark chocolate brown interiorly; thickly encrusting and amorphous; soft and compressable. The small piece of the holotype we examined was consistent with this description, though we were unable to compare ectosome and choanosome.

## Skeleton

Previously described as having vauge and somewhat plumose spicule tracts, with spicules and spicule bundles protruding through the ectosome. Lacking a special ectosomal skeleton. In the small piece we examined, and could only determine that the chromosomal skeleton was confused.

## Spicules

Curved oxeas, 355–476–552 x 10–18–22 μm, n=47.

## Distribution and habitat

Collected in situ only from the type location: Carmel River Beach Point, Central California, on a rock reef at 7 m depth. It was described as rare at this location (Ristau, 1978). Several samples were also collected in beach wrack in Northern California at Bodega Head.

## Remarks

All known samples of this species were collected from 1973 to 1977 and were included in the original description (Ristau, 1978). We examined a small piece of the holotype in order to attempt DNA extraction, which failed. We also performed a spicule preparation in order to publish more detailed information on the distribution of spicule sizes, as described above. It was difficult to characterize the skeleton in the small piece we examined, and could only determine that the chromosomal skeleton was confused.

## Conclusions

Here, we have documented a remarkable diversity of sympatric sponges in the family Halichondriidae. We combined extensive field collections, genomic analysis, and traditional morphological characters to reject both the "single globally-distributed invasive species" and "cryptic endemic species in different regions" hypotheses. We reject the single-species hypothesis by using multi-locus genomic data to infer reproductive isolation between sympatric populations. When molecular phylogenies are built separately for mitochondrial genomes and the nuclear-encoded ribosomal locus, we find numerous clades with reciprocal monophyly in both compartments. Because the mitochondrial and nuclear genomes are unlinked, reciprocal monophyly across these compartments is strong evidence of reproductive isolation when species are sympatric (Jennings, 1917; Rannala and Yang, 2020). We are also able to reject the "cryptic endemic species in different regions" hypothesis for many (but not all) of the delineated species because we find a lack of correspondence between geographic region and species range. Instead, we find numerous species that have likely been dispersed by human activity.

The discovery of cryptic species in this group is not entirely surprising, given the few known morphological traits that distinguish them. It is also not surprising that some species have been spread through accidental human introductions, particularly since they are common on floating docks and other areas where introduced species have been observed, and other species in the family are known to be introduced (Carlton and Eldredge, 2009; Turner, 2020).

However, we did not expect to find so many cryptic introduced species, sympatrically distributed at even small scales. *Halichondria bowerbanki*, *Ha. zabra* sp. nov., *Ha. urca* sp. nov., *Ha. pinaza* sp. nov., and *Ha. hygeia* sp. nov. were all found growing on the same floating docks in a small harbor in Santa Barbara, California, and we document at least 4 of these species in Eastern North America and/or Europe. We hope that these unintentional "common gardens" of related sponges will facilitate future studies of their evolution and ecology.

We were unable to find morphological traits to diagnose many of these species. This presents challenges for ecological monitoring projects that lack genotyping capabilities.

However, accurately delineating reproductively isolated groups is essential for monitoring both native diversity and potentially invasive species. Additionally, biologists studying physiology, biochemistry, or development need to know which species they are examining so that others can build on their findings. Despite their morphological similarities, we suspect that many of these species will prove to be biologically distinct. For example, *Halichondria panicea* and *Ha. bowerbanki* are estimated to share a common ancestor in the mid-Cretaceous period, nearly 100 million years ago (Pankey et al., 2022). Even more dramatically, *Halichondria panicea* and *Ha. galea* sp. nov. have been independently evolving since the Jurassic (Pankey et al., 2022), splitting around the same time as the common ancestor of all Eutherian mammals (Archibald, 2003; Luo et al., 2011). While the Eutherian clade was diversifying into bats, whales, and primates, what were these sponges up to? We hope that our results will motivate future work on this question.

## Acknowledgements

We are grateful for the help and support of many people in UCSB’s Marine Science Institute and Diving & Boating Program, especially Robert Miller, Clint Nelson, Christoph Pierre, and Christian Orsini. Numerous NOAA personnel provided logistics and planning support for collections in the Monterey Bay and Olympic Coast National Marine Sanctuaries. Raquel Pereira graciously shared samples from Sweden and Portugal, and Brooke Weigel shared samples from Tatoosh, Washington. Gustav Paulay, Hugh MacIntosh, Abigail Reft and Christina Piotrowski provided access to their collections at the Florida Museum of Natural History, Royal British Columbia Museum, Smithsonian National Museum of Natural History, California Academy of Sciences, respectively. We also thank Matt Whalen, Margot Hessing-Lewis, and the staff, students and volunteers of the Hakai Institute for support during the 2017 Hakai-MarineGeo bioblitz, and the Makah Tribal Nation for supporting the study of sponges from Tatoosh Island. We thank DFO colleagues Andrew Cooper, Peter Lawton, Laura Teed and Torben Brydges from Saint Andrews Biological Station, who partnered on the Eastern Shores habitat survey project and participated in specimen collection.

## Supporting Information

Data available from the Dryad Digital Repository (https://doi.org/10.5061/dryad.bvq83bkk6).

